# Ancient admixture catalyzes homoploid hybrid speciation and intense genomic erosion in an Asian langur genus

**DOI:** 10.64898/2026.06.17.732823

**Authors:** Jiwei Qi, Qian Zhao, Liye Zhang, Dilina Rusitanmu, Ying Shen, Gaoming Liu, Xiang Li, Yang Teng, Roberto Portela Miguez, Nguyen Van Truong, Minh D. Le, Tilo Nadler, Ralph Schönfelder, Xuming Zhou, Zhijin Liu, Christian Roos, Ming Li

**Author notes:** Corresponding author:* Prof. Ming Li, Key Laboratory of Animal Ecology and Conservation Biology, Institute of Zoology, Chinese Academy of Sciences, 1-5 Beichenxi Road, Chaoyang District, Beijing 100101, China. Prof. Christian Roos, Gene Bank of Primates, Primate Genetics Laboratory, German Primate Center, Leibniz Institute for Primate Research, 37077 Göttingen, Germany. Prof. Zhijin Liu, College of Life Sciences, Capital Normal University, 105 Xisanhuanbei Road, Haidian District, Beijing 100048, China. Contributed equally to this work (Jiwei Qi, Qian Zhao, Liye Zhang).

## Abstract

Ancient admixture catalyzes evolutionary innovation, yet its long-term genomic consequences for newly formed lineages remain poorly understood. Here, based on 53 genomes covering 19 species of the Asian langur genus *Trachypithecus*, we explore admixture’s role in shaping a reticulated radiation. Genome-wide analyses of phylogenomic triplet topologies demonstrate that phylogenetic discordance across this radiation is primarily driven by widespread introgression rather than incomplete lineage sorting. To mitigate historical noise and resolve the ancestral species tree, we anchored our phylogenetic analysis on the X-linked recombination desert (XLRD), which exhibits an 84.5% introgression reduction compared to autosomes. We identify Delacour’s langur as a clear case of homoploid hybrid speciation, arising from ancient admixture with ∼70:30 genomic contributions from ancestral northern and southern limestone langur lineages. This hybrid species fixed key reproductive isolation loci; notably, alternate inheritance of pigmentation genes (*SLC45A4*, *HPS5*, *ADCY10*) systematically coupled with a fixed *RNF175* chimeric allele to drive its diagnostic pelage phenotype. This illustrates how introgressed multi-gene networks rapidly establish prezygotic visual barriers and profound phenotypic divergence. However, subsequent spatial isolation within fragmented karst landscapes forced a major conversion of genetic burden into realized load. Genome-wide, over 82% of loss-of-function variants occur in a homozygous state, reflecting the expression of lethal recessive mutations within long runs of homozygosity. Together, our findings demonstrate that ancient admixture can trigger homoploid hybrid speciation, yet subsequent ecological restriction locks derived lineages into severe, long-term genomic erosion, revealing a fundamental trade-off in reticulate evolutionary radiations.

## Introduction

Ancient admixture is now recognized as a pervasive evolutionary force that can accelerate diversification by introducing novel genetic variation and facilitating rapid adaptation^1,2^. However, quantifying its role in shaping genome architecture and long-term evolutionary trajectories remains challenging, particularly because ancient admixture often coincides with rapid species radiations^3,4^. In such radiations, evolutionary history rarely follows a simple bifurcating pattern. Instead, diversification proceeds through reticulate evolution, in which lineage splitting is repeatedly intersected by gene flow among partially diverged taxa^5–7^. Under these conditions, phylogenetic signal is shaped jointly by incomplete lineage sorting (ILS), the stochastic retention of ancestral polymorphism and ancient introgressive hybridization, producing discordant genealogies across the genome and obscuring the true sequence of speciation events^3,5,6^. Importantly, admixture in such systems is not merely a source of analytical noise; it can fundamentally reorganize genomes and redirect evolutionary trajectories. A highly creative yet structurally complex outcome of ancient admixture is homoploid hybrid speciation (HHS)^2,8–10^, in which hybridization gives rise to a reproductively isolated lineage without a change in chromosome number^9^. Although theoretical and empirical studies suggest that hybridization can facilitate adaptive novelty and ecological expansion, clear genomic demonstrations of HHS—especially in animals—remain rare.

Moreover, existing work has largely focused on the short-term creative potential of hybridization, leaving its long-term genomic consequences insufficiently explored^11,12^, particularly regarding how initial adaptive gains withstand the systemic mutational costs imposed by subsequent ecological specialization and confinement^13,14^. This imbalance gives rise to a key evolutionary paradox. Although ancestral admixture is widely viewed as a driver of adaptive novelty, it may also impose lasting genomic costs. Following hybrid origin, demographic contraction, ecological specialization, and the accumulation of genetic incompatibilities may promote genome-wide loss of ancestral variation and elevated mutational load^13,15^. Whether hybrid lineages retain their initial adaptive advantages or instead experience progressive genomic erosion remains a fundamental unresolved question.

Asian langurs of the genus *Trachypithecus* provide an exceptional system for investigating these processes. Monkeys of this genus are distributed across a complex biogeographic landscape spanning mainland Southeast Asia and Sundaland^16–23^, a region characterized by repeated geological fragmentation, climatic oscillations, and secondary contact. Consequently, langurs exhibit remarkable phenotypic, behavioral and ecological diversity^18,24–28^. They comprise 22 species, traditionally assigned to four species groups: the *pileatus*, *francoisi*, *cristatus*, and *obscurus* groups^22^ (Fig. 1A). Yet this apparent taxonomic structure masks a long history of phylogenetic instability, suggesting that reticulate evolutionary processes have played a prominent role in shaping the radiation. Indeed, despite decades of study, the evolutionary relationships within *Trachypithecus* remain highly contentious. Morphology-based classifications and early molecular analyses have generated profoundly conflicting topologies^16,17,22,29^. Recently, investigations within specific sibling lineages of this radiation have proposed that a baseline of stochastic lineage sorting, with ILS accounting for 8.9% of whole-genome segments, can maintain ancestral polymorphisms to drive local phenotypic mixing^11^. However, whether this restricted baseline of ILS is sufficient to explain the diversification of *Trachypithecus* langurs remains highly questionable. It remains unresolved whether the genus-wide phylogenetic discordance is merely an artifact of deep coalescent variance, or if pervasive interspecific gene flow has fundamentally reorganized their genome-wide architecture^24,30^.

**Fig. 1.**
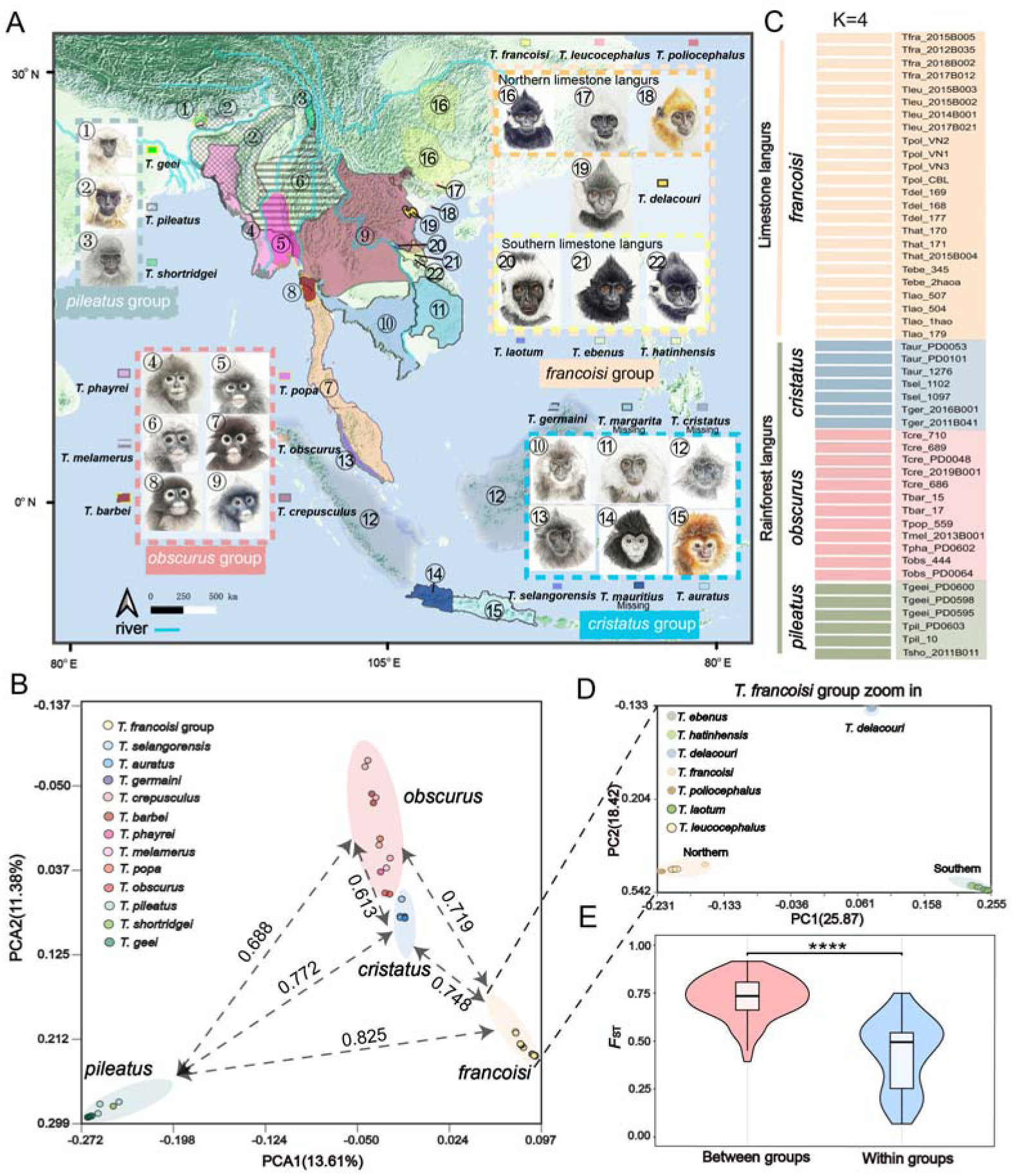
Geographic distribution and population genetic structure of the genus *Trachypithecus*. **A** Geographic distribution of all *Trachypithecus* species across Southeast Asia. The ranges of the four species groups (*pileatus*, *obscurus*, *cristatus*, and *francoisi*) are delineated by different colors. **B** Principal component analysis (PCA) of all sequenced *Trachypithecus* individuals. Dashed lines and associated values indicate the average pairwise *F*_ST_ between the four species groups. **C** Population genetic structure inferred based on Admixture at K = 4, perfectly clustering the individuals into the four recognized species groups. **D** A zoomed-in PCA plot exclusively focusing on the *T. francoisi* group. **E** Comparison of genome-wide *F*_ST_ values between and within species groups. Significance was calculated using a two-sided Wilcoxon rank-sum test (\*\*\*\**P* < 0.0001). Facial portraits of the 22 recognized *Trachypithecus* species were generated using Gemini AI.

Beyond resolving phylogenies, deciphering the genomic legacy of *Trachypithecus* langurs is critical for their survival. Driven by historical isolation and severe anthropogenic pressures, over 70% of these primates are now classified as Endangered or Critically Endangered^31^. For these highly threatened species, evaluating extinction risk requires a direct understanding of how geographic isolation shapes their genetic health^12,13,32^. This assessment is especially urgent for the limestone langurs strictly restricted to fragmented karst landscapes. Within these narrow, island-like habitats, prolonged isolation and genetic drift act as a severe genomic crucible, forcing historically masked deleterious mutations into high realized genetic load^15,33^. Determining the genetic consequences of this intense restriction is therefore essential.

In this study, we compiled a genomic dataset of 53 whole genomes representing 19 species, capturing most of the taxonomic and geographic diversity of the genus *Trachypithecus* (Table S1). Using this dataset, we first resolved the complex reticulate evolutionary relationships across the radiation, and then investigated how the initial adaptive benefits arising from hybrid origins can impose a long-term evolutionary debt when followed by geographic restriction. This framework allowed us to assess the ultimate genomic consequences of reticulate speciation, particularly under the crucible of karst isolation for specialized lineages. Our results demonstrate that ancient hybridization and subsequent genomic reorganization were not peripheral complications but central forces shaping the reticulate radiation of *Trachypithecus*. Crucially, by connecting these adaptive origins to the subsequent accumulation of realized mutational burden, this study exposes the profound dual nature of reticulate evolution—illuminating both the rapid ecological breakthroughs and the long-term genomic fragilities that dictate the survival of endangered, highly specialized primates.

## Results

### Sample authentication, genome sequencing and quality control of museum specimens

To achieve genus-level genomic coverage for *Trachypithecus*, we newly sequenced whole genomes from 12 individuals representing four species. Among these, genomic data for *T. barbei*, *T. popa*, and some *T. crepusculus* individuals were obtained from museum specimens. DNA authenticity was verified via deamination patterns and endogenous content^34^. The majority of museum-derived samples exhibited characteristic deamination signatures and fragment properties consistent with historical DNA, supporting their authenticity (Fig. S1). The newly sequenced genomes were subsequently integrated with 41 previously published *Trachypithecus* genomes. All sequencing reads were processed using a unified pipeline and aligned to the golden snub-nosed monkey reference genome (*Rhinopithecus roxellana*; GCA_007565055.1) to ensure consistency across datasets and facilitate comparative genomic analyses. Using *R. roxellana* and *R. bieti* as outgroups, we constructed a comprehensive dataset of 55 individuals for phylogenetic analysis (Fig. 1A and Table S1). Although we were unable to collect genomic data for three species (*T. margarita*, *T. cristatus*, and *T. mauritius*), our dataset encompasses 19 of the 22 currently recognized species of *Trachypithecus*.

After quality control, we identified 156,262,744 high-quality single-nucleotide variants (SNVs) across the dataset (Fig. S2 and Table S2). Kinship assessment revealed no individuals are related at the second-degree or higher (Fig. S3), allowing all 53 *Trachypithecus* individuals to be retained for phylogenetic reconstruction. For analyses particularly sensitive to contamination, including tests of introgression, we applied more stringent authentication criteria. Four historical samples (Tcre_598, Tcre_599, Tcre_600, Tpop_674) that lacked characteristic deamination signatures were conservatively excluded from gene flow analyses. This additional filtering resulted in a high-confidence dataset of 49 *Trachypithecus* genomes representing 19 species. The retained museum samples exhibited high mapping rates (78.69–97.44%) and moderate to substantial sequencing depth (14.19–25.14×), indicating that, despite their historical origin, these samples provide reliable genomic information for downstream evolutionary analyses. Together, these authentication and quality control steps ensured a robust and conservatively curated dataset suitable for phylogenetic and population genomic inference.

### X-Linked recombination-depleted regions resolve phylogenetic discordance among species groups

Whole-genome principal component analysis (PCA) and ADMIXTURE profiling consistently identified four distinct genetic clusters corresponding to the four species groups. The first principal component (PC1; 13.61%) primarily captured ecological divergence between the karst-adapted (i.e., *francoisi* group) and rainforest-adapted lineages (i.e., *pileatus*, *obscurus* and *cristatus* groups) (Figs. 1B, S4 and S5). At K=4, the four species groups were distinctly separated (Figs. 1C and S4). Within the *francoisi* group, substructure analysis revealed a clear latitudinal gradient along PC1 (25.87%), differentiating northern from southern populations (Fig. 1D), a pattern supported by ADMIXTURE (Figs. S6 and S7). Consistent with these patterns, pairwise fixation indices (*F*_ST_) showed high values between representative lineages (0.635–0.825), which were significantly higher than within-group divergence (Wilcoxon test *P* < 0.001; Fig. 1E). Together, these results firmly support clear genetic boundaries consistent with the four-group taxonomic classification^22^ (Fig. 1B; Table S3).

Phylogenetic reconstruction based on concatenated autosomal SNVs using Maximum Likelihood (ML) revealed the *pileatus* group as the earliest-diverging lineage and the *francoisi* group as sister clade to the common ancestor of the *obscurus* and *cristatus* groups (T1) (Figs. 2A, 2B and S8). Similarly, the ML tree constructed from whole mitochondrial genomes broadly supported the T1 topology, except that the Indochinese grey langur (*T. crepusculus*) of the *obscurus* group clustered distantly with the *francoisi* clade (Fig. S9). In contrast, X-linked SNVs supported an alternative topology (T2), in which the *pileatus* group remains basal, but the *obscurus* group was resolved as the sister lineage to the common ancestor of the *francoisi* and *cristatus* groups (Figs. 2B, S10 and S11).

**Fig. 2.**
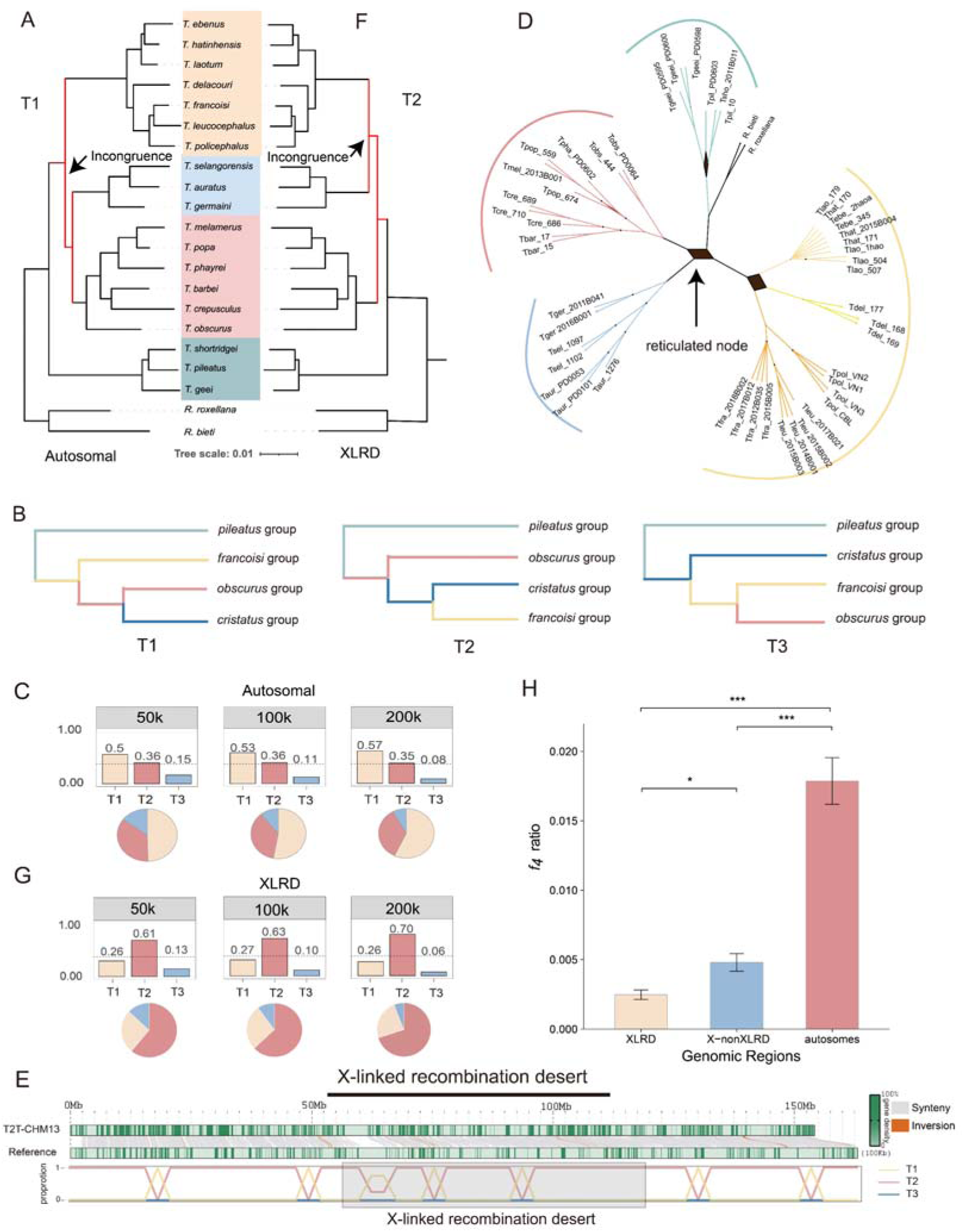
Resolution of pervasive phylogenetic incongruence within *Trachypithecus* using the X-linked recombination desert (XLRD). **A** Maximum Likelihood (ML) phylogeny reconstructed using concatenated genome-wide autosomal SNVs (T1). Species groups are highlighted with different colors. **B** Schematic representation of the three competing phylogenetic hypotheses (T1, T2, and T3) regarding the branching order of the *francoisi*, *obscurus*, and *cristatus* groups. **C** Frequencies of the three competing topologies (T1, T2, and T3) inferred from local autosomal gene trees using non-overlapping sliding windows of 50 kb, 100 kb, and 200 kb. Bar plots show the proportion of windows supporting each topology, with corresponding pie charts below. **D** SplitTree phylogenetic network based on autosomal SNVs, revealing distinct reticulated nodes (black arrows) indicative of ancient genetic admixture or complex evolutionary scenarios at the base of major clades. **E** Synteny analysis between the *Rhinopithecus roxellana* reference genome and the human T2T-CHM13 genome, delineating the evolutionary conserved X-linked recombination desert (XLRD, shaded grey box). The bottom track displays the rapidly fluctuating topology weights across the X chromosome. **F** ML phylogeny reconstructed using exclusively SNVs from the identified XLRD region (T2), which robustly resolves the *obscurus* group as the sister lineage to the *francoisi*-*cristatus* clade. **G** Frequencies of the three competing topologies across the entire X chromosome using local gene trees at 50 kb, 100 kb, and 200 kb window sizes. In stark contrast to the autosomal data, the X chromosome predominantly supports Topology 2 (T2). **H** Comparison of introgression levels, measured by the *f*4 ratio, across the XLRD and non-XLRD regions on the X chromosome, and autosomes. The XLRD exhibits a profound reduction in introgression compared to both autosomal regions (Wilcoxon rank-sum test, *** *P* < 0.001 and * *P* < 0.05), quantitatively demonstrating its robust resistance to interspecific gene flow.

Genomic heterogeneity across the dataset was assessed by conducting ASTRAL analyses^35^ using non-overlapping sliding windows of 50 kb (n=13,974), 100 kb (n=27,928), and 200 kb (n=13,960), which revealed widespread topological discordance across the genome (Fig. 2C; Fig. S12). At the focal node resolving the *obscurus*, *francoisi*, and *cristatus* trichotomy, T1 received 50% to 57% quartet support, whereas T2 also commanded substantial support (35–36%; PP=1) (Fig. 2C, Fig. S12 and S13). Comparable levels of discordance were observed at other phylogenetically challenging nodes, including the one within the *pileatus* group (Tp1=48%, Tp2=37%, Tp3=15%), within the southern limestone langur clade (Ts1=41%, Ts2=31%, Ts3=28%), and regarding the placement of *T. delacouri* (Tf1=40%, Tf2=33%, Tf3=27%) (Fig. S12). This pervasive incongruence was further visualized by Twisst^36^, which revealed a genome-wide mosaic distribution of topologies indicative of a rapid radiation characterized by pervasive ILS (Fig. S14). Consistently, PhyBin analysis further characterized the diversity and frequency of competing phylogenetic signals by identifying 138 distinct topologies (Fig. S15), and DensiTree displayed an extensive overlap and branching uncertainty among gene trees (Fig. S16), reflecting high phylogenetic uncertainty. SplitsTree networks^37^ further revealed distinct reticulate structures at central nodes, suggesting that ancient gene flow may have contributed to the observed discordance in addition to ILS (Fig. 2D and Fig. S17).

To reduce the confounding effects of introgression, we focused our reconstruction on the X-linked recombination desert (XLRD), a genomic region resistant to introgression and enriched for ancestral signals^38^. We extracted SNVs from the reference region syntenic to the human T2T-CHM13 genome and reconstructed the phylogeny, which yielded a well-resolved species tree supporting the T2 topology (Fig. 2E and 2F). Sliding-window analyses across the entire X chromosome further confirmed this pattern, with 61%, 63%, and 70% of windows at 50 kb, 100 kb, and 200 kb scales, respectively, recovering T2 as the predominant topology (Fig. 2E and 2G). To quantify this resistance to gene flow, we compared introgression levels (measured by *f*4-ratio) across the XLRD and non-XLRD regions of the X chromosome, and autosomes. This analysis revealed that introgression within the XLRD was significantly reduced by 84.5% compared to autosomal regions (Wilcoxon rank-sum test, *P* < 0.001) and by 40.1% relative to non-XLRD regions on the same chromosome (Wilcoxon rank-sum test, *P* < 0.05) (Fig. 2H). Together, these results indicate that T2 most parsimoniously represents the underlying species tree for the genus.

Spatiotemporally, our dating and dispersal models reconstruct a dynamic southeastward radiation for *Trachypithecus* (Figs. 3C and S18). Following an initial divergence from *Semnopithecus* near northeastern India and Bhutan (∼4.87 million years ago [Mya]), the basal *pileatus* group separated from the remaining lineages (∼3.96 Mya). The genus then underwent an explosive radiation (∼3.27 Mya): the *obscurus* group dispersed south toward the Malay Peninsula, while the *cristatus* group originated (∼3.03 Mya) and later capitalized on cyclical Early Pleistocene sea-level regressions (∼1.7–1.3 Mya) that exposed the vast Sunda Shelf, facilitating a dramatic southward colonization of Sumatra, Borneo, and Java^39–41^. Finally, initiating its own radiation (∼1.36 Mya), the *francoisi* group expanded eastward into fragmented karst landscapes. Demographically, the pairwise sequentially Markovian coalescent (PSMC) analyses^42^ revealed that around the Plio-Pleistocene transition (2-3 Mya), all four groups experienced a synchronous massive demographic expansion (Ne ∼ 10L), providing the crucial foundation for the observed explosive diversification and pervasive ILS (Fig. S19). Consistent with these inferences, species distribution models (SDMs)^43^ revealed climate-driven range dynamics characterized by repeated cycles of expansion and contraction, with karst-associated glacial refugia playing a particularly important role in shaping the distribution of the *francoisi* group (Fig. S20 -S23).

**Fig. 3.**
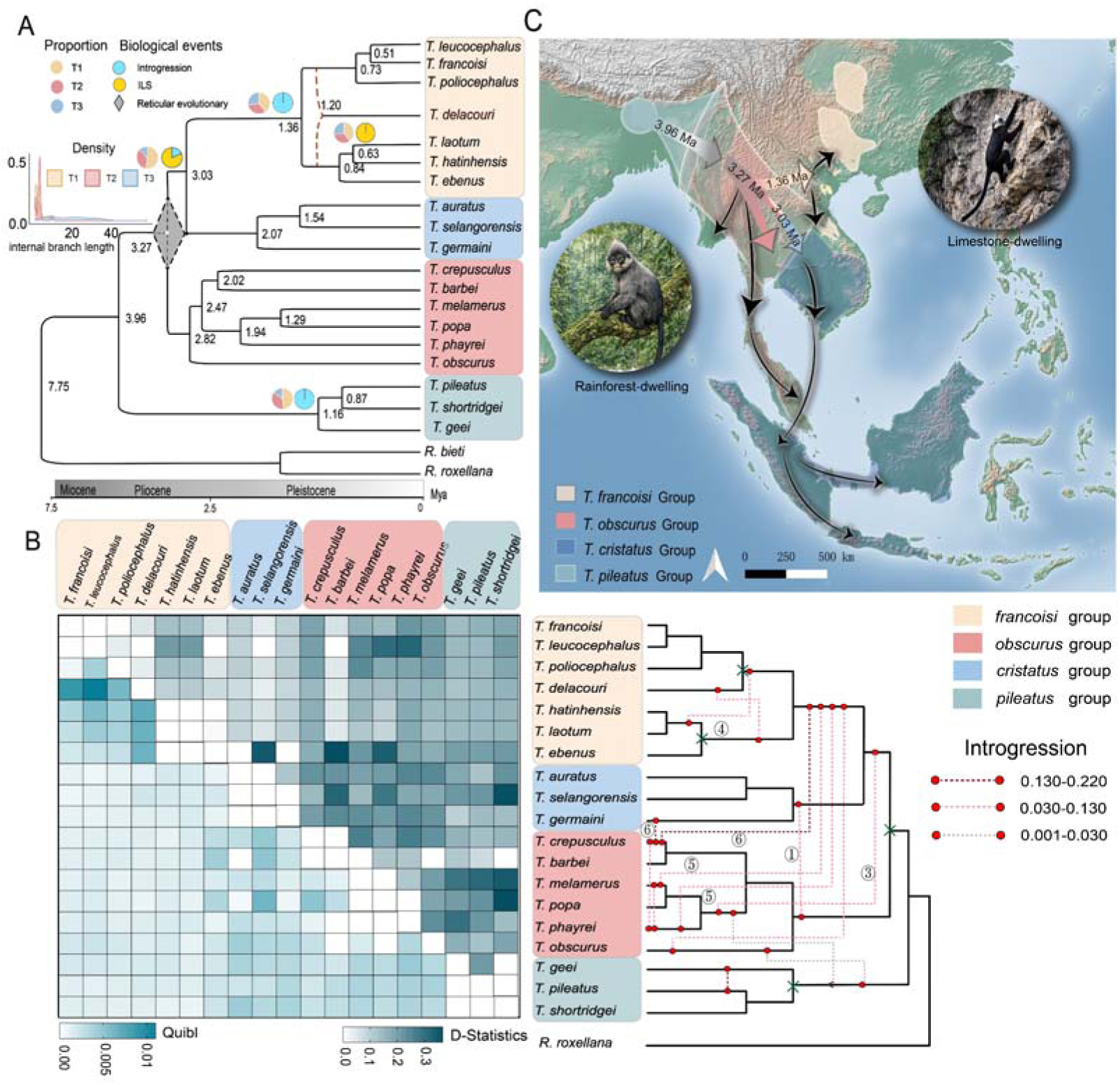
Incomplete lineage sorting (ILS), complex reticulate evolution, and spatiotemporal dispersal across the genus *Trachypithecus*. A Time-calibrated species tree of *Trachypithecus*. Pie charts at key ancestral nodes denote the genome-wide proportions of the three competing topologies (T1, T2, and T3). Adjacent symbols indicate the primary microevolutionary mechanisms driving local phylogenetic discordance: ILS (yellow circles) or introgressive hybridization (blue circles), as partitioned by QuIBL. The grey dashed diamond highlights a major macroevolutionary reticulated node anchoring the base of the *obscurus*, *cristatus*, and *francoisi* groups. The inset density plot displays the distribution of internal branch lengths for the three alternative topologies, exhibiting a pronounced right-shifted peak for T2 (the parsimonious underlying species tree) and a sharp left-shifted peak for T1, a classical signature of rapid evolutionary radiation fueled by ancestral variance. **B** Left: A pairwise genetic admixture heatmap quantifying multi-layered reticulation, with the lower -left triangle showing QuIBL admixture proportions and the upper-right triangle tracking *D*-statistics. Right: A reconstructed phylogenetic network resolving comprehensive introgression pathways across the radiation. Red dotted lines indicate the direction and genetic magnitude of localized gene flow, with circled numbers (1 to 6) explicitly mapping the six massive, multi-directional introgression events identified by the *f*branch matrix. **C** Spatiotemporal dispersal model illustrating the progressive east and southward radiation of the genus. Arrows depict the hypothesized ancestral migration routes and estimated divergence timelines (Mya) of the four species groups across the dynamic biogeographic landscape of Southeast Asia and the Sunda Shelf.

### Heterogeneous contributions of incomplete lineage sorting and introgression to phylogenetic discordance

To disentangle the relative contributions of incomplete lineage sorting (ILS) and introgression to the observed phylogenetic discordance, we first utilized QuIBL (Quantifying Introgression via Branch Lengths)^3^ to evaluate internal branch lengths (C2) across alternative topologies. Analyses of internal branch supported T2 as the primary species tree, exhibiting the highest genomic density (0.5) with a right-shifted peak indicative of a persistent phylogenetic signal (Fig. 3A). Conversely, the discordant T1 topology displayed a substantial left-shifted peak, suggesting rapid radiation and short internal branches predominantly driven by ILS (Fig. 3A). Extending these analyses to all 126 triplets among the *obscurus*, *francoisi*, and *cristatus* groups revealed widespread phylogenetic complexity. T1 displayed a markedly higher proportion of loci where discordance was predominantly attributable to ILS (77.8%) compared to T2 (40.5%), although both topologies exhibited evidence of introgression (23 and 30 triplets supporting an “ILS + Introgression” model, respectively; Fig. S24, Tables S4 and S5). Furthermore, T3 topology yielded significant signals for the “ILS + Introgression” model (ΔBIC < −10; Tables S4 and S5), indicating a history of recurrent, temporally distinct admixture episodes (internal branch ranging from 7.21 to 43.96). Further examination of other recalcitrant nodes highlighted heterogeneity in the processes shaping genome-wide discordance. The southern limestone langur node (i.e., *T. laotum* -*T. ebenus* -*T. hatinhensis*) exhibited strong ILS signals (ΔBIC > 10), whereas significant introgression was detected within the *pileatus* group (between *T. pileatus* and *T. geei*), and prominently in *T. delacouri*, which exhibited significant introgression signals across all tested triplets that could not be accounted for by ILS alone (Fig. 3A and Table S4).

To localize these gene flow signals onto specific internal branches, we utilized the *f*_branch_ (*f*_b_) statistic implemented in the D-suite framework^44^. Testing was conducted under both T2 and T1 topologies. Six recurrent and well-supported introgression events were identified. First, we detected gene flow between the *obscurus* group and the common ancestor of the *cristatus* group (Figs. 3B, S25 and Table S6; symbol 1). Second, broad genetic affinities were observed between the *francoisi* and *cristatus* groups (Fig S25; symbol 2), indicating extensive historical admixture between these clades. Third, consistent signals of introgression were inferred between the *francoisi* group and the ancestral branches leading to *T. phayrei*, *T. melamerus*, and *T. popa* (Figs. 3B, S25; symbol 3). Fourth, excess allele sharing was detected between the ancestral lineage of *T. laotum* and *T. hatinhensis* and the northern limestone langurs (Figs. 3B, S25; symbol 4). Fifth, evidence for secondary and recurrent gene flow involving the *francoisi* group was identified following the divergence of *T. melamerus* and *T. phayrei* (Figs. 3B, S25; symbol 5). Finally, particularly expansive introgression signals were concentrated around *T. crepusculus* (Fig. 3B, S25; symbol 6), suggesting a complex admixture history involving multiple ancestral sources. Deconstructing this complex admixture in *T. crepusculus* via *f*4-ratio statistics revealed primary genetic contributions from the *T. melamerus* + *T. phayrei* clade (∼18–21%), followed by *T. germaini* (∼17%) and the ancestral *francoisi* group (∼15%) (Fig. S26A), with these introgressed tracts exhibiting a discontinuous, mosaic distribution across the chromosomes (Fig. S26B), consistent with multiple, temporally distinct introgression events rather than a single admixture pulse.

To integrate these branch-specific introgression signals into a unified evolutionary framework, we reconstructed an admixture graph using qpGraph^45^. The best-fitting model achieved an optimal likelihood and the lowest worst residual (WR = 75.62) (Fig. S27–S33), indicating a satisfactory fit between the model and the observed allele frequency correlations. This graph fully corroborated the *f*_branch_ results by successfully resolving all six multi-directional introgression events within a single, coherent framework—including the complex, multi-source histories inferred for *T. crepusculus* and *T. delacouri*. Together, these complementary analyses demonstrate that ancient introgression across the *Trachypithecus* radiation was pervasive, highly heterogeneous, and a dominant force shaping genomic discordance through recurrent reticulation.

### Homoploid hybrid speciation of Delacour’s langur

To systematically test whether reticulate evolution has generated entirely novel lineages within the *francoisi* group, and to decipher the genomic architecture maintaining their evolutionary boundaries, we evaluated the hypothesized homoploid hybrid speciation (HHS) origin of Delacour’s langur (*T. delacouri*). Consistent with ADMIXTURE and PCA analyses positioning *T. delacouri* intermediately with mixed ancestry (Figs. 1D and S6), genome-wide scanning revealed comparable support for alternative affinities, with ∼40% of windows supporting the northern limestone clade (Tf1) and ∼33% supporting the southern clade (Tf2) (node 25; Fig. S12). Multi-locus phylogenetic analyses further corroborated this profound discordance, with the placement of *T. delacouri* alternating between sister to the northern and southern limestone clades across different genomic partitions (Figs. 4A and 4B). The *f*_branch_ (*f*_b_) statistics confirmed this dual affinity, showing that *T. delacouri* shares a significant excess of derived alleles with both the northern and southern clades (Figs. 4C and 4D). Because these signals cannot be explained by ILS alone (all tested triplets involving *T. delacouri* ΔBIC < −10; Fig. S34 and Table S4), we evaluated the hypothesis of HHS using the HyDe framework^8,46,47^. Hybridization signals were highly significant across all species combinations (P = 0; Z = 42.72–105.77), estimating substantial genomic contributions from both parental lineages (approximately 69.9%–71.2% derived from the northern lineage and 28.8%–30.1% from the southern lineage) (Fig. 4E; Tables S7 and S8). Subsequent qpAdm analyses further corroborated this admixture pattern, estimating approximately 72% and 28% ancestry contributions from the northern and southern lineages, respectively (Table. S9).

**Fig. 4.**
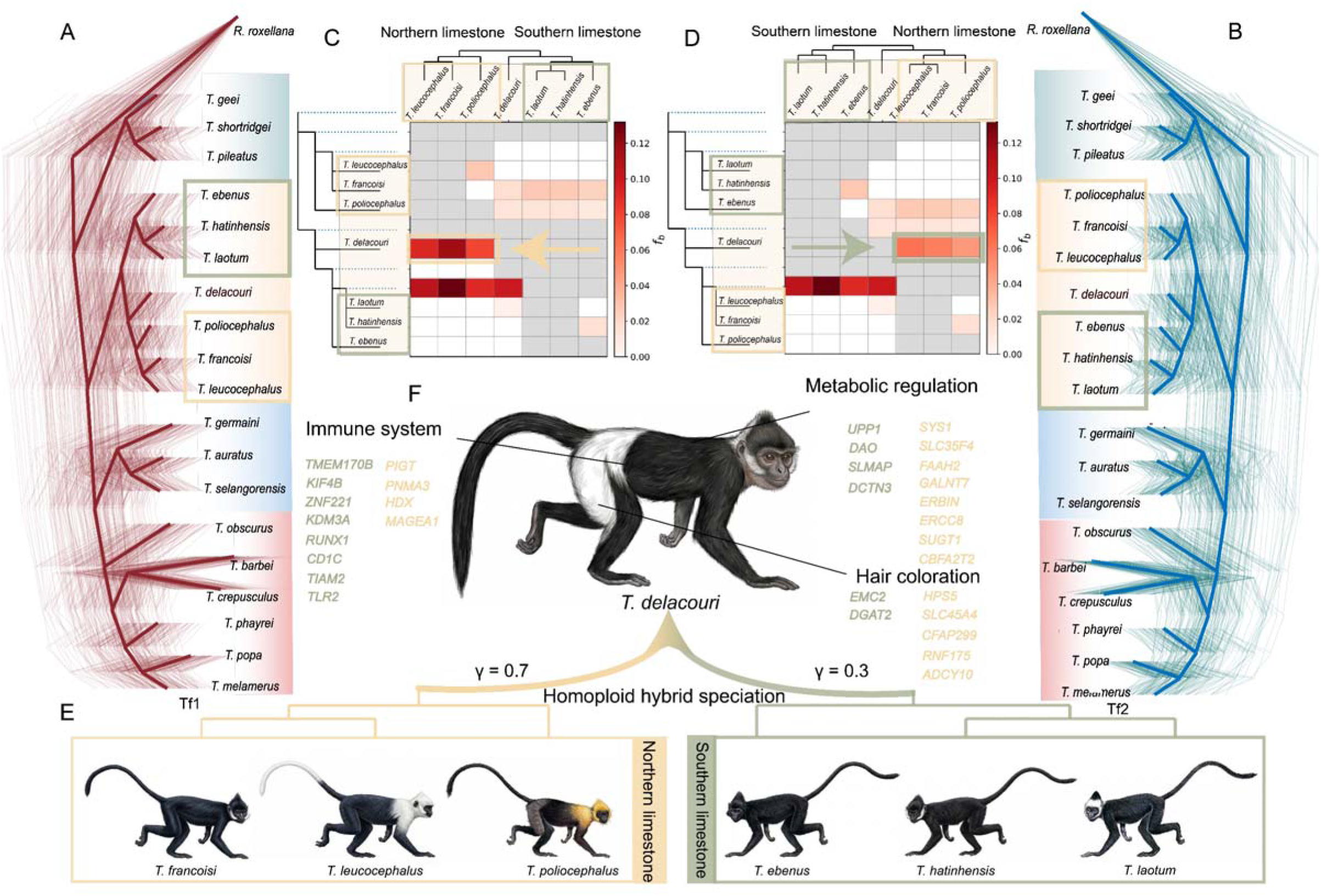
Genomic evidence for the homoploid hybrid speciation of *Trachypithecus delacouri*. **A, B** DensiTree visualizations illustrating the two dominant genome-wide phylogenetic topologies across the *Trachypithecus* genus. Trees were filtered and constructed using phybin. **C, D** Heatmaps of the *f*_branch_ statistics calculated via D-suite, revealing signals of historical gene flow between the northern and southern limestone langur clades. Arrows indicate the direction of introgression. **E** The proposed homoploid hybrid speciation model for *T. delacouri*, originating from the ancestral admixture between the northern (_γ_ = 0.7) and southern (_γ_ = 0.3) limestone langur lineages. **F** Core introgression genes and differentially inherited genes under positive selection associated with the immune system, metabolic regulation, and hair coloration in *T. delacouri*. Gene names in yellow indicate inheritance from the northern limestone lineage, while those in green denote inheritance from the southern limestone lineage. Drawings were generated using Gemini AI.

To map this adaptive hybrid mosaic, we integrated *F_d_*^48^ and *D_xy_*^49^ scans with ERICA (Evolutionary Relationship Inference using a CNN-based Approach)^50^. By utilizing convolutional neural networks, we rigorously disentangled true introgression tracts from the pervasive background noise of ILS. Through a stringent intersection of outlier windows, we identified 425 and 348 introgressed genes originating from the northern and southern limestone clades, respectively (Figs. S35, S36 and Table S10). Northern-introgressed genes were significantly enriched in inorganic ion transmembrane transport (Benjamini-Hochberg; *P* < 0.05) and cardiac/neural conduction systems (Benjamini-Hochberg; *P* < 0.05), whereas southern-introgressed fragments were predominantly associated with postsynaptic density (Benjamini-Hochberg; *P* < 0.05) and glutamatergic synaptic transmission (Benjamini-Hochberg; *P* < 0.05) (Figs. S35 and S36). Rigorous visual screening of the top 20 strongest loci pinpointed a core set of introgressed genes linked to extreme karst survival, encompassing immune regulators (*TLR2*, *CD1C*), metabolic networks (*UPP1*, *SLMAP*), and hair coloration (*DGAT2*, *RNF175*) (Figs. 4F, S37 and S38).

Crucially, a fixed, chimeric allele at the *RNF175* locus was found located within the introgressed region. This gene is integral to hair follicle morphogenesis and pigmentation fiber characteristics^51,52^. Driven by ancestral intra-genic recombination, *T. delacouri* inherited the 5’ region (encoding V9) from the northern clade and the 3’ region (encoding I165) from the southern clade, resulting in a novel ‘V+I’ amino acid configuration (Fig. 5A and 5B). This novel combination is fixed in *T. delacouri*, fundamentally contrasting with the “V+V” or “L+I” configurations of the parental populations. This hybrid-derived phenotypic shift presents a plausible genetic basis for the profound depigmentation characteristic of the ‘white trouser’ phenotype (Fig. 4F), which might have tentatively contributed to early sexual selection or prezygotic isolation^8,47,53^. Furthermore, to identify candidate genes underlying reproductive isolation, positively selected genes (PSGs) were scanned^54^ and 94 and 84 PSGs inherited by *T. delacouri* from its northern and southern limestone langur parental lineages, respectively (Tables S11 and S12). Notably, multiple alternatively inherited PSGs are strictly associated with reproductive system phenotypes. For instance, *SHCBP1L* and *CEP72* are critical for maintaining spindle integrity during male meiosis^55^, while *ADCY10*, *DNAH1*, *DNAH17*, *LRIT3*, *TTC21A*, and *ACADL* are essential for intraflagellar transport (IFT) and flagellar motility during spermatogenesis^56,57^. Furthermore, PSGs were also involved in melanogenesis and coat coloration, notably *SLC45A4*^58^ *HPS5*^59^ and *ADCY10*^60^, which have been shown to drive depigmentation and specific pelage colors in other vertebrates^58^. The alternate inheritance and selective fixation of these specific functional genes may have formed the core pre- and post-zygotic barriers maintaining the species identity of *T. delacouri*^54^.

**Fig. 5.**
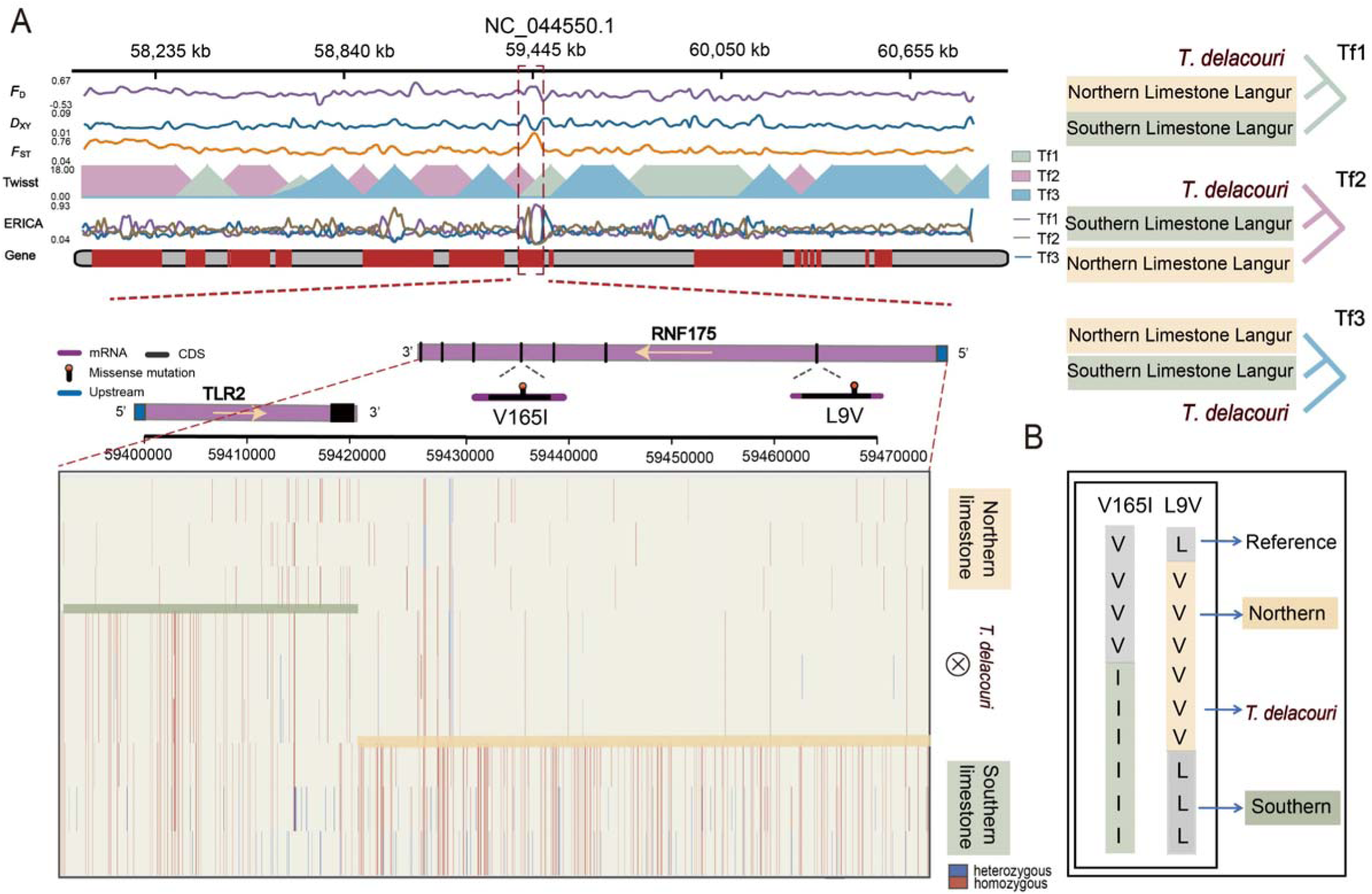
Local genomic landscape and intra-genic recombination of the introgressed *RNF175* allele in *T. delacouri*. **A** Local genomic landscape of adaptive introgression from northern limestone langurs into *T. delacouri* mapped on scaffold NC_044550.1. From top to bottom, the tracks represent: (1) *F*_d_ statistic, indicating the fraction of the genome shared via introgression between *T. delacouri* and the northern limestone clade; (2) *D*xy statistic, representing absolute sequence divergence between *T. delacouri* and the northern limestone clade; (3) *F*_ST_ statistic, showing relative genetic differentiation between *T. delacouri* and the southern limestone clade; (4) Twisst results, depicting local topology weighting for the three alternative within-group topologies Tf1, Tf2, and Tf3 across this genomic interval; (5) ERICA inference, showing local ancestry classification based on a CNN framework; and (6) Gene annotation, where red blocks indicate protein-coding regions. The red dashed box highlights the introgressed segment encompassing the *RNF175* gene. The zoomed-in schematic displays the gene structure and the locations of two key non-synonymous mutations (L9V and V165I). The bottom panel displays the visualized genotype landscape across this region for the northern limestone, *T. delacouri*, and southern limestone clades. Colors represent the genotypes at polymorphic sites (blue for heterozygous, red for homozygous). The horizontal bars within the heatmap clearly delineate a recombination breakpoint within the *RNF175* gene in *T. delacouri*. **B** The unique amino acid combination at the *RNF175* locus. Driven by ancestral intra-genic recombination, *T. delacouri* inherited the 5’ region (encoding V9) from the northern limestone clade and the 3’ region (encoding I165) from the southern limestone clade, resulting in a fixed, novel “V+I” chimeric allele, whereas parental populations retain the “V+V” or “L+I” configurations.

### Genomic erosion and the purging of severe genetic load

At the genus level, *Trachypithecus* species exhibit low levels of genetic diversity, with the critically endangered Cat Ba langur (*T. poliocephalus*) showing some of the lowest heterozygosity values reported among primates (Fig. S39). Within *Trachypithecus*, this overall paucity of diversity is accompanied by pronounced contrasts associated with geographic range size. Species with highly restricted distributions show a marked depletion of genome-wide heterozygosity alongside a substantial accumulation of long runs of homozygosity (*F*_ROH_) (Figs. 6A and B), indicative of recent and recurrent inbreeding. This pattern is most pronounced in *T. poliocephalus*^61^, which exhibits exceptionally low heterozygosity (0.28/kbp) and extensive genomic fraction contained within ROH (*F*_ROH_ = 0.865). Similarly, *T. delacouri* showed a severely depressed genome-wide heterozygosity of 0.47/kbp and an extensive *F*_ROH_ of 0.767, consistent with severe genomic erosion. This elevated homozygosity is accompanied by a shift in the architecture of genetic load, from masked load to realized load (Fig. 6C and 6D). In highly inbred species, recessive deleterious variants are increasingly exposed in the homozygous state, leading to a marked increase in the proportion of homozygous loss-of-function (LoF) variants (Fig. 6E). Notably, the proportion of homozygous LoF variants exceeds 82% across all *Trachypithecus* species, indicating that the vast majority of LoF mutations are no longer concealed in heterozygotes but instead contribute directly to realized genetic load.

**Fig. 6.**
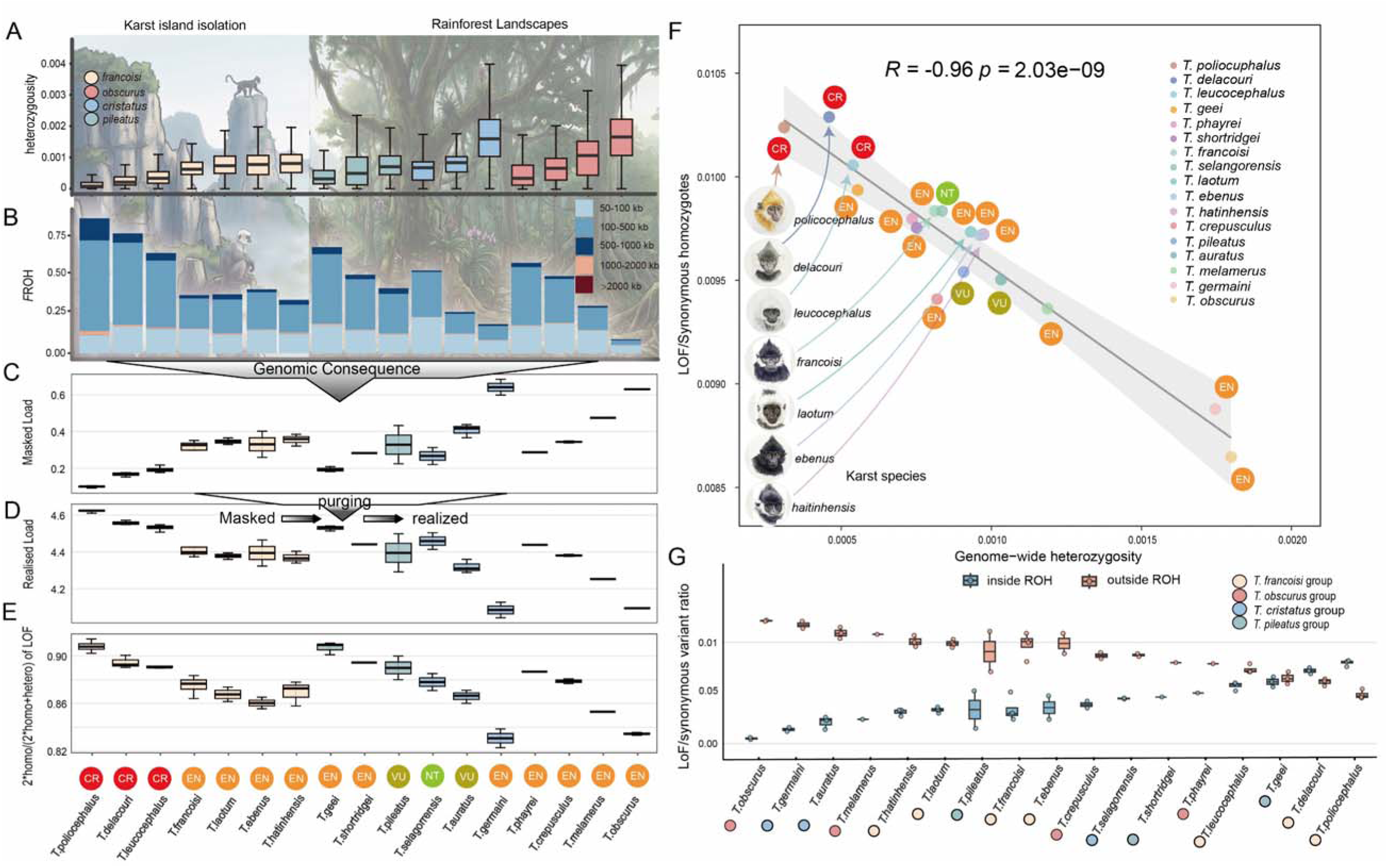
Genomic diversity, inbreeding, and genetic load across *Trachypithecus* species. **A** Genome-wide heterozygosity across *Trachypithecus*. The species are ordered along the shared x-axis corresponding to the labels at the bottom. Colors represent different species groups (*francoisi*, *obscurus*, *cristatus*, and *pileatus* groups). **B** The fraction of the genome in runs of homozygosity (*F*ROH). Stacked bar charts show the distribution of ROH lengths in different size categories (in kb). **C, D** Estimated masked and realized genetic load (deleterious alleles in heterozygous) **(C)** and realized genetic load (deleterious alleles in homozygous state) **(D)** across species. **E** The proportion of Loss of Function (LoF) alleles present in a homozygous state, calculated as 2*homo/(2*homo+hetero) of LoF sites. The circles below panel E denote the IUCN Red List conservation status for each species: CR (Critically Endangered, red), EN (Endangered, orange), VU (Vulnerable, olive), and NT (Near Threatened, green). **F** Pearson correlation analysis between genome-wide heterozygosity (x-axis) and the ratio of LoF to synonymous homozygotes (y-axis). Each point represents a species, colored by species group and annotated with its IUCN status. The grey shaded area represents the 95% confidence interval of the regression line. *R* and *p* values were calculated using two-sided Pearson correlation. **G** The ratio of LoF to synonymous variants located inside (blue) versus outside (red) Runs of Homozygosity (ROH) regions across different species. The karst and rainforest background was generated by Gemini AI.

To investigate whether this unmasking facilitated the selective removal of highly deleterious mutations, we analyzed the relationship between genetic diversity and mutational burden. This transition from masked to realized load is robustly supported by a strong positive correlation between genome-wide heterozygosity and masked genetic load (*R* = 0.89, *p* = 1.63×10^−6^. Fig. S40), alongside a striking negative correlation between genome-wide heterozygosity and realized load (*R* = −0.96, *p* = 2.03×10^−9^; Fig. 6F). Together, these dynamics demonstrate that highly restricted lineages, particularly those in fragmented karst landscapes, have largely depleted their reservoir of masked deleterious alleles. Instead, they bear a significantly higher realized genetic load, as recessive LoF mutations are increasingly forced into a homozygous state and exposed to natural selection^15,33^. Furthermore, across all *Trachypithecus* species, the LoF-to-synonymous variant ratio is consistently and profoundly depressed inside runs of homozygosity (ROH) compared to regions outside ROH (Fig. 6G). This localized depletion indicates that ROH efficiently facilitates the purging of recessive genetic load; once identical-by-descent inheritance exposes these severe, recessive LoF alleles in the homozygous state, they are rapidly selected against and selectively filtered out^62,63^. Taken together, these results indicate that prolonged demographic contraction and inbreeding in *Trachypithecus* have not only reshaped patterns of genome-wide diversity but have also fundamentally altered the balance between masked and realized genetic load. While purging within ROH may mitigate the most deleterious effects over time, the exceptionally high realized load observed in several species underscores the ongoing genomic vulnerability of these endangered primates.

## Discussion

### Resolving phylogenomic incongruence: XLRD versus genomic noise

Reconstructing the phylogeny of the Asian langur genus *Trachypithecus* presents one of the most formidable challenges in evolutionary biology. This complexity is driven not only by incomplete lineage sorting (ILS) but, more profoundly, by extensive and complex gene flow^16,17,22,29^. A comparison with other primate radiations highlights this distinct evolutionary dynamic: previous QuIBL analyses of the 120 triplets (trios) examined in the genus *Macaca* demonstrated that phylogenetic discordance is primarily ILS-driven, with 60.8% of trios diagnosed as “ILS only” and merely 5.8% attributed to introgression^64^. In stark contrast, our own analysis of 969 triplets in *Trachypithecus* reveals that while 22.7% of discordances result from ILS, a remarkable 54.9% are driven by significant introgression (Table S4). This widespread introgression, interacting with a massive ancestral effective population size (Ne ∼ 10^5^) (PSMC; Fig. S19) and the rapid Plio-Pleistocene radiation, creates a highly reticulated evolutionary history. Under such explosive diversification, alleles lacked sufficient time to coalesce between speciation events^65,66^, while frequent secondary contact facilitated genetic admixture, generating a profound hemiplastic cloud of genomic noise (Fig. S16)^3,4^. Disentangling these alternative evolutionary forces requires genomic windows resistant to both recombination and introgression. Like most primate species, genus *Trachypithecus* exhibit female philopatry accompanied by male-biased dispersal^67^. This behavioral ecology carries profound implications for phylogenomic reconstruction, particularly by enhancing the utility of the X chromosome. Given an approximately 1:1 sex ratio, females (XX) harbor two-thirds of the population’s X chromosomes. Under strict female philopatry, this majority fraction remains geographically anchored. Because migrating males (XY) possess only a single copy, they transmit X chromosomes across population boundaries at a significantly lower rate than autosomes^68,69^. Consequently, male-mediated gene flow intrinsically restricts X-linked introgression during secondary contact^70^. This sex-biased demography significantly shields the X chromosome from the homogenizing effects of admixture, rendering it an exceptionally reliable phylogenetic marker capable of retaining deep, ancient evolutionary signals despite pervasive genome-wide reticulation^70–72^.

Despite this perfect storm (massive ILS coupled with an enhanced risk of X-linked introgression),the X-linked recombination desert (XLRD)—a region characterized by suppressed recombination and enriched for reproductive isolation loci^38^—demonstrates remarkable evolutionary resilience. Our *f*4-ratio analysis explicitly quantifies this resilience revealing a striking 84.5% reduction in introgression within the XLRD compared to autosomes. This profound differential resistance directly explains the stark topological conflict observed in our phylogenomic reconstructions: while autosomes overwhelmingly support an admixed history (T1), the XLRD recovers the ancestral divergence (T2). Because severe recombination suppression within the XLRD prevents the decoupling of introgressed tracts from incompatible reproductive isolation alleles, these foreign genetic blocks are aggressively purged by purifying selection^38^. Consequently, autosomes are disproportionately homogenized by genome-wide admixture, whereas the XLRD serves as an impenetrable barrier that preserves the original divergence signal. Our analysis reveals that approximately 70% of the XLRD regions unequivocally support the true species tree topology (Figs. 2E and 2G). Furthermore, the remaining topological fluctuations (the ∼30% non-T2 signal) do not reflect a breakdown of this introgression barrier. Instead, they represent the unavoidable noise of residual deep ILS resulting from the massive ancestral effective population size (*N*e ∼10^5^) during the rapid Plio-Pleistocene radiation. Consistent with recent pan-mammalian genomic assessments, our findings confirm that the XLRD functions as an ancient speciation supergene; by acting as a barrier to interspecific gene flow, it suppresses introgression to retain an undistorted record of species divergence, even when genome-wide ancestry is otherwise thoroughly dominated by admixture^38^.

### Reticulate evolution as a creative engine for colobine diversification

Historically, phylogenetic studies of the genus *Trachypithecus* have been plagued by profound incongruence, as traditional morphological classifications and early molecular analyses yielded fundamentally conflicting topologies^16,17,22,29^. Early inferences heavily relied on short mitochondrial DNA (mtDNA) fragments; however, these maternal markers are exceptionally susceptible to historical introgression and incomplete lineage sorting^73–75^. Furthermore, pervasive interference from nuclear mitochondrial DNA (NUMTs) frequently precipitated erroneous conclusions (e.g. misplacing the *pileatus* group as a sister clade to *Semnopithecus*) thereby spawning inaccurate hypotheses of ancient hybridization events^76,77^.

Our genomic analyses reveal that the biological root of this historical confusion is the profoundly reticulate evolutionary history of the genus (Fig. 3B). We demonstrate that widespread gene tree discordances across *Trachypithecus* are far too extensive to be fully explained by ILS alone (Fig. S24 and Table S4). Crucially, these events are not isolated; many involve recurrent (e.g. the *francoisi* group following the divergence of the *T. melamerus* and *T. phayrei* lineage; Fig. S25; symbol 5) or even multi-source introgression into a single lineage. A striking example is the Indochinese grey langur (*T. crepusculus*), where our genetic dissection uncovered complex admixture from at least three distinct ancestral sources (Fig. S26). Such multifaceted introgression origins perfectly illustrate why previous studies struggled to confidently resolve the phylogenetic placement of this and similar taxa^78^. Nevertheless, we interpret these specific introgression patterns with caution, as complex signatures can sometimes emerge as analytical artifacts from topological misspecification, intricate network structures, or signal ‘smearing’ across closely related nodes^44,79^—a known limitation of triplet-based methods that lack the power to strictly distinguish introgression directions^79–81^. Therefore, while the multi-source hybrid nature of *T. crepusculus* is evident, tracing its precise introgression pathways warrants further validation with more extensive population and taxon sampling.

Yet, the evolutionary consequences of such pervasive gene flow extend far beyond methodological complications. While complex admixture webs can blur species boundaries and act as a confounding ‘phylogenetic nuisance,’ our results demonstrate that reticulate evolution simultaneously operates as a profound generative force. Conventionally, morphological taxonomy has classified Delacour’s langur (*T. delacouri*) as a distinct species of the *francoisi* group^16,17^. However, in-depth genomic scans, D-statistics (Fig. 4C and 4D), and HyDe admixture analyses (Fig. 4E) conclusively demonstrate that it actually originated from an ancient, deep hybridization event between the ancestors of the northern limestone clade (including *T. leucocephalus*, *T. francoisi*, and *T. poliocephalus*) and the southern limestone clade (including *T. laotum*, *T. hatinhensis*, and *T. ebenus*) (Fig. 4E). Building upon this profound genomic mosaicism, our analysis elevates *T. delacouri* as a rare primate instance of homoploid hybrid speciation (HHS) (Fig. 4E)^8,47,53^.

Much like the reticulate origins documented in the gray snub-nosed monkey (*Rhinopithecus brelichi*), hybridization in *T. delacouri* has clearly functioned as a potent creative evolutionary engine that radically accelerates ecological adaptation (Fig. 4F). Karst landscapes are inherently harsh, characterized by steep limestone cliffs, alkaline soils, and exceptionally high concentrations of calcium and other metal ions that pose severe physiological challenges, including potential cellular and genetic toxicity^28,82^. Survival in these unique ecological niches requires profound, multi-system evolutionary innovations, including the accelerated adaptive evolution of ion channels (such as voltage-gated sodium, potassium, and calcium channels) to maintain intracellular homeostasis amidst high mineral intake, as well as the enhancement of DNA damage response and repair pathways^28,82^.

However, adapting to such multifaceted extremes through *de novo* mutations alone is evolutionarily protracted^83^. HHS mitigated this bottleneck by synthesizing a modular genetic toolkit that seamlessly integrates these broad, ancestral physiological adaptations with discrete introgressed elements from both parental lineages^84^. Simultaneously, the adaptive introgression of a polygenic immune module (i.e., encompassing key loci such as *TLR2*, *CD1C*, and *RUNX1*) likely conferred the hybrid lineage with enhanced immunological resilience against localized pathogenic pressures, or provided a unique combination of immunological defenses distinct from either parental lineage (Fig. 4F). Beyond these physiological benefits, this adaptive landscape is reinforced by a heavy concentration of alternatively inherited PSGs that erect a dual reproductive barrier framework. On one hand, the alternate inheritance of key pigmentation-related PSGs—notably *SLC45A4*^58^, *HPS5*^59^, and *ADCY10*^60^—systematically coupled with the fixed *RNF175* chimeric variant (which combined the 5’ region encoding V9 from the northern clade and the 3’ region encoding I165 from the southern clade; Fig. 5A and 5B) to drive the profound depigmentation characteristic of the ‘white trouser’ phenotype (Fig. 4F). In the visually dominant sensory ecology of primates, highly contrasting pelage patterns serve as primary specific-mate recognition systems^8,47,53^. Therefore, this multi-gene driven visual divergence likely triggered strong sexual selection^16,22^, establishing an immediate prezygotic barrier against parental back-introgression and stabilizing the new hybrid lineage. On the other hand, multiple alternatively inherited PSGs are strictly tied to intrinsic postzygotic isolation; these include *SHCBP1L* and *CEP72* (essential for spindle integrity during male meiosis) alongside *ADCY10*, *DNAH1*, and *DNAH17* (critical for flagellar motility during spermatogenesis). The selective fixation of these alternative functional alleles may have created postzygotic barriers that effectively insulated the hybrid genome from subsequent genetic swamping^8,10^. Ultimately, rather than a mere blending of genomes, this hybrid divergence provided the exact synergistic adaptive capacity required for *T. delacouri* to successfully establish and maintain its unique evolutionary trajectory under its specific environmental regime, facilitating localized specialization amid the wider karst landscape^9,13,64^.

### Genomic erosion and conservation implications for limestone langurs

Prolonged isolation and anthropogenic threats have driven limestone langurs to the brink of extinction^61,85^. Conservation genomics now reveals the profound crises concealed within these specialized lineages. Given that over 70% of *Trachypithecus* species are currently classified as Endangered or Critically Endangered by IUCN^31^, establishing robust conservation genomics frameworks for this lineage is of paramount importance. While we deliberately excluded historical museum specimens to ensure the highest reliability of our population-level genomic inferences, the genomic data for certain lineages (*T. auratus* and *T. obscurus*) were derived from captive individuals. Because artificial environments typically drive rapid depletion of nucleotide diversity and expansion of ROH^86,87^, our metrics likely overestimate natural inbreeding levels for those two species^88^.

Crucially, our findings offer a compelling perspective on the evolutionary paradox posed by their reticulate history: whether the adaptive advantages of hybridization can be sustained under extreme geographic restriction. The genomic evidence points toward a complex evolutionary trade-off. While ancient admixture likely equipped lineages like *T. delacouri* with the initial chimeric ‘adaptive toolkit’ to colonize isolated karst niches, this early hybrid advantage appears to have been gradually attenuated. Prolonged confinement within these fragmented limestone ‘islands’ ultimately seems to have offset these ancestral advantages over time. Extreme historical isolation and pervasive homozygosity have driven a dangerous accumulation of realized genetic load (Fig. 6B, 6C and 6D). This profound architectural shift has simultaneously catalyzed an intense regime of purifying selection. By forcing historically masked, highly deleterious mutations into phenotypic expression, massive ROH tracts have functioned as a severe genomic crucible (Fig. 6G), aggressively shedding fatal recessive alleles from the gene pool^15,63^. Consequently, these severely restricted lineages exhibit a ‘fragile resilience’—they persist in extreme karst environments specifically because they have paid the demographic price to purge their most lethal defects^15,62,63,89,90^.

## Conclusion

This study reconstructs the evolutionary history of the Asian langur genus *Trachypithecus*, revealing a radiation shaped by ancient divergence, recurrent admixture, and demographic isolation. Notably, the identification of Delacour’s langur as a homoploid hybrid species underscores ancient admixture as a powerful driver of rapid evolutionary innovation. However, the long-term genomic consequences of this reticulate evolution are profound. In lineages restricted to fragmented karst landscapes, we detect a stark transition from masked deleterious variation to a realized genetic load. Driven by sustained population reductions, this shift exposes a cryptic but critical threat to population viability and severely elevates extinction risk.

Ultimately, ancient admixture exerts dual evolutionary effects: it facilitates early diversification while predisposing derived lineages to long-term genomic erosion under isolation. To mitigate this escalating mutational burden, conservation strategies must prioritize restoring connectivity through ecological corridors to rescue gene flow among isolated populations. Concurrently, minimizing anthropogenic disturbances is essential to preserve their specialized limestone habitats. Integrating these in situ conservation efforts with genomic monitoring will be critical for safeguarding the long-term resilience of *Trachypithecus* langurs.

## Materials and methods

### Sample collection and library preparation

We obtained 12 *Trachypithecus* specimens for whole-genome sequencing: 10 historical museum samples (6 *T. crepusculus*, 2 *T. barbei*, and 2 *T. popa*) sourced from the natural history museums in Chicago, New York, London and Leiden, and two *T. selangorensis* blood samples from the Gene Bank of Primates (German Primate Center, Göttingen, Germany). For museum specimens, skin scrapings were collected from the interior of damaged sections to minimize external contamination.

To prevent exogenous and cross-sample contamination, all museum DNA extractions and library preparations were performed in a dedicated ancient DNA (aDNA) facility following stringent decontamination protocols (e.g., UV irradiation, positive air pressure, and negative controls). For historical samples, including the holotype, DNA was extracted using a column-based method optimized for the recovery of degraded, short fragments^91,92^. For blood samples, total genomic DNA was extracted using the DNeasy Blood & Tissue Kit (Qiagen) following the manufacturer’s instructions. DNA quantity and integrity were assessed using a Qubit 4.0 fluorometer (Thermo Fisher Scientific) and a Bioanalyzer 2100 or TapeStation (Agilent Technologies). For historical samples, sequencing libraries were constructed using the NEBNext Ultra II DNA Library Prep Kit (New England Biolabs) with approximately 50 ng of input DNA. To maximize recovery, adapter-ligated libraries were purified without size selection. For blood samples, double-stranded libraries were prepared using the Blunt-End Single-Tube (BEST) protocol^93^ with dual-indexing. All libraries, including a DNA extraction negative control, were quantified via qPCR using the NEBNext Library Quant Kit. Final libraries were sequenced on the Illumina NovaSeq 6000 platform (150 bp paired-end) at a target depth of ∼30X. Raw sequencing data are available under accession PRJCA062116 at the Genome Sequence Archive (https://ngdc.cncb.ac.cn/gsa) of China’s National Genomics Data Center.

In addition, we integrated genomic data from 41 other *Trachypithecus* individuals and two *Rhinopithecus* species (*R. roxellana* and *R. bieti*) obtained from the NCBI Short Read Archive (accession no. show in Table S1). In total, a comprehensive dataset comprising 55 individuals—representing 19 *Trachypithecus* species and two *Rhinopithecus* individuals as outgroups—was utilized for subsequent phylogenetic analyses. The present study was approved by the Animal Ethics Committee of the Institute of Zoology, Chinese Academy of Sciences. The procedure of blood and tissue collection was in strict accordance with the Animal Ethics Procedures and Guidelines of the People’s Republic of China.

### Phylogeny of mitogenomes

NOVOPlasty v4.3^94^ was used to *de novo* assemble the mitochondrial genomes (mitogenomes) of all samples. They were annotated with MitoZ v2.4^95^. Control regions were manually excluded from each sequence prior to alignment to avoid alignment artifacts associated with high variability. The remaining mitochondrial DNA sequences were aligned using MAFFT v7^96^. Indels and poorly aligned positions were removed by Gblocks v0.91b^97^. A ML tree was reconstructed using RAxML v8.2.12^98^ with a GTRGAMMA substitution model and 1,000 bootstrap replicates. The tree was visualized with FigTree v1.4.4 (http://tree.bio. ed.ac.uk/software/figtree/) and iTOL v5 (https://itol.embl.de/).

### Genome-wide variant calling and quality control

Raw sequencing reads were processed with fastp v0.20^99^ to remove adapter sequences and low-quality bases. The resulting paired-end reads were aligned to the *Rhinopithecus roxellana* reference genome (GCA_007565055.1) using Burrows-Wheeler Aligner (BWA) mem v0.7.17^100^. Subsequent BAM files were sorted using the sort function in SAMtools v1.9^101^, and PCR duplicates were identified and removed using markdup with the -r flag. To ensure high-quality alignments, unmapped reads and those with a mapping quality (MAPQ) score below 30 were filtered out using the view function (parameters: -F 4 -q 30). Finally, we employed mapDamage v2^34^ to assess potential DNA damage patterns, specifically C-to-T transitions resulting from deamination, in the historical skin samples. While eight samples exhibited characteristic 5’ and 3’ cytosine deamination, four (Tcre_598, Tcre_599, Tcre_600, and Tpop_674) lacked these diagnostic signatures (Fig. S1, Table S1). Notably, Tpop_674 yielded an endogenous content of only 60.92%, suggesting potential modern contamination or poor preservation. While these four individuals were included in the phylogenetic tree to maximize taxonomic coverage, they were excluded from downstream analyses (e.g., introgression inference) to avoid potential biases from DNA damage or modern contamination. Since *T. crepusculus* and *T. barbei* remained well-represented by other high-quality samples, this exclusion did not compromise our analytical power.

Variant calling was done for each sample using GATK v4.2^102^ HaplotypeCaller to generate GVCFs, followed by joint genotyping using CombineGVCFs and GenotypeGVCFs. Insertions and deletions (InDels) were excluded from the downstream analysis. Genotypes exhibiting extreme coverage depths were masked. Autosomes required masking if coverage was below 0.5 × or above 2 × the sample average, whereas separate criteria were applied to the X chromosome. The Y chromosome was excluded entirely. Subsequent SNV filtering was done using GATK VariantFiltration with stringent thresholds (QD < 2.0; QUAL < 30.0; SOR > 3.0; FS > 60.0; MQ < 40.0; MQRankSum < −12.5; ReadPosRankSum < −8.0). In addition, heterozygous genotypes with minor allele support below 0.25 were masked. Finally, repetitive regions within the reference genome were identified and masked using the snpable pipeline (k-mer size = 100 bp). Additional filters were used to remove SNVs with missingness ≥10%, quality < 20 (phred-scale), and non-biallelic sites. All variants were annotated through SnpEff v4.3^103^ using the GCA_007565055.1 GFF3 annotation from the NCBI (Table S2).

### Inference of kinship in the population

To examine the kinship coefficient within a species, we estimated the relatedness of the samples in each population using KING v2.1.3^104^ with default parameters. An estimated kinship coefficient range >0.354, [0.177, 0.354], [0.0884, 0.177], and [0.0442, 0.0884] corresponds to duplicate sample/monozygotic twins with 1^st^-degree, 2nd-degree, and 3rd-degree relationships, respectively. Crucially, kinship inference using KING confirmed the absence of cryptic relatedness greater than the second degree (coefficient > 0.177), justifying the retention of all samples (Fig. S3).

### Phylogenetic reconstruction using the window-based strategy

Consensus sequences for all individual genomes were generated from VCF files and aligned on a per-chromosome basis, excluding sex chromosomes and mitochondrial scaffolds. During this process, we specifically omitted variants present in the *Rhinopithecus roxellana* reference that were invariant within *Trachypithecus*. The per-chromosome alignments were partitioned into non-overlapping windows of 50, 100, and 200 kb. The genomic windows within 5 Mb of both chromosomal ends, as well as those within 5 Mb upstream and downstream of the centromere, were removed. For each window, ML gene trees were reconstructed using RAxML v8.2.12^98^ with 100 bootstrap replicates under the GTR + GAMMA substitution model. Based on the 50, 100, and 200 kb window trees, a species tree was inferred using ASTRAL v5.7.8^35^ under the multispecies coalescent model, with quartet scores and local posterior probabilities calculated to assess nodal support. The resulting species tree was re-rooted at the common ancestor of the golden snub-nosed monkey (*R. roxellana*) and the black snub-nosed monkey (*R. bieti*). To explore phylogenetic conflict, consensus networks of the window trees were generated using SplitsTree v5^37^ with varying thresholds. Genealogical discordance across the genome was further quantified using Twisst v0.2^36^. Finally, all gene tree topologies were clustered via PhyBin (https://github.com/rrnewton/PhyBin), and the top two dominant topologies were visualized using DensiTree v2.6.3^105^.

### Phylogenetic reconstruction using the X-linked low-recombining desert

To reconstruct a species tree and mitigate the confounding effects of interspecific gene flow and ILS, we focused our phylogenetic analysis on the X-linked recombination desert (XLRD)^38^. The XLRD is a large, evolutionarily conserved region characterized by significantly suppressed recombination, which acts as a recurrent barrier to gene flow and has been shown to consistently retain the true species history even when introgression dominates the rest of the genomic landscape^38^. Following the genomic coordinates identified in previous mammalian studies (spanning approximately the interval between *JADE3* and *CHRDL1* located at chromosome X), we first generated a genome-wide chain file between our reference genome to T2T-CHM13 (GCF_009914755.1). Specifically, we used minimap2 v2.1^106^ with the -cx asm5 parameter to generate a Pairwise Mapping Format (PAF) file. Subsequently, we employed transanno v0.4.4 (https://github.com/informationsea/transanno) to construct the chain file between the two reference assemblies. Coordinate transformation of our VCF files was then performed using the transanno liftvcf command. To isolate the phylogenetic signal from this “speciation supergene”, we extracted the VCF data corresponding to the XLRD region and generated consensus sequences using bcftools v1.7^107^. Finally, we reconstructed a ML phylogenetic tree using RAxML v8, employing the GTR + Gamma substitution model with 100 bootstrap replicates to assess nodal support (Fig. 2A).

### Phylogenetic analysis of concatenated whole-genome SNV data

SNVs were extracted from the filtered VCF file and converted into individual consensus sequences (pseudo-haplotypes) using vcf2phylip v2.0 (https://doi.org/10.5281/zenodo.2540861). These sequences were concatenated to generate a comprehensive multiple sequence alignment (MSA) encompassing all individuals. Based on this alignment, autosomal and X-linked SNVs were separately extracted and ML phylogenetic trees were reconstructed using RAxML v8.2.12 with 100 bootstrap replicates under the GTR + GAMMA substitution model. The resulting phylogenies were visualized and refined using FigTree v1.4.4 and iTOL v5. To further characterize the genetic relationships among samples, Principal Component Analysis (PCA) was performed on the filtered autosomal SNVs using the EIGENSOFT package v5.0^108^. Unsupervised clustering was performed with ADMIXTURE v1.3.0^109^ using the same SNV dataset. Cross-validation was conducted for K values ranging from 2 to 8 (three replicates each), with K = 4 selected based on the lowest CV error. The convergence threshold was set to 0.0001 (-c 0.0001).

### Quantifying Introgression with QuIBL

To distinguish between incomplete lineage sorting (ILS) and introgression, we employed QuIBL (https://github.com/miriammiyagi/QuIBL)^3^ to analyze the internal branch length distributions across all evaluated species triplets. QuIBL estimates the proportion of introgression and calculates the probability that a specific locus aligns with either an “ILS-only” model or a “combination of ILS and introgression” model.

Given QuIBL’s sensitivity to recombination, we extracted 20-kb windows separated by 400-kb intervals from each sample to minimize the risk of including recombination breakpoints within any single window. We then filtered the inferred ML trees, retaining only those with at least 10 parsimony-informative sites. This resulted in a final dataset of 5,050 trees used as input for QuIBL. Using the established species tree topology, we assigned appropriate outgroups to each triplet and calculated the percentage of loci that supported discordant topologies with significant evidence of introgression.

To determine whether the internal branch lengths were best explained by ILS alone (Scenario 1) or a combination of ILS and introgression (Scenario 2), we compared their likelihoods using the Bayesian Information Criterion (BIC). The model fit was assessed via the difference in BIC values ΔBIC (BIC_scenario2_-BIC_scenario1_). Since the BIC value is less than 0, when ΔBIC is greater than 10, the scenario of ILS only with the lower BIC value is preferred. Conversely, when ΔBIC is less than −10, the scenario of a mixture of ILS and introgression with the lower BIC value is preferred. In all other cases (−10 < ΔBIC < 10), the two scenarios are considered statistically indistinguishable (Tables S4 and S5).

### Phylogenomic analysis and divergence time estimation

Coding sequences (CDS) and protein sequences for representative primate species—including *Trachypithecus francoisi* (GCA_009764315.1), *Piliocolobus tephrosceles* (GCF_002776525.5), *Rhinopithecus roxellana* (GCF_007565055.1), *Macaca fascicularis* (GCF_037993035.1), *Homo sapiens* (GCF_009914755.1), and *Pan troglodytes* (GCF_028858775.2)—were retrieved from the NCBI database. To ensure genomic consistency, *Trachypithecus* sequences were mapped to the *R. roxellana* reference genome (GCF_007565055.1), and CDS were extracted based on shared genomic coordinates. Single-copy orthologous genes across all species were identified using OrthoFinder v2.5.4^110^. Subsequently, the CDS of these orthologs were aligned using PRANK (v.170427) ^111^ under a codon-based model (-codon).

To minimize the impact of selection pressure on time estimation, four-fold degenerate (4D) sites were extracted using MEGA7^112^ and concatenated into a super-matrix. Divergence times were estimated using the MCMCTree program within the PAML v4.9 package^113^. We employed the independent rates (clock=2) model to account for rate heterogeneities among lineages.

Fossil calibrations were strategically applied following established frameworks ^23,85,114–116^. Specifically, the Hominoidea-Cercopithecoidea divergence was constrained to 29.2 – 33.1 million years ago (Mya) based on the Oligocene fossil *Saadanius hijazensis*. Additionally, the divergence between *Homo* and *Pan* was calibrated at 6.0–7.0 Mya. The split between Cercopithecinae and Colobinae was set at 18.0 – 20.6 Mya, while the divergence between African colobines (*Piliocolobus*) and Asian colobines (*Rhinopithecus* and *Trachypithecus*) was calibrated at 10.0–12.5 Mya. Within the Asian colobine clade, the divergence of odd-nosed monkeys from classical langurs was set with a prior of ∼7.5 Mya (95% HPD: 6.5–8.5 Mya).

The MCMC analysis was performed with a burn-in of 1,000,000 iterations, followed by 5,000,000 iterations sampled every 100 steps to ensure a sufficient effective sample size (ESS > 200). To verify convergence, independent duplicate runs were conducted starting from different random seeds, yielding highly consistent posterior time estimates for the *Trachypithecus* radiation.

### Analysis of demographic history using genome-wide data

To estimate the demographic history of each *Trachypithecus* individual, we performed Pairwise Sequentially Markovian Coalescent (PSMC, v0.6.5-r67)^42^ analysis. Input files were generated using Samtools v1.7^101^ mpileup with the parameter “-C 50” to improve mapping quality, and only autosomal SNVs were retained to construct the diploid consensus sequences. PSMC was run for 25 iterations (-N 25) with a maximum 2N0 coalescent time (-t) of 15 and an initial theta/rho ratio (-r) of 5. To minimize false-positive fluctuations in our demographic inferences, we set the atomic time interval pattern to “4+25*2+4+6“^117^. The resulting PSMC plots were scaled using a mutation rate (μ) of 7.2×10^−9^ per site per generation and a generation time (g) of 12 years for all *Trachypithecus* species^31,85^.

### Ecological niche modeling

A total of 6,890 occurrence localities of *Trachypithecus* langurs were obtained from the Global Biodiversity Information Facility (GBIF; http://data.gbif.org). To ensure data quality, we applied the R package *CoordinateCleaner*^118^ to eliminate duplicate entries and coordinates with geo-referencing inaccuracies. Spatial autocorrelation and potential model overfitting were mitigated by applying a spatial filtering technique: we established a 10,000-m circular buffer around each site, randomly retaining only a single occurrence per buffer. This procedure yielded a final refined dataset of 1,350 distribution points for subsequent analyses.

We used 19 bioclimatic variables available from the WorldClim v1.4 database^119^ as environmental predictors across three target epochs: the contemporary period, the Last Glacial Maximum (LGM), and the Last Interglacial (LIG) (http://www.worldclim.org/). All climate layers were standardized to a spatial resolution of 2.5 arc-minutes. Specifically, the LIG dataset was resampled from its original 30 arc-second resolution to 2.5 arc-minutes using the ‘aggregate’ function in the R *raster* package^120^. The study extent was then defined by cropping the climate layers to encompass the known distribution range of Southeast Asian langurs (70 to 135 E; −15 to 40 N). To address multicollinearity among the environmental variables, a Pearson correlation analysis was conducted using the extracted point data. For any highly correlated variable pair (|r| ≥ 0.8)^121^, the variable demonstrating a lower predictive contribution was excluded from the final modeling process.

Species distribution models across the three time periods were generated using MaxEnt v3.4.4^43^, incorporating the spatially filtered occurrence data and the rigorously selected bioclimatic variables. Model calibration and evaluation were performed by partitioning the occurrence data into a 75% training subset and a 25% testing subset. To assess model robustness and stability, we executed 10 cross-validation replicates, keeping all other MaxEnt parameters at their default configurations (Fig. S20 -S23).

### Introgression analysis using *D*-statistics

We performed *D*-statistic analyses using Dsuite v0.5^44^ across all possible combinations of the 19 *Trachypithecus* species. To account for phylogenetic uncertainty, we tested two alternative phylogenetic hypotheses as input topologies. Specifically, the Dtrios module was employed to calculate *D*-statistics and *f*4-ratio for all possible trios following the topology (((P1, P2), P3), Out), with *R. roxellana* and *R. bieti* serving as outgroups. This approach yielded a total of 969 groupings for each candidate topology. Subsequently, the *f*branch module was applied to assign introgression signals to specific branches within these topologies. The resulting statistics were visualized using the dtools.py script provided in the Dsuite package (Table S6).

### Modeling admixture graphs with qpGraph

To reconstruct the complex branching and admixture history of *Trachypithecus*, we employed the ADMIXTOOLS2 v2.0.0 framework^45^, which provides an efficient implementation for fitting and comparing admixture graphs. To minimize the impact of physical linkage while retaining sufficient genomic information, we thinned the genome-wide SNV dataset by selecting one SNV per 100-bp window, resulting in a final set of 26,721,227 SNVs. We first computed a global matrix of *f*_2_-statistics to serve as the basis for model fitting.

Using the inferred species tree (T2) as the starting topology, we iteratively added admixture edges to explore alternative reticulate scenarios. We evaluated the fit of each model using the qpgraph() and fstat() functions, prioritizing models with the lowest worst residuals (the maximum difference between observed and predicted *f*_2_-statistics). Furthermore, we utilized out-of-sample scores, a cross-validation approach unique to ADMIXTOOLS2, to perform unbiased comparisons between competing models, ensuring that the selected graph provided the most parsimonious and statistically robust representation of the group’s evolutionary history.

### Hybridization detection with HyDe

To further validate the introgression patterns identified by D-statistics, we employed HyDe v0.4.3^46^ (Hybridization Detection). This method automates the detection of hybridization across numerous species while accounting for ILS, allowing for hypothesis testing at both population and individual levels by estimating the admixture proportion (γ).

The analysis was conducted as follows: first, we used the run_hyde_mp.py script to perform population-level tests on all possible triplets within the *Trachypithecus* lineages, using *R. roxellana* and *R. bieti* as outgroups. Results were filtered based on statistical significance (*P* values <0.05). In this framework, the γ value quantifies the contribution of parental lineages: a γ value of approximately 0.5 indicates a balanced (50:50) hybrid, while values near the extremes (e.g., 0.01 or 0.99) suggest low levels of admixture, which may reflect more ancient hybridization or other evolutionary processes.

Our initial screening revealed that hybridization events with a 30:70 ratio were significantly concentrated among the northern limestone langurs (*T. leucocephalus*, *T. francoisi*, and *T. poliocephalus*), southern limestone langurs (*T. laotum*, *T. hatinhensis*, and *T. ebenus*), and *T. delacouri*. To confirm these findings at a finer scale, we tested each individual within the putative hybrid population (*T. delacouri* clade) using specified triplets (Tables S7 and S8). Finally, we performed bootstrap resampling with 500 replicates using the bootstrap_hyde.py script to assess the robustness of the hybridization evidence within these specific lineages.

### Admixture proportion estimation using qpAdm

To formally model the admixture history and estimate ancestry proportions for the target lineage, we employed qpAdm implemented in the ADMIXTOOLS2 v2.0.0 package^45^. This method relies on a matrix of *f*4-statistics to assess the goodness-of-fit of user-defined demographic models. In our framework, the “left” populations comprised the target species alongside the northern and southern limestone langur groups, which represented the potential proxy sources. A set of distantly related taxa was assigned as the “right” populations to serve as differential references. We systematically iterated through all possible combinations of northern and southern limestone langurs as paired source populations. A given admixture model was considered statistically plausible and a good fit if it yielded a *P*-value > 0.05 and the inferred admixture proportions fell within the biologically meaningful range of 0 to 1.

### Introgression analysis with *F*_d_ and *D*_xy_ values

To identify putatively introgressed regions across the *Trachypithecus* genomes, we implemented a scanning strategy based on the methods described by Martin *et al*.^122^ and van der Valk *et al*.^49^. This approach is predicated on the expectation that introgression between species within a specific genomic region significantly reduces interspecific divergence relative to the genomic background. Using the genomics_general Python toolkit (https://github.com/simonhmartin/genomics_general), we calculated *F*_d_ and *D*_xy_ values in 10-kb sliding windows across the entire genome for each *Trachypithecus* species. Both statistics are sensitive to the density of informative sites within each window; therefore, these metrics were utilized to pinpoint regions with significantly reduced divergence and elevated shared variation. Genomic regions demonstrating a joint signature of elevated shared variation (top 3% of *F*_d_ values) and minimized absolute sequence divergence (bottom 3% of *D*_xy_ values) were rigorously identified as introgressed candidate segments (Table S9). Functional enrichment analysis was performed on overlapping gene sets using clusterProfiler v4.6.0 (R package)^123^ (Fig. S35B and S36B).

### Introgression analysis with efficient local ancestry inference (LAI)

To resolve the genomic mosaic of introgression at a finer scale, we utilized ERICA (Evolutionary Relationship Inference using a CNN-based Approach), a deep learning-based framework designed to identify local introgressed regions directly from sequence data^50^. Unlike conventional methods that require species-specific demographic modeling, we implemented ERICA using its pre-trained default models. These models were trained on a generalized dataset covering a broad spectrum of evolutionary scenarios, including varying degrees of ILS and gene flow across diverse divergence times, ensuring robust performance across different taxa^50^.

In our analysis, we specifically investigated the local ancestry of the putative hybrid lineage *T. delacouri*, by evaluating its relationship with northern limestone langurs (*T. leucocephalus*, *T. francoisi*, and *T. poliocephalus*) and southern limestone langurs (*T. laotum*, *T. hatinhensis*, and *T. ebenus*). To identify which specific genomic regions were inherited from either parental clade, we performed a sliding window scan across the genome. Following the author’s recommendations for optimal resolution and accuracy, we defined a window size of 10 kb (-w 10000). Genomic segments supporting alternative (discordant) topologies were identified as introgressed regions using a stringent probability threshold of 0.4 (-d 0.4), a value suggested to effectively distinguish true introgression signals from background noise caused by ILS.

### Genomic scans for positively selected genes

To identify positively selected genes (PSGs) associated with the hybrid speciation of *T. delacouri*, we established two comparative grouping strategies: (1) the merged population of *T. delacouri* and northern limestone langurs compared against southern limestone langurs; and (2) the merged population of *T. delacouri* and southern limestone langurs compared against northern limestone langurs.

Following the methodology proposed by Wang *et al.*^54^, we analyzed the full coding sequence (CDS) and the 1-kb upstream promoter region for each gene. For both comparisons, we calculated the ratio of inter-group fixed differences (*F*_ST_ ≥ 0.95) to intra-group polymorphic loci. Genes were classified as candidate PSGs if their fixed-to-polymorphic loci ratio was significantly higher than the genome-wide average (Pearson’s chi-square test, *P* < 0.01).

### Genetic diversity and inbreeding

To maintain data comparability and analytical validity, we explicitly excluded museum-preserved historical samples and restricting our dataset exclusively to contemporary high-quality samples. Individual heterozygosity was estimated using ANGSD v0.938^124^, with quality filters set at -minQ 20 and -minmapq 30 to exclude low-quality bases and misaligned reads. Fixation coefficient (*F*_ST_) and nucleotide diversity (π) was calculated in 50-kb sliding windows with a 25-kb step size using VCFtools v0.1.15^125^. Comparative genetic diversity statistics for other primates were retrieved from published data^85^. Differences in diversity between groups were assessed using Welch’s two-sample t-tests. Due to insufficient sequencing depth (<20X), five samples (Tcre_686, Tcre_689, Tcre_710, Tbar_15, and Tbar_17) were excluded from both the diversity and ROH analyses. Genome-wide runs of homozygosity (ROH) were identified using PLINK v2.0^126^. We employed a sliding-window approach with the following parameters: --homozyg-kb 100, --homozyg-snp 50, --homozyg-density 50, and --homozyg-window-snp 50. To account for potential genotyping errors while maintaining stringency, windows were permitted a maximum of one heterozygous call (--homozyg-window-het 1) and five missing calls (--homozyg-window-missing 5); segments were considered terminated if these thresholds were exceeded.

### Variant annotation and genetic load assessment

Variants were functionally annotated using SnpEff v4.3^103^. High and moderate impact variants were broadly defined as loss-of-function (LoF) mutations (e.g., stop-gained, frameshifts, exon loss) and missense/splice-altering variants, whereas low-impact variants comprised synonymous and non-disruptive sequence alterations. To assess evolutionary conservation, GERP scores were transferred from the human (hg38) 100-vertebrate multiZ alignment via LiftOver^127,128^. Derived alleles at highly conserved sites (GERP > 4) were defined as strictly deleterious^129,130^. Genetic load was subsequently partitioned into two categories: masked load (calculated as the sum of GERP scores for heterozygous deleterious alleles per called genotype) and realized load (the sum of GERP scores for homozygous deleterious alleles per called site).

To evaluate the purging efficacy of highly deleterious recessive alleles, we compared the frequency of LoF variants within ROH to those in non-ROH regions^33^. To control for baseline homozygosity variations among individuals, raw LoF counts were normalized against synonymous variant rates within their respective genomic regions. The statistical significance of localized LoF depletion inside ROH was evaluated using paired t-tests.

## Acknowledgments

National Natural Science Foundation of China (W2511016, 32470446), State Key Laboratory of Animal Biodiversity Conservation and Integrated Pest Management (SKLA2504), German Research Foundation (RO 3055/7-1) and Sino-German Mobility Programme (M-0084). We express our sincere gratitude to Prof. James Mallet from Harvard University for his invaluable comments, critical revisions, and insightful suggestions on the data analysis and manuscript structure. Furthermore, we extend our sincere appreciation to Adam Ferguson from the Field Museum of Natural History (FMNH), Neil Duncan from the American Museum of Natural History (AMNH), and Pepijn Kamminga from the Naturalis Biodiversity Center for their invaluable assistance in facilitating access to the specimen samples used in this study.

## Author Contributions

M.L. and C.R. conceived and designed the study. J.Q., Q.Z. and L.Z. conducted analyses. J.Q., Y.T., R.P.M., N.V.T., M.D.L., T.N., R.S. and Z.L. conducted the sampling. D.R., Y.S., G.L. and X.L. provided help for data analysis. J.Q., Q.Z., L.Z., X.Z., N.V.T., T.N., Z.L., C.R. and M.L. contributed to the revisions. J.Q., M.L. and C.R. wrote and edited the manuscript.

## Data Availability Statement

The raw sequence data reported in this paper have been deposited in the Genome Sequence Archive in National Genomics Data Center, China National Center for Bioinformation / Beijing Institute of Genomics, Chinese Academy of Sciences (GSA: PRJCA062116) that are publicly accessible at https://ngdc.cncb.ac.cn/gsa.

## Supplementary Information

Figure S1-Figure S40; Table S1-Table S12

## Reference

1 Seehausen, O. Hybridization and adaptive radiation. Trends Ecol Evol 19, 198–207 (2004).

2 Lamichhaney, S. et al. Rapid hybrid speciation in Darwin’s finches. Science 359, 224–227 (2018).

3 Edelman, N. B. et al. Genomic architecture and introgression shape a butterfly radiation. Science 366, 594–599 (2019).

4 Fontaine, M. C. et al. Extensive introgression in a malaria vector species complex revealed by phylogenomics. Science 347, 1258524 (2015).

5 Brawand, D. et al. The genomic substrate for adaptive radiation in African cichlid fish. Nature 513, 375–381 (2014).

6 Steenwyk, J. L., Li, Y., Zhou, X., Shen, X. X. & Rokas, A. Incongruence in the phylogenomics era. Nat. Rev. Genet. 24, 834–850 (2023).

7 Morlon, H. et al. Phylogenetic insights into diversification. Annu. Rev. Ecol. Evol. Syst. 55, 1–21 (2024).

8 Wu, H. et al. Hybrid origin of a primate, the gray snub-nosed monkey. Science 380, eabl4997 (2023).

9 Mallet, J. Hybrid speciation. Nature 446, 279–283 (2007).

10 Schumer, M., Rosenthal, G. G. & Andolfatto, P. How common is homoploid hybrid speciation? Evolution 68, 1553–1560 (2014).

11 Guo, Y. Q. et al. Incomplete lineage sorting shaped mixed traits during a colobine primate radiation. Proc Natl Acad Sci U S A 123, e2524833123 (2026).

12 Harrison, R. G. & Larson, E. L. Hybridization, introgression, and the nature of species boundaries. J. Hered. 105, 795–809 (2014).

13 Marques, D. A., Meier, J. I. & Seehausen, O. A Combinatorial View on Speciation and Adaptive Radiation. Trends in Ecology & Evolution 34, 531–544 (2019).

14 Fitzpatrick, M. J. & Edelsparre, A. H. The genomics of climate change. Science 359, 29–30 (2018).

15 Diez-Del-Molino, D., Sanchez-Barreiro, F., Barnes, I., Gilbert, M. T. P. & Dalen, L. Quantifying Temporal Genomic Erosion in Endangered Species. Trends Ecol Evol 33, 176–185 (2018).

16 Groves, C. P. Primate Taxonomy. (Smithsonian Institution Press, 2001).

17 Roos, C., Boonratana, R., Supriatna, J., Fellowes, J. R. & Schneider, H. An updated taxonomy and conservation status of Asian primates. Asian Primates J. 4, 2–38 (2014).

18 Wang, X. P., Yu, L., Roos, C. & Zhang, Y. P. Phylogenetic relationships among Colobinae monkeys revisited: New insights from analyses of complete mitochondrial genomes and 11 nuclear genes. Molecular Phylogenetics and Evolution 63, 26–33 (2012).

19 Stewart, C. B. & Disotell, T. R. Primate evolution - in and out of Africa. Current Biology 8, R582–R588 (1998).

20 Meyer, D., Rinaldi, I. D., Ramlee, H. & Roos, C. Mitochondrial phylogeny of leaf monkeys (genus Presbytis) reveals a Recent Neo-Sundaic radiation. Molecular Phylogenetics and Evolution 59, 491–498 (2011).

21 Perelman, P., Johnson, W. E., Roos, C. & O’Brien, S. J. A molecular phylogeny of living primates. PLoS genetics 7, e1001342 (2011).

22 Roos, C. et al. Mitogenomic phylogeny of the Asian colobine genus Trachypithecus with special focus on Trachypithecus phayrei and description of a new species. Zoological Research 41, 656–669 (2020).

23 Springer, M. S., Meredith, R. W. & Gatesy, J. Macroevolutionary dynamics and historical biogeography of primate diversification inferred from a species-level phylogeny. PloS one 7, e49521 (2012).

24 Liu, Z., Tan, X. L., Liu, G. J. & Roos, C. Full-length Numt analysis provides evidence for hybridization between the Asian colobine genera Trachypithecus and Semnopithecus. American Journal of Primatology 77, 57–69 (2015).

25 Geissmann, T. et al. A new species of lutung (Trachypithecus) from central Myanmar. Zoological Research 41, 656–669 (2020).

26 Liu, Z., Liu, G. J., Huang, C. M. & Li, M. Living on the rocks: Genomic analysis of limestone langurs provides novel insights into the adaptive evolution in extreme karst environments. Science China Life Sciences 63, 507–519 (2020).

27 Huang, C. M., Zhou, Q. H. & Li, M. The limestone langurs of Vietnam and China: Ecology and conservation. American Journal of Primatology 77, 50–65 (2015).

28 Liu, Z. J. et al. Living on the Rocks: Genomic Analysis of Limestone Langurs Provides Novel Insights into the Adaptive Evolution in Extreme Karst Environments. Genomics Proteomics & Bioinformatics 23 (2025).

29 Wang, X. P., Yu, L., Roos, C. & Zhang, Y. P. Phylogenetic relationships among Colobinae monkeys revisited: new insights from analyses of complete mitochondrial genomes and 11 nuclear genes. Mol. Phylogenet. Evol. 63, 26–33 (2012).

30 Wang, B. et al. Phylogenetic relationships and divergence times of the Asian colobines: Evidence from nuclear mitochondrial pseudogenes (NUMTs). BMC Evolutionary Biology 15, 114 (2015).

31 Iucn. The IUCN Red List of Threatened Species. Version 2024-2. (2024).

32 Theissinger, K. et al. How genomics can help biodiversity conservation. Trends Genet. 39, 545–559 (2023).

33 Dussex, N. et al. Population genomics of the critically endangered kākāpō. Cell Genomics 1 (2021).

34 Jónsson, H. et al. mapDamage2.0: fast approximate Bayesian estimates of ancient DNA damage parameters. Bioinformatics 29, 1682–1684 (2013).

35 Sayyari, E. & Mirarab, S. Fast Coalescent-Based Computation of Local Branch Support from Quartet Frequencies. Molecular biology and evolution 33, 1654–1668 (2016).

36 Martin, S. H. & Van Belleghem, S. M. Exploring Evolutionary Relationships Across the Genome Using Topology Weighting. Genetics 206, 429–438 (2017).

37 Huson, D. H. & Bryant, D. Application of phylogenetic networks in evolutionary studies. Molecular biology and evolution 23, 254–267 (2006).

38 Foley, N. M. et al. An ancient recombination desert is a speciation supergene in placental mammals. Nature 649, 1228–1236 (2026).

39 Voris, H. K. Maps of Pleistocene sea levels in Southeast Asia: shorelines, river systems and time durations. Journal of Biogeography 27, 1153–1167 (2000).

40 Qi, J. et al. Insights into genetic variation and demographic history of sharpnose rays: examinations of three species of Telatrygon (Elasmobranchii, Dasyatidae) from the Indo-West Pacific. Integrative zoology 17, 1063–1077 (2022).

41 Hanebuth, T., Stattegger, K. & Grootes, P. M. Rapid flooding of the Sunda Shelf: A late-glacial sea-level record. Science 288, 1033–1035 (2000).

42 Li, H. & Durbin, R. Inference of human population history from individual whole-genome sequences. Nature 475, 493–496 (2011).

43 Phillips, S. J., Anderson, R. P. & Schapire, R. E. Maximum entropy modeling of species geographic distributions. Ecological Modelling 190, 231–259 (2006).

44 Malinsky, M., Matschiner, M. & Svardal, H. Dsuite -Fast D-statistics and related admixture evidence from VCF files. Molecular Ecology Resources 21, 584–595 (2021).

45 Patterson, N. et al. Ancient admixture in human history. Genetics 192, 1065–1093 (2012).

46 Blischak, P. D., Chifman, J., Wolfe, A. D., Kubatko, L. S. & Posada, D. HyDe: A Python Package for Genome-Scale Hybridization Detection. Systematic Biology 67, 821–829 (2018).

47 Zhang, B.-L. et al. Comparative genomics reveals the hybrid origin of a macaque group. Sci. Adv. 9, eadd3580 (2023).

48 Murrell, B. et al. Gene-Wide Identification of Episodic Selection. Molecular biology and evolution 32, 1365–1371 (2015).

49 Van der Valk, T. et al. The Genome of the Endangered Dryas Monkey Provides New Insights into the Evolutionary History of the Vervets. Molecular biology and evolution 37, 183–194 (2020).

50 Zhang, Y. B. et al. Inferring Historical Introgression with Deep Learning. Systematic Biology 72, 1013–1038 (2023).

51 Arzik, Y. et al. Genome-wide scan of wool production traits in Akkaraman sheep. Genes 14, 713 (2023).

52 Qin, Z., Sun, X., Sun, L., Yu, M. & Jiang, H. Transcriptome sequencing reveals the key genes associated with hair follicle development in Qianhua Mutton Merino. Front. Vet. Sci. 12, 1699868 (2025).

53 Allen, W. L., Stevens, M. & Higham, J. P. Character displacement of Cercopithecini primate visual signals. Nat Commun 5, 4266 (2014).

54 Wang, Z. F. et al. Hybrid speciation via inheritance of alternate alleles of parental isolating genes. Mol. Plant 14, 208–222 (2021).

55 Liu, M. X. et al. SHCBP1L, a conserved protein in mammals, is predominantly expressed in male germ cells and maintains spindle stability during meiosis in testis. Molecular Human Reproduction 20, 463–475 (2014).

56 Ben Khelifa, M., et al. Mutations in DNAH1, which Encodes an Inner Arm Heavy Chain Dynein, Lead to Male Infertility from Multiple Morphological Abnormalities of the Sperm Flagella. American Journal of Human Genetics 94, 95–104 (2014).

57 Balbach, M. et al. Soluble adenylyl cyclase inhibition prevents human sperm function essential for fertilization. Mol. Hum. Reprod. 27, gaab054 (2021).

58 Brito, S. et al. The *Slc45a4* gene regulates pigmentation in a manner distinct from that of the *OCA4* gene *Slc45a2*. J. Invest. Dermatol. 144, 720–722 (2024).

59 Hart, J. C. & Miller, C. T. Sequence-Based Mapping and Genome Editing Reveal Mutations in Stickleback Hps5 Cause Oculocutaneous Albinism and the casper Phenotype. G3-Genes Genomes Genetics 7, 3123–3131 (2017).

60 Zhou, D. et al. Mammalian pigmentation is regulated by a distinct cAMP-dependent mechanism that controls melanosome pH. Sci. Signal. 11, eaau7987 (2018).

61 Zhang, L. et al. Genomic adaptation to small population size and saltwater consumption in the critically endangered Cat Ba langur. Nat Commun 15, 8531 (2024).

62 Dussex, N., Morales, H. E., Grossen, C., Dalén, L. & van Oosterhout, C. Purging and accumulation of genetic load in conservation. Trends Ecol. Evol. 38, 961–969 (2023).

63 Hedrick, P. W. & Garcia-Dorado, A. Understanding Inbreeding Depression, Purging, and Genetic Rescue. Trends Ecol Evol 31, 940–952 (2016).

64 Tan, X. et al. Phylogenomics reveals high levels of incomplete lineage sorting at the ancestral nodes of the macaque radiation. Mol. Biol. Evol. 40, msad229 (2023).

65 Knowles, L. L. Estimating Species Trees: Methods of Phylogenetic Analysis When There Is Incongruence across Genes. Systematic Biology 58, 463–467 (2009).

66 Degnan, J. H. & Rosenberg, N. A. Gene tree discordance, phylogenetic inference and the multispecies coalescent. Trends in Ecology & Evolution 24, 332–340 (2009).

67 Zinner, D., Arnold, M. & Roos, C. The Strange Blood: Natural Hybridization in Primates. Evolutionary Anthropology 20, 96–103 (2011).

68 Handley, L. J., Berset-Brändli, L. & Perrin, N. Disentangling reasons for low Y chromosome variation in the greater white-toothed shrew (*Crocidura russula*). Proc. R. Soc. B 273, 2025–2031 (2006).

69 Goldberg, A., Verdu, P. & Rosenberg, N. A. Autosomal admixture levels are informative about sex bias in admixed populations. Genetics 198, 1209–1229 (2014).

70 Fraïsse, C. & Sachdeva, H. The rates of introgression and barriers to genetic exchange between hybridizing species: sex chromosomes vs autosomes. Genetics 217, iyaa025 (2021).

71 Fontaine, M. C. et al. Extensive introgression in a malaria vector species complex revealed by phylogenomics. Science 347, 1258524 (2015).

72 Martin, S. H. et al. Genome-wide evidence for speciation with gene flow in *Heliconius* butterflies. Genome Res. 23, 1817–1828 (2013).

73 Moore, J. Dispersal, nepotism, and primate social-behavior. International Journal of Primatology 13, 361–378 (1992).

74 Silk, J. B. & Brown, G. R. Local resource competition and local resource enhancement shape primate birth sex ratios. Proceedings of the Royal Society B-Biological Sciences 275, 1761–1765 (2008).

75 Borries, C., Larney, E., Derby, A. M. & Koenig, A. Temporary absence and dispersal in Phayre’s leaf monkeys (Trachypithecus phayrei). Folia Primatologica 75, 27–30 (2004).

76 Osterholz, M., Walter, L. & Roos, C. Phylogenetic position of the langur genera Semnopithecus and Trachypithecus among Asian colobines, and genus affiliations of their species groups. BMC Evol Biol 8, 58 (2008).

77 Karanth, K. P. Primate numts and reticulate evolution of capped and golden leaf monkeys (Primates: Colobinae). Journal of Biosciences 33, 761–770 (2008).

78 Liedigk, R., Thinh, V. N., Nadler, T., Walter, L. & Roos, C. Evolutionary history and phylogenetic position of the Indochinese grey langur (Trachypithecus crepusculus). Vietnamese Journal of Primatology 3, 1–8 (2009).

79 Cheng, S. R., Flouri, T., Zhu, T. Q. & Yang, Z. H. The impact of incomplete taxon sampling on inference of gene flow by Bayesian and summary methods using genomic sequence data. Syst. Biol. 75, syag023 (2026).

80 Pang, X. X. & Zhang, D. Y. Detection of Ghost Introgression Requires Exploiting Topological and Branch Length Information. Systematic Biology 73, 207–222 (2024).

81 Tricou, T., Tannier, E. & De Vienne, D. M. Ghost lineages highly influence the interpretation of introgression tests. Syst. Biol. 71, 1147–1158 (2022).

82 Liu, Z. et al. Genomic Mechanisms of Physiological and Morphological Adaptations of Limestone Langurs to Karst Habitats. Molecular biology and evolution 37, 952–968 (2020).

83 Barrett, R. D. H. & Schluter, D. Adaptation from standing genetic variation. Trends in Ecology & Evolution 23, 38–44 (2008).

84 Tung, J. & Barreiro, L. B. The contribution of admixture to primate evolution. Current Opinion in Genetics & Development 47, 61–68 (2017).

85 Kuderna, L. F. K. et al. A global catalog of whole-genome diversity from 233 primate species. Science 380, 906–913 (2023).

86 Willoughby, J. R. et al. The impacts of inbreeding, drift and selection on genetic diversity in captive breeding populations. Molecular ecology 24, 98–110 (2015).

87 Galla, S. J. et al. A comparison of pedigree, genetic and genomic estimates of relatedness for informing pairing decisions in two critically endangered birds: Implications for conservation breeding programmes worldwide. Evolutionary Applications 13, 991–1008 (2020).

88 Díez-del-Molino, D., Sánchez-Barreiro, F., Barnes, I., Gilbert, M. T. P. & Dalén, L. Quantifying Temporal Genomic Erosion in Endangered Species. Trends in Ecology & Evolution 33, 176–185 (2018).

89 Formenti, G. et al. The era of reference genomes in conservation genomics. Trends Ecol Evol 37, 197–202 (2022).

90 Gargiulo, R., Budde, K. B. & Heuertz, M. Mind the lag: understanding genetic extinction debt for conservation. Trends in Ecology & Evolution 40, 228–237 (2025).

91 Rohland, N., Siedel, H. & Hofreiter, M. Nondestructive DNA extraction method for mitochondrial DNA analyses of museum specimens. Biotechniques 36, 814–821 (2004).

92 Dabney, J. et al. Complete mitochondrial genome sequence of a Middle Pleistocene cave bear reconstructed from ultrashort DNA fragments. Proceedings of the National Academy of Sciences of the United States of America 110, 15758–15763 (2013).

93 Caroe, C. et al. Single-tube library preparation for degraded DNA. Methods in Ecology and Evolution 9, 410–419 (2018).

94 Dierckxsens, N., Mardulyn, P. & Smits, G. NOVOPlasty: *de novo* assembly of organelle genomes from whole genome data. Nucleic Acids Res. 45, e43 (2017).

95 Meng, G., Li, Y., Yang, C. & Liu, S. MitoZ: a toolkit for animal mitochondrial genome assembly, annotation and visualization. Nucleic Acids Res. 47, e63 (2019).

96 Yamada, K. D., Tomii, K. & Katoh, K. Application of the MAFFT sequence alignment program to large data-reexamination of the usefulness of chained guide trees. Bioinformatics 32, 3246–3251 (2016).

97 Castresana, J. Selection of conserved blocks from multiple alignments for their use in phylogenetic analysis. Molecular Biology & Evolution 17, 540–552 (2000).

98 Stamatakis, A. RAxML version 8: a tool for phylogenetic analysis and post-analysis of large phylogenies. Bioinformatics 30, 1312–1313 (2014).

99 Chen, S. F. Ultrafast one-pass FASTQ data preprocessing, quality control, and deduplication using fastp. iMeta 2, e107 (2023).

100 Li, H. & Durbin, R. Fast and accurate short read alignment with Burrows-Wheeler transform. Bioinformatics 25, 1754–1760 (2009).

101 Li, H. et al. The Sequence Alignment/Map format and SAMtools. Bioinformatics 25, 2078–2079 (2009).

102 McKenna, A. et al. The Genome Analysis Toolkit: A MapReduce framework for analyzing next-generation DNA sequencing data. Genome Research 20, 1297–1303 (2010).

103 Cingolani, P. et al. A program for annotating and predicting the effects of single nucleotide polymorphisms, SnpEff: SNPs in the genome of *Drosophila melanogaster* strain w(1118); iso-2; iso-3. Fly 6, 80–92 (2012).

104 Manichaikul, A. et al. Robust relationship inference in genome-wide association studies. Bioinformatics 26, 2867–2873 (2010).

105 Bouckaert, R. R. DensiTree: making sense of sets of phylogenetic trees. Bioinformatics 26, 1372–1373 (2010).

106 Li, H. Minimap2: pairwise alignment for nucleotide sequences. Bioinformatics 34, 3094–3100 (2018).

107 Danecek, P. et al. Twelve years of SAMtools and BCFtools. Gigascience 10 (2021).

108 Patterson, N., Price, A. L. & Reich, D. Population structure and eigenanalysis. PLoS genetics 2, 2074–2093 (2006).

109 Alexander, D. H., Novembre, J. & Lange, K. Fast model-based estimation of ancestry in unrelated individuals. Genome Research 19, 1655–1664 (2009).

110 Emms, D. M. & Kelly, S. OrthoFinder: phylogenetic orthology inference for comparative genomics. Genome Biol. 20, 238 (2019).

111 Loytynoja, A. Phylogeny-aware alignment with PRANK. Methods in molecular biology (Clifton, N.J.) 1079, 155–170 (2014).

112 Kumar, S., Stecher, G. & Tamura, K. MEGA7: Molecular Evolutionary Genetics Analysis Version 7.0 for bigger datasets. Mol. Biol. Evol. 33, 1870–1874 (2016).

113 Yang, Z. H. PAML 4: Phylogenetic analysis by maximum likelihood. Molecular biology and evolution 24, 1586–1591 (2007).

114 Benton, M. J., Donoghue, P. C. J. & Asher, R. J. Constraints on the phylogeny of placental mammals. Palaeontology 58, 1–45 (2015).

115 Zalmout, I. S., Sanders, W. J. & MacLatchy, L. M. New Oligocene primate from Saudi Arabia and the divergence of apes and Old World monkeys. Nature 466, 360–364 (2010).

116 de Vries, D. & Beck, R. M. D. Twenty-five well-justified fossil calibrations for primate divergences. *Palaeontol*. Electron. 26, a8 (2023).

117 Hilgers, L. et al. Avoidable false PSMC population size peaks occur across numerous studies. Current Biology 35 (2025).

118 Zizka, A. et al. CoordinateCleaner: Standardized cleaning of occurrence records from biological collection databases. Methods in Ecology and Evolution 10, 744–751 (2019).

119 Hijmans, R. J., Cameron, S. E., Parra, J. L., Jones, P. G. & Jarvis, A. Very high resolution interpolated climate surfaces for global land areas. International Journal of Climatology 25, 1965–1978 (2005).

120 Hijmans, R. J. raster: Geographic data analysis and modeling. R package version 2.3-12 (2014).

121 Schmitt, S. et al. ssdm: An r package to predict distribution of species richness and composition based on stacked species distribution models. Methods in Ecology and Evolution 8, 1795–1803 (2017).

122 Martin, S. H., Davey, J. W. & Jiggins, C. D. Evaluating the Use of ABBA-BABA Statistics to Locate Introgressed Loci. Molecular biology and evolution 32, 244–257 (2015).

123 Yu, G., Wang, L.-G., Han, Y. & He, Q.-Y. clusterProfiler: an R Package for Comparing Biological Themes Among Gene Clusters. Omics-a Journal of Integrative Biology 16, 284–287 (2012).

124 Korneliussen, T. S., Albrechtsen, A. & Nielsen, R. ANGSD: analysis of next generation sequencing data. BMC Bioinformatics 15, 356 (2014).

125 Danecek, P. et al. The variant call format and VCFtools. Bioinformatics 27, 2156–2158 (2011).

126 Chang, C. C. et al. Second-generation PLINK: rising to the challenge of larger and richer datasets. GigaScience 4, 7 (2015).

127 Rosenbloom, K. R. et al. The UCSC Genome Browser database: 2015 update. Nucleic Acids Research 43, D670–D681 (2015).

128 Kuhn, R. M., Haussler, D. & Kent, W. J. The UCSC genome browser and associated tools. Briefings in Bioinformatics 14, 144–161 (2013).

129 Smeds, L. & Ellegren, H. From high masked to high realized genetic load in inbred Scandinavian wolves. Molecular ecology 32, 1567–1580 (2023).

130 Huber, C. D., Kim, B. Y. & Lohmueller, K. E. Population genetic models of GERP scores suggest pervasive turnover of constrained sites across mammalian evolution. PLoS Genet. 16, e1008827 (2020).

