## supplementary file for "Ancient admixture catalyzes homoploid hybrid speciation and intense genomic erosion in an Asian langur genus"

**Supplementary Figures**

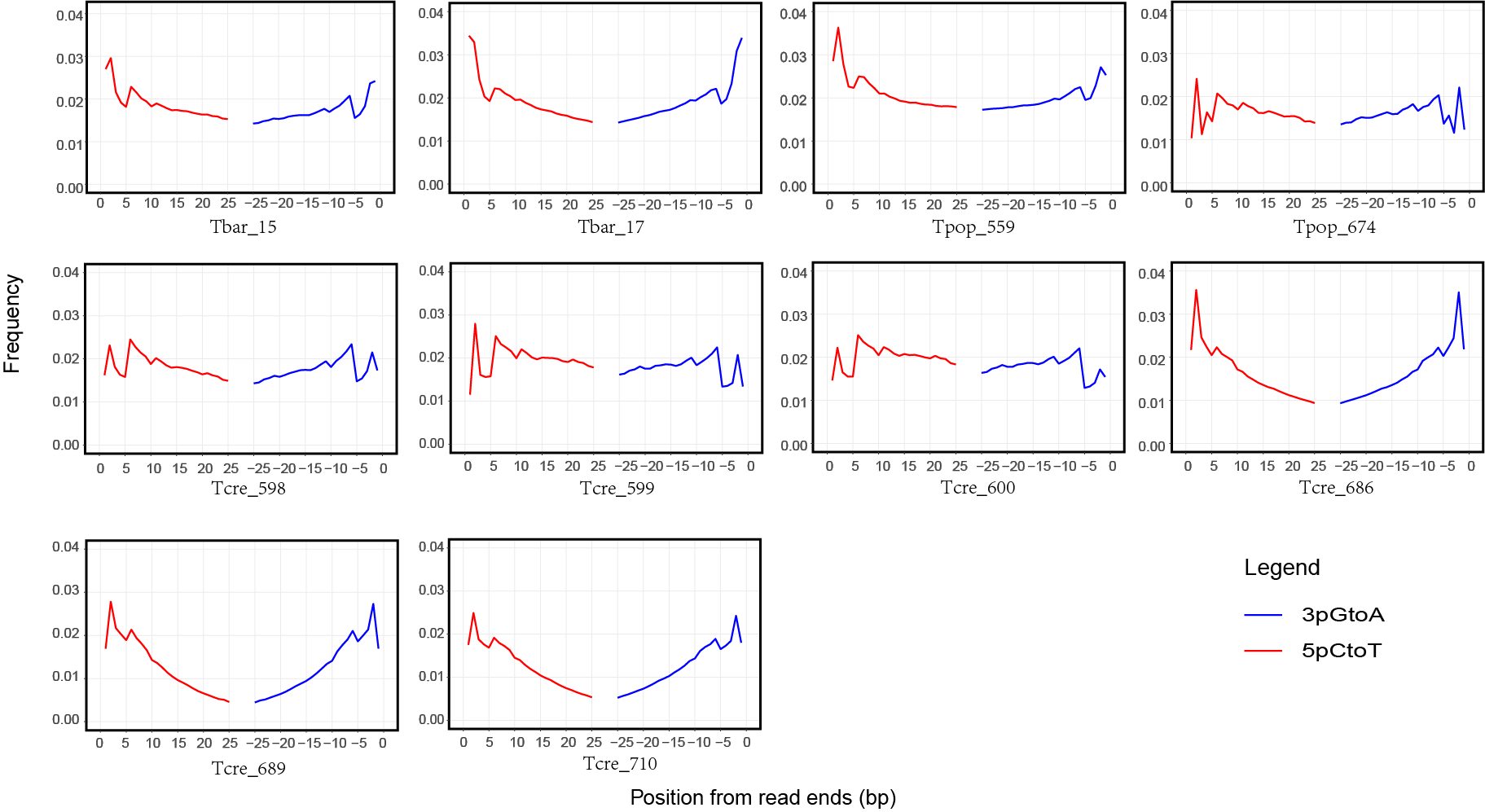

**Supplementary Figure 1 | DNA damage profiles of *Trachypithecus* museum specimens.** Nucleotide substitution frequencies are plotted as a function of distance from the 5′ (red lines, C-to-T transitions) and 3′ (blue lines, G-to-A transitions) ends of the sequencing reads. These patterns, generated using mapDamage v2.0, illustrate the post-mortem DNA deamination characteristic of historical samples. Samples such as Tbar_15 and Tcre_689 exhibit typical terminal spikes in deamination, confirming the authenticity of the museum-derived DNA. Conversely, samples Tcre_598, Tcre_599, Tcre_600, and Tpop_674 lack these diagnostic damage signatures, suggesting either exceptional DNA preservation or potential modern DNA contamination.

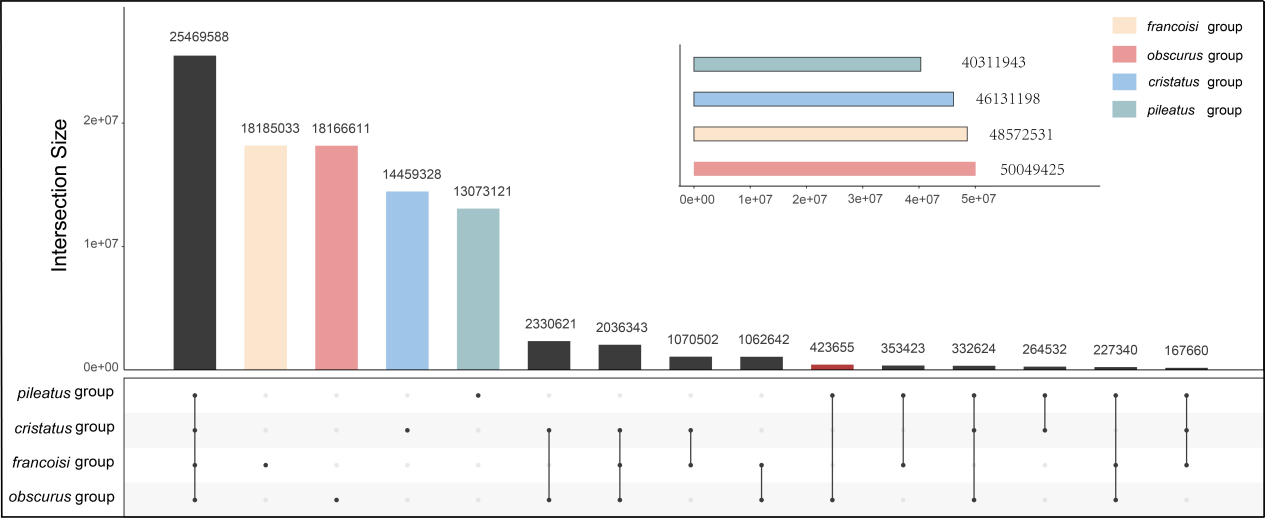

**Supplementary Figure 2 | Distribution of shared and unique SNVs across *Trachypithecus* species groups.** An UpSet plot illustrating the intersection of 156,262,744 high-quality SNVs identified across 55 individuals. The four species groups analyzed are the *pileatus* group (green), *cristatus* group (blue), *francoisi* group (yellow), and *obscurus* group (red). The horizontal bar chart (top right) represents the total SNV set size identified within each species group. The vertical bar chart (main) indicates the size of each intersection; single-colored dots denote SNVs unique to a specific group, while connected dots (black) signify SNVs shared across multiple groups. The largest intersection (black bar on the far left) highlights variants shared across all four groups, reflecting core genomic variation within the genus *Trachypithecus*.

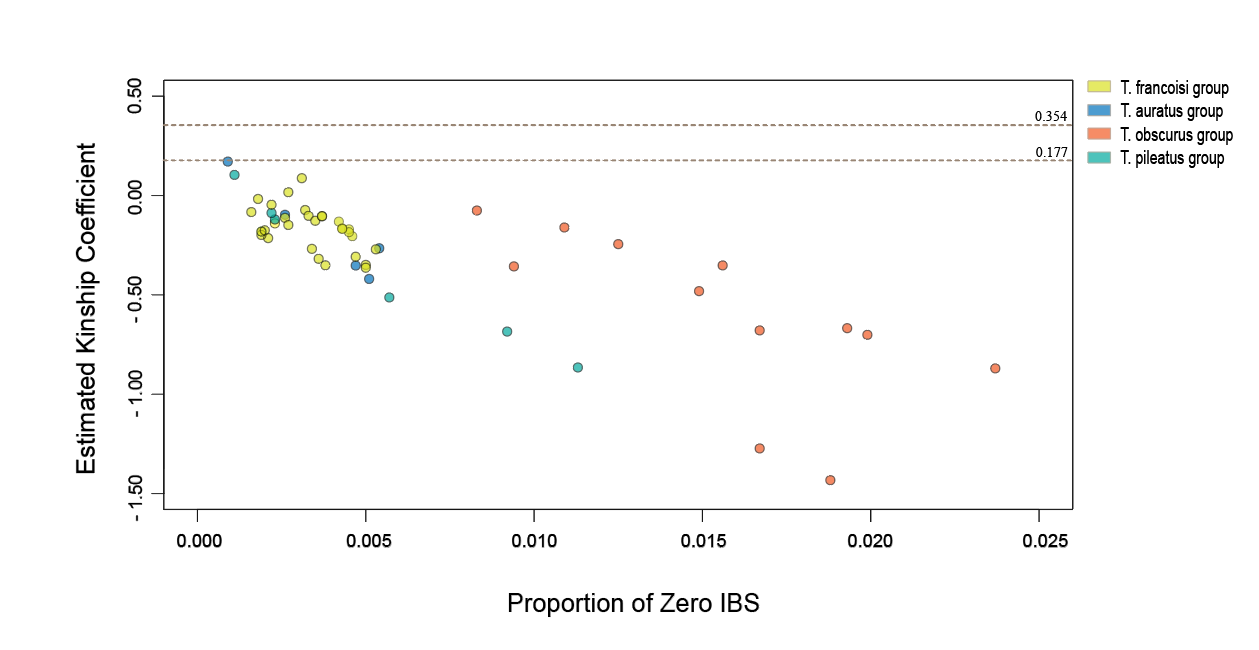

**Supplementary Figure 3 | Pairwise kinship inference among *Trachypithecus* samples using KING robust.** Scatter plot of estimated robust kinship coefficients (y-axis) against the proportion of Zero Identity By State (IBS) loci (x-axis) for all pairwise sample comparisons in the dataset. Data points are color-coded by species group as indicated in the legend: *francoisi* group (yellow), *cristatus* group (blue), *obscurus* group (orange), and *pileatus* group (green). The horizontal dashed lines represent standard KING thresholds for relatedness: 0.354 (first-degree relatives, e.g., parent-offspring or full siblings) and 0.177 (second-degree relatives).

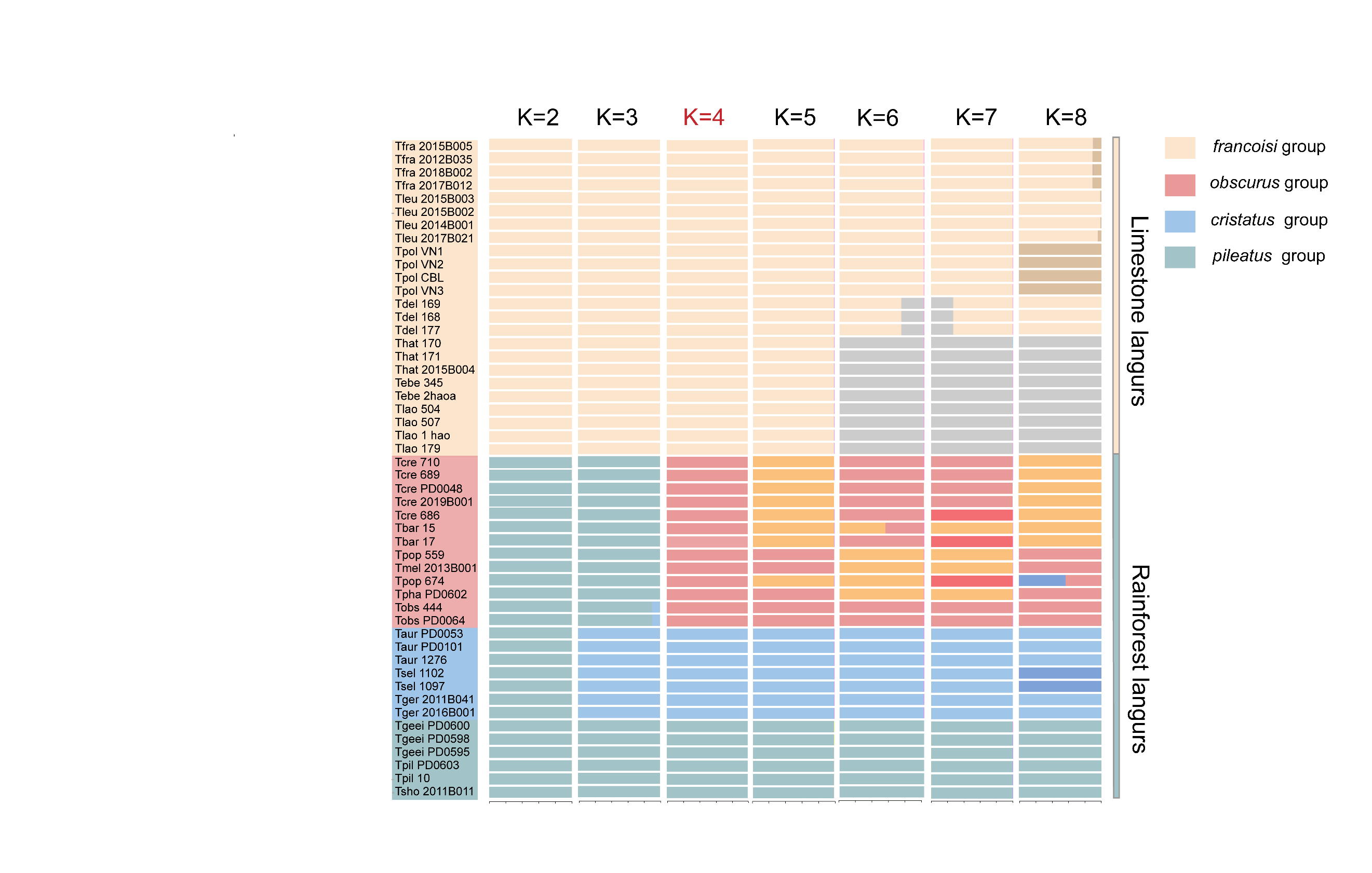

**Supplementary Figure 4 | Population clustering of *Trachypithecus* species groups.** In the ADMIXTURE plots, each vertical bar represents an individual, with colors indicating the estimated proportion of ancestry from each of the K clusters. At K=2, the analysis resolves the primary ecological split between the limestone langurs (cyan cluster) and the rainforest langurs (red cluster). At K=4, four distinct genetic components emerge, corresponding to the four species groups (*francoisi*, *obscurus*, *cristatus*, and *pileatus* groups). The vertical bars on the far right denote the two major ecological guilds.

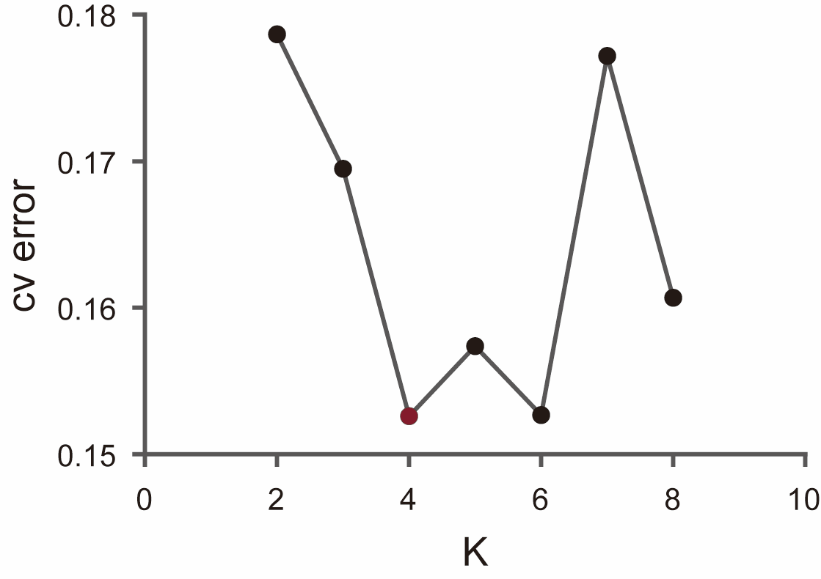

**Supplementary Figure 5 | ADMIXTURE cross-validation (CV) error plot.** The plot shows the CV error for K values ranging from 2 to 8. Lower CV error values indicate higher statistical support for the number of ancestral clusters. The local minima at **K=4** and **K=6** suggest these are the most robust population structure models for the *Trachypithecus* dataset.

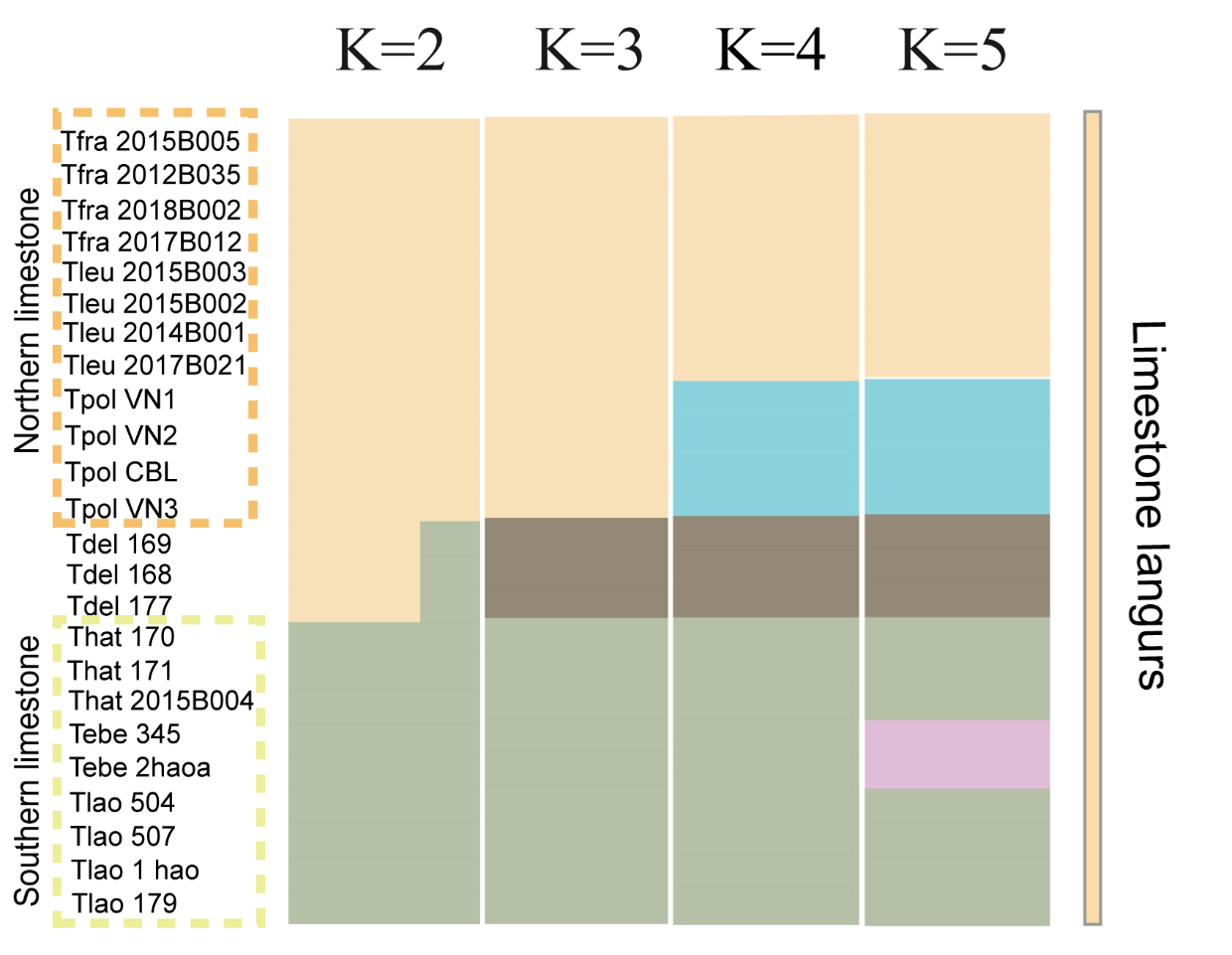

**Supplementary Figure 6 | Population genetic structure and ancestral components of the *francoisi* group.** Model-based clustering analysis using ADMIXTURE for 24 individuals representing all seven species of the *francoisi* group, with the number of ancestral clusters (K) ranging from 2 to 5. Each horizontal bar represents an individual, with colors indicating the estimated proportion of ancestry from each K cluster. The individuals are organized by geographic distribution (northern vs. southern limestone langurs, *T. delacouri* geographically between these two groups) as indicated on the left. At **K=2**, a primary genetic split is observed between northern populations (beige) and southern populations (green), with *T. delacouri* displaying a substantial mosaic of both northern and southern ancestry. At **K=3**, *T. delacouri* (Tdel) emerges as a distinct genetic cluster (brown), while higher values of K (K=4, 5) further differentiate specific lineages, such as the separation of *T. poliocephalus* (Tpol, cyan) and internal structure within the southern clade.

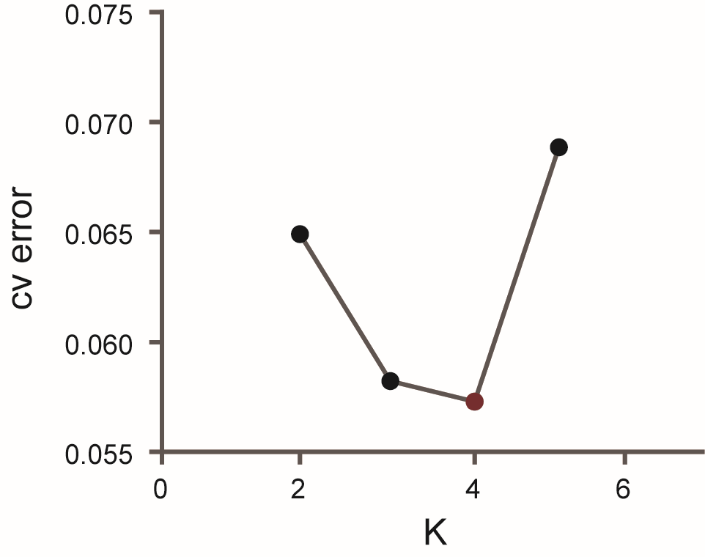

**Supplementary Figure 7 | ADMIXTURE cross-validation (CV) error plot.** The plot shows the CV error for K values ranging from 2 to 5. Lower CV error values indicate higher statistical support for the number of ancestral clusters. The global minimum at **K=4** suggests this is the most robust population structure model for the *francoisi* group dataset.

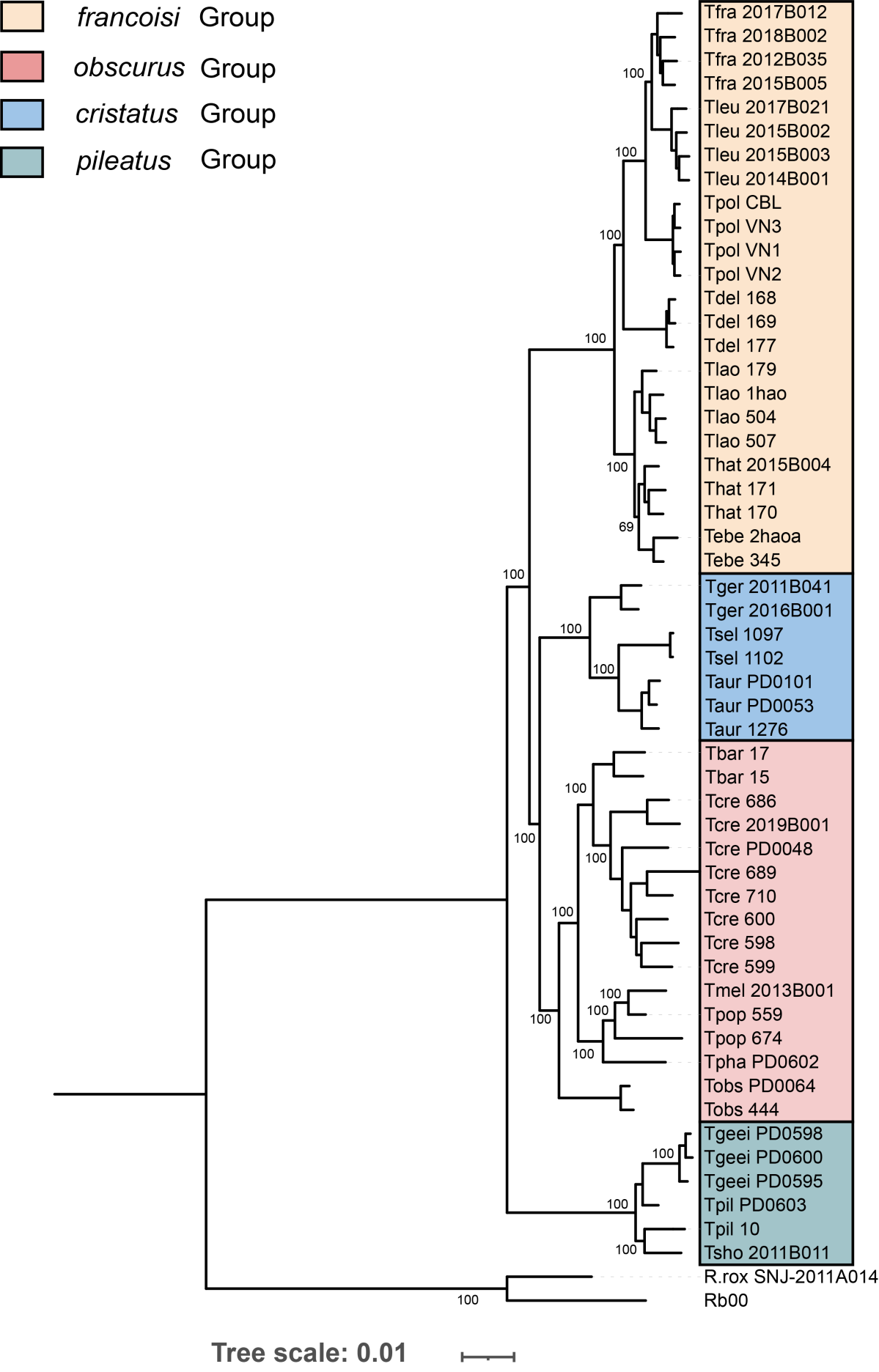

**Supplementary Figure 8 | Maximum likelihood (ML) phylogeny of the genus *Trachypithecus* based on autosomal SNVs.** The phylogenetic tree was reconstructed using RAxML based on a concatenated dataset of autosomal SNVs from 53 individuals, representing 19 species. Bootstrap support values are indicated at internal nodes. The analysis resolves the *pileatus* group (greenish-blue clade) as the basal lineage of the genus. The *francoisi* group (beige clade) is positioned as a sister group to the common ancestor of the *obscurus* and *cristatus* groups (blue and red clades, respectively). Within the *francoisi* group, *T. delacouri* (Tdel) is phylogenetically intermediate between the northern and southern limestone langur clades, consistent with its geographic distribution. Two *Rhinopithecus* individuals (*R. roxellana* and *R. bieti*) were used as outgroups to root the tree. Tree scale represents 0.01 substitutions per site.

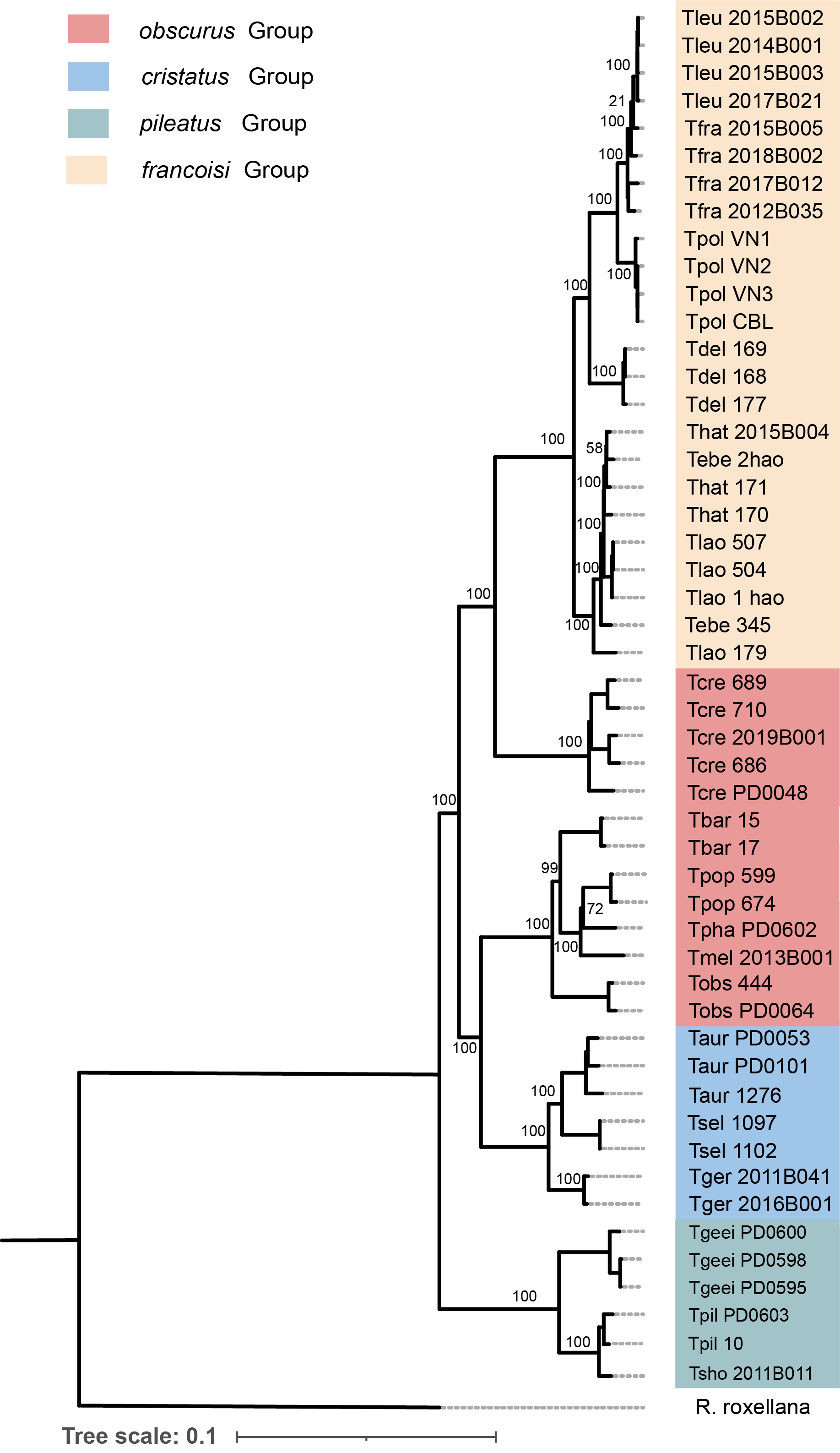

**Supplementary Figure 9 | Maximum likelihood (ML) phylogeny of the genus *Trachypithecus* based on mitochondrial genomes.** The phylogenetic tree was reconstructed using complete mitochondrial genomes (mitogenomes) assembled *de novo* from 50 individuals. To minimize alignment artifacts and ensure phylogenetic accuracy, highly variable regions (e.g., the control region) were excluded prior to analysis. Bootstrap support values are indicated at internal nodes. This maternal phylogeny resolves the *pileatus* group (teal clade) as the basal lineage within the genus. The *francoisi* group (yellow clade) is recovered as a monophyletic lineage, within which *T. delacouri* (Tdel) is nested. The tree reveals the maternal relationships between the species groups, providing a complementary perspective to the nuclear genomic data. *Rhinopithecus roxellana* (R. rox) was used as the outgroup to root the tree. Tree scale represents 0.1 substitutions per site, reflecting the higher evolutionary rate of the mitochondrial genome compared to nuclear DNA.

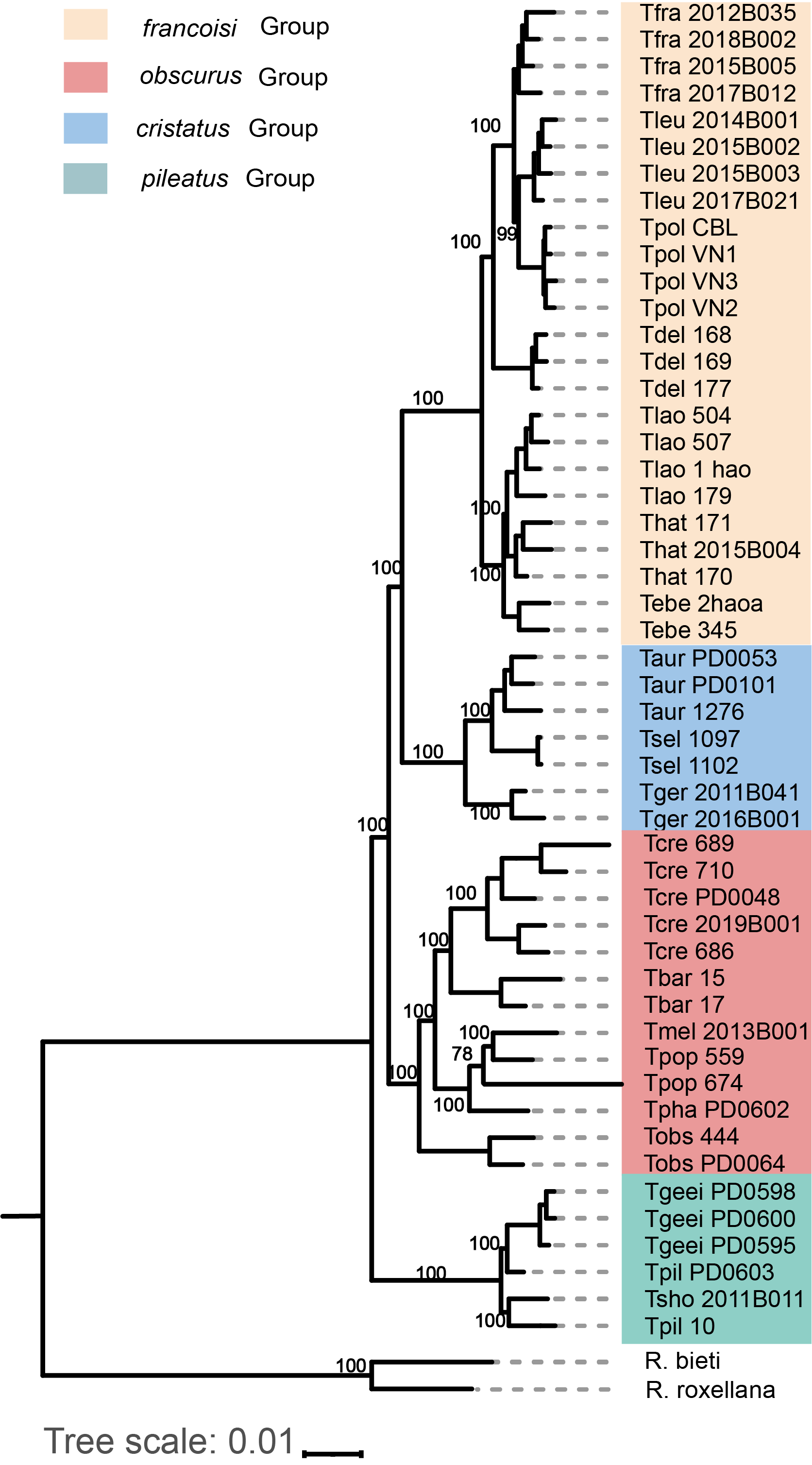

**Supplementary Figure 10 | Maximum likelihood (ML) phylogeny of the genus *Trachypithecus* based on X-linked SNVs.** The phylogenetic tree was reconstructed based on X-linked SNVs from 50 individuals, covering 19 species. Bootstrap support values are indicated at internal nodes. The topology derived from X-chromosomal data supported an alternative topology (T2) (Fig. 2B): while the *pileatus* group retained basal, the *obscurus* group was resolved as the sister lineage to the common ancestor of the *francoisi* and *cristatus* groups. Within the *francoisi* group, *T. delacouri* remains phylogenetically positioned intermediate between the northern and southern limestone langur clades. Two *Rhinopithecus* individuals (*R. roxellana* and *R. bieti*) were used as outgroups to root the tree. Tree scale represents 0.01 substitutions per site.

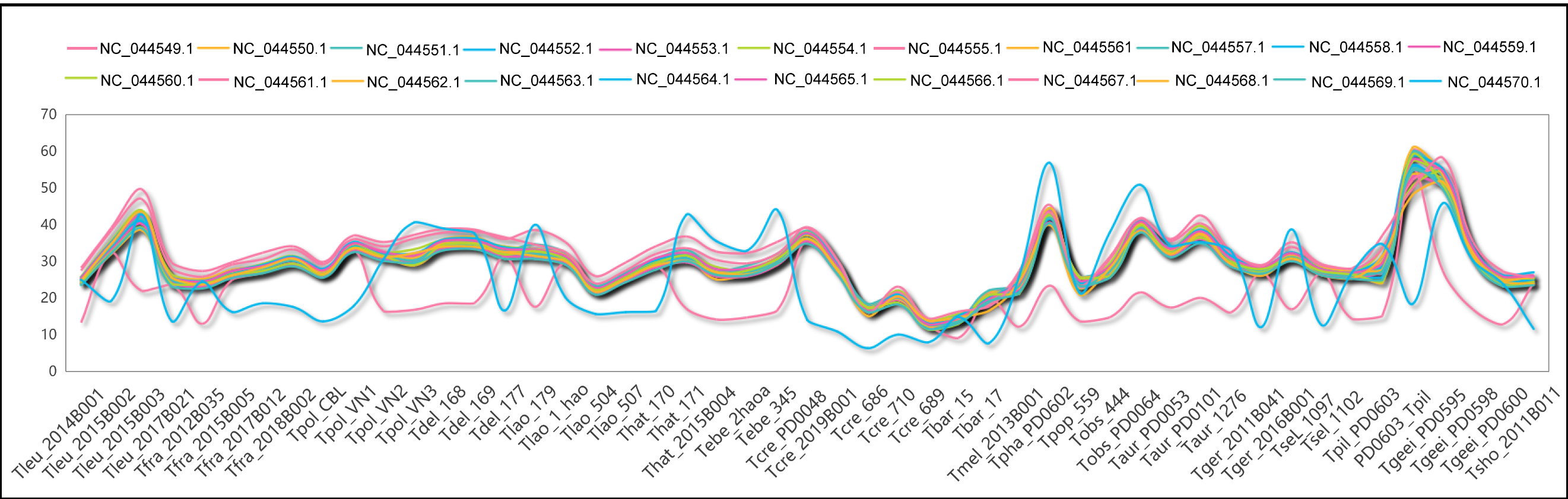

**Supplementary Figure 11 | Sex identification of *Trachypithecus* individuals based on chromosomal sequencing depth.** The plot displays the mean sequencing depth for each chromosome [autosomes NC_044549.1–NC_044554.1, NC_044556.1–NC_044569.1, NC_044555.1(chrX), and NC_044570.1(chrY)] across the 49 individuals in the dataset. Sex was determined by evaluating the normalized coverage of sex chromosomes relative to autosomes: males (XY) are characterized by an X-chromosome depth (pink line) approximately half that of the autosomal depth, accompanied by significant coverage of the Y-chromosome (light blue line). In contrast, females (XX) exhibit X-chromosome depth comparable to autosomal depth and negligible Y-chromosome coverage.

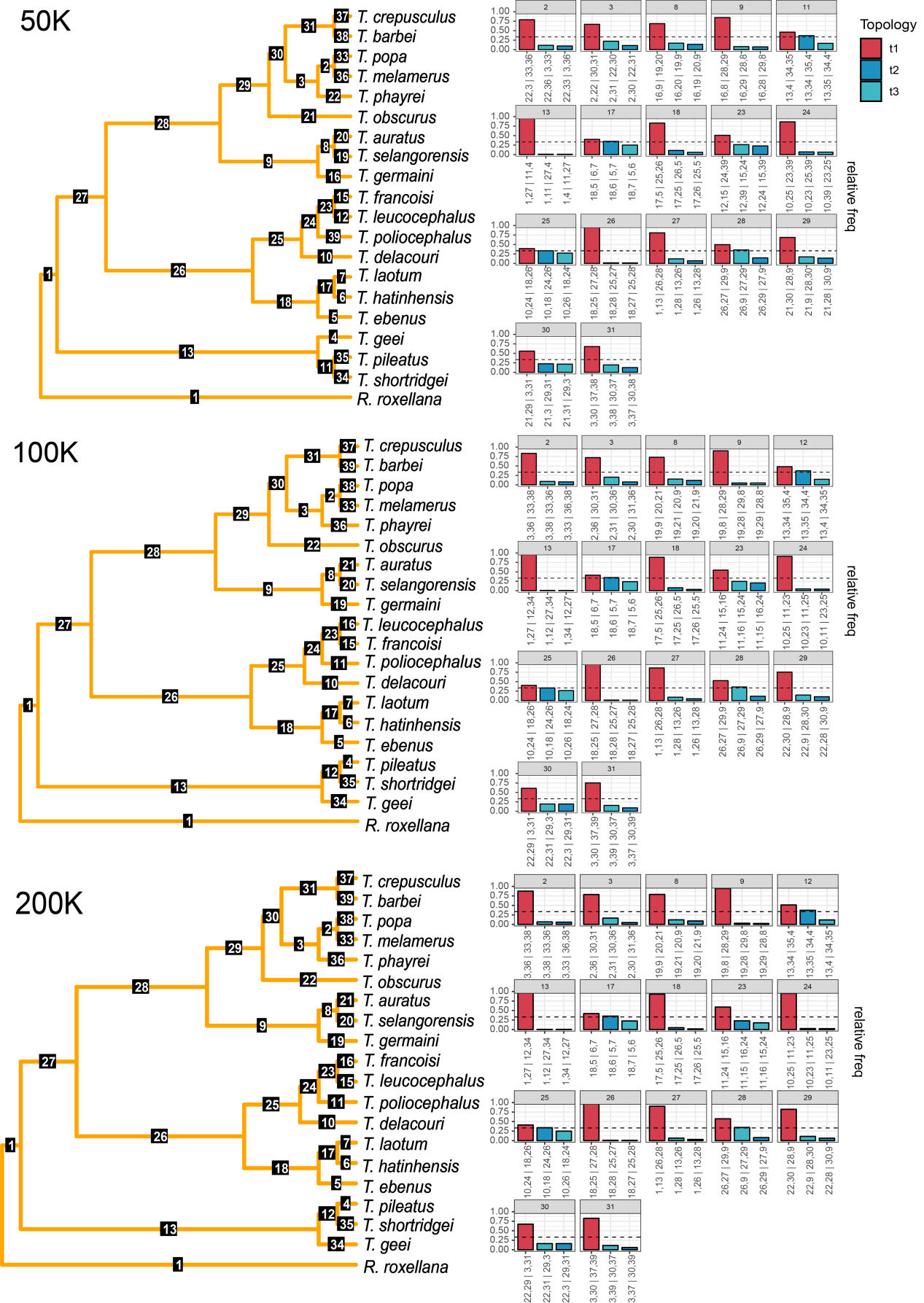

**Supplementary Figure 12 | Species tree reconstruction and genealogical discordance across different genomic window sizes.** Species trees were inferred using ASTRAL based on individual gene trees reconstructed from non-overlapping genomic windows of 50 kb, 100 kb, and 200 kb. For each analysis, the species tree topology is shown on the left with internal nodes numbered. The bar plots on the right display the relative frequencies of the three possible quartet topologies (T1, T2, and T3) for each corresponding internal branch. T1 (red) represents the primary topology consistent with the species tree, while T2 and T3 (blue) represent alternative topologies. These quartet frequencies illustrate the level of genealogical discordance (e.g., due to incomplete lineage sorting or introgression) across the *Trachypithecus* genome. The results show high topological consistency across all three window sizes, confirming the robustness of the inferred phylogenetic relationships within the genus.

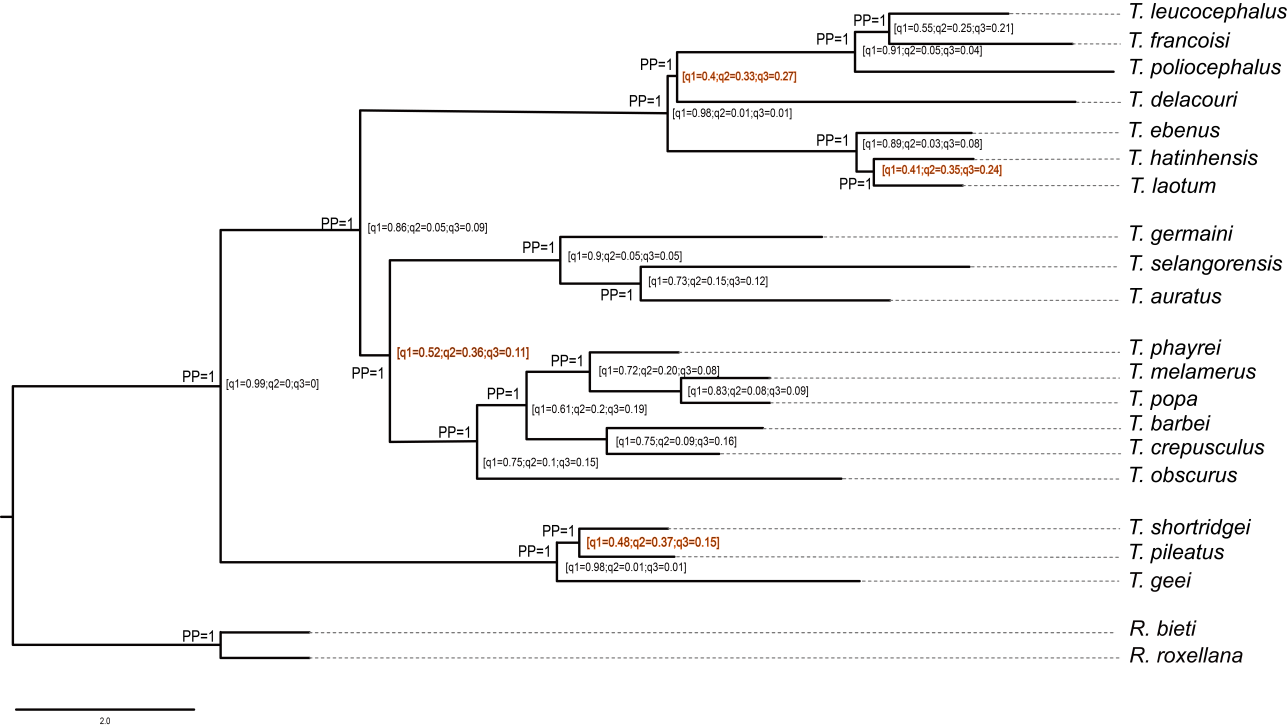

**Supplementary Figure 13 | Multi-species coalescent (MSC)-based species tree of the genus *Trachypithecus*.** The species tree was inferred using ASTRAL based on gene trees reconstructed from genome fragments. Support values at the nodes represent local posterior probabilities (PP) and ASTRAL quartet scores (q1:q2:q3). The quartet scores indicate the relative frequencies of the primary topology (q1) and the two alternative quartet topologies (q2 and q3). Branches labeled with red text highlight specific nodes with notable genealogical discordance (e.g., lower q1 values), suggesting potential historical gene flow or incomplete lineage sorting (ILS) within these lineages. The tree is rooted with *Rhinopithecus roxellana* and *R. bieti*. The scale bar represents 2.0 coalescent units.

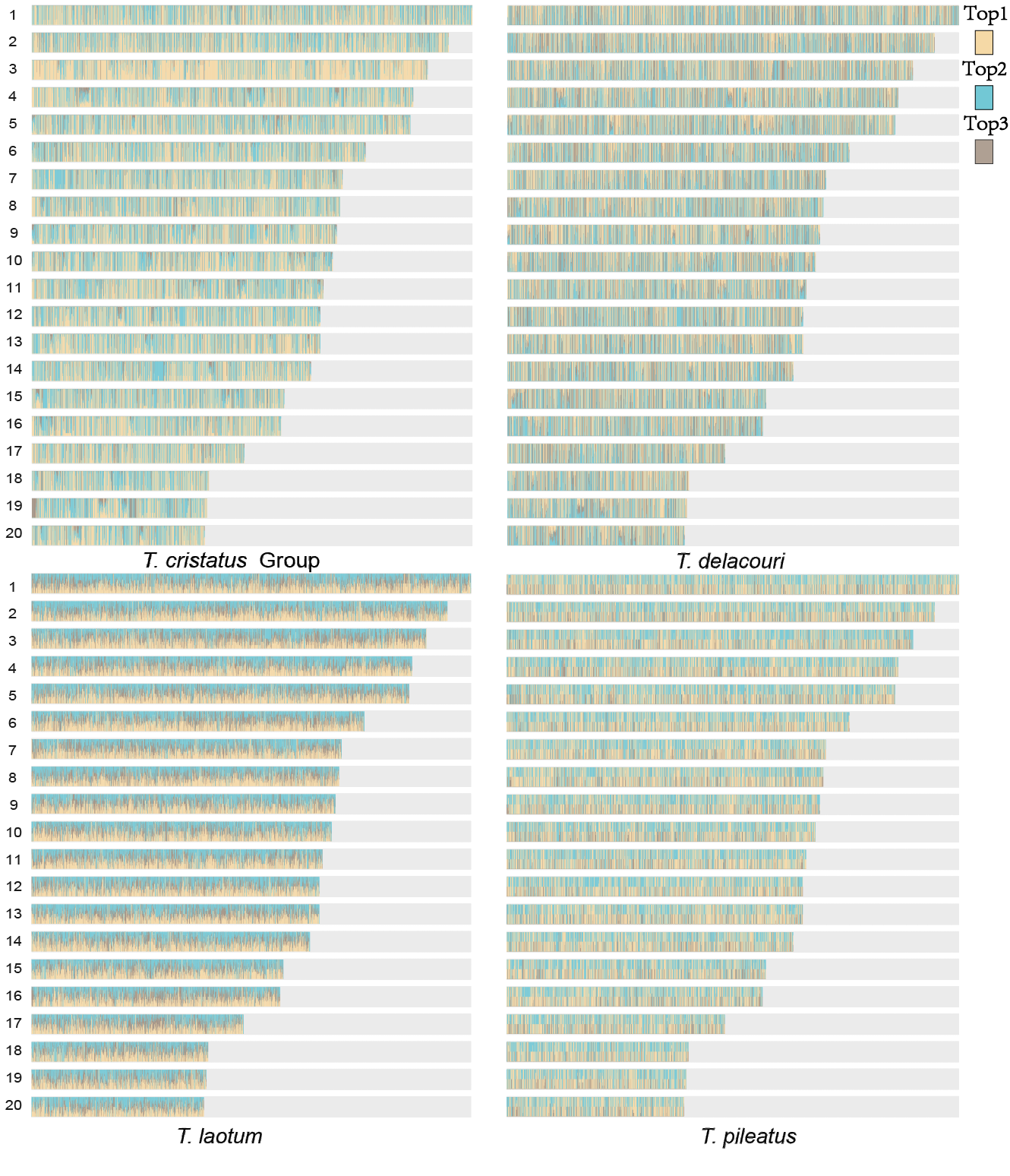

**Supplementary Figure 14 | Genome-wide topology weighting (Twisst) across four focal *Trachypithecus* lineages.** The plots visualize the weighting of three alternative phylogenetic topologies across the 20 autosomes (chr1–20) using Twisst. Each of the four panels corresponds to a lineage or node exhibiting significant phylogenetic discordance: the *T. cristatus* group (top-left), *T. delacouri* (top-right), *T. laotum* (bottom-left), and *T. pileatus* (bottom-right). For each panel, horizontal bars represent individual chromosomes, with colors (beige, cyan, and brown) indicating the relative weights of the three possible quartet topologies at each genomic position. The highly mosaic and fluctuating patterns across the genome illustrate the extent of genealogical discordance, likely resulting from incomplete lineage sorting (ILS) or historical gene flow events.

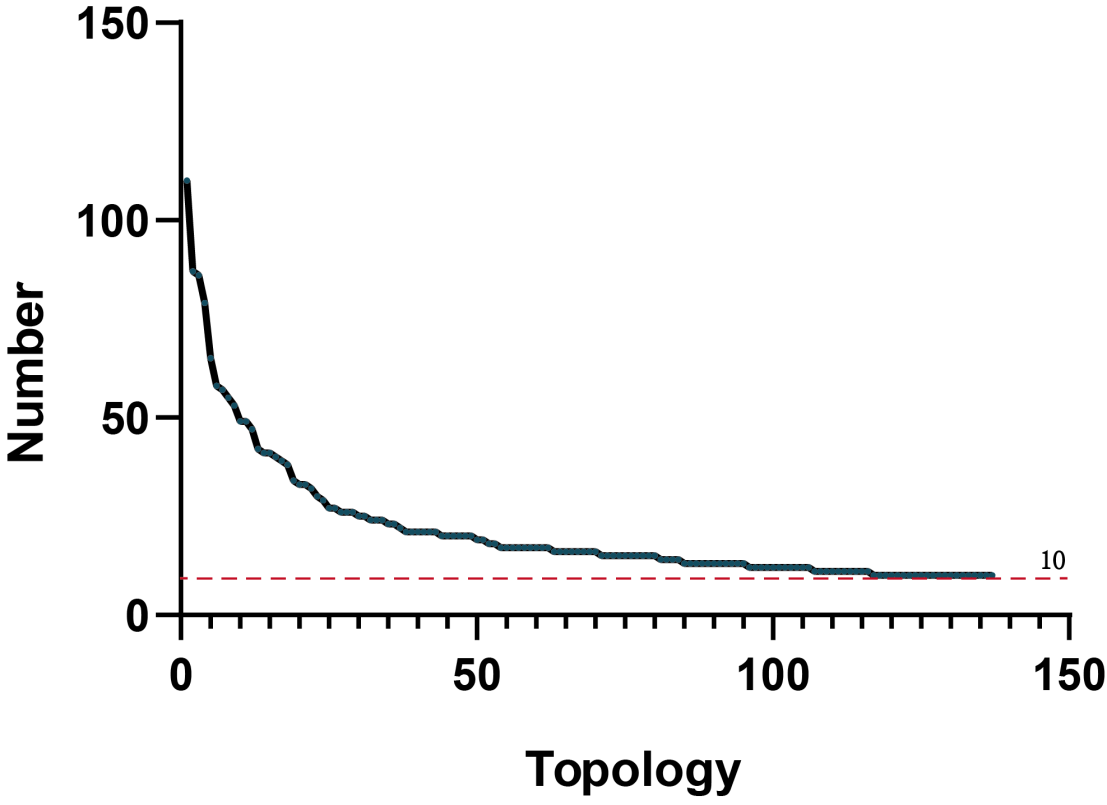

**Supplementary Figure 15 | Frequency distribution of unique gene tree topologies.** The plot illustrates the distribution of unique topologies identified from genomic window-based gene trees (e.g., 200 kb windows) using phybin. The x-axis represents unique topologies ranked by their occurrence frequency, and the y-axis indicates the number of gene trees supporting each specific topology. The red dashed line denotes a frequency threshold of 10; a total of 138 unique topologies are supported by at least 10 gene trees, representing the most common phylogenetic signals in the dataset.

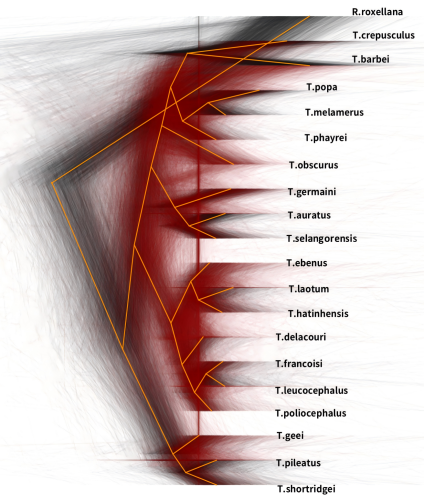

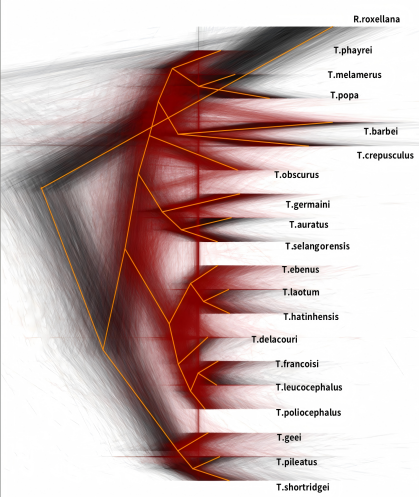

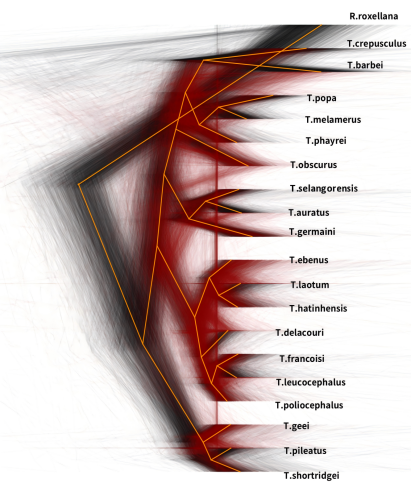

50k	100k	200k

**Supplementary Figure 16 | Cloudograms illustrating pervasive phylogenetic discordance across the *Trachypithecus* genome.** DensiTree visualizations of maximum-likelihood gene trees inferred from non-overlapping genomic windows of 50 kb (left), 100 kb (middle), and 200 kb (right). The dense, overlapping network of tree topologies (background cloud) highlights the extensive incomplete lineage sorting (ILS) and reticulated evolution characterizing the rapid radiation of the genus. The solid orange line superimposed on the cloudograms represents the inferred species tree topology. *Rhinopithecus roxellana* is included as the outgroup.

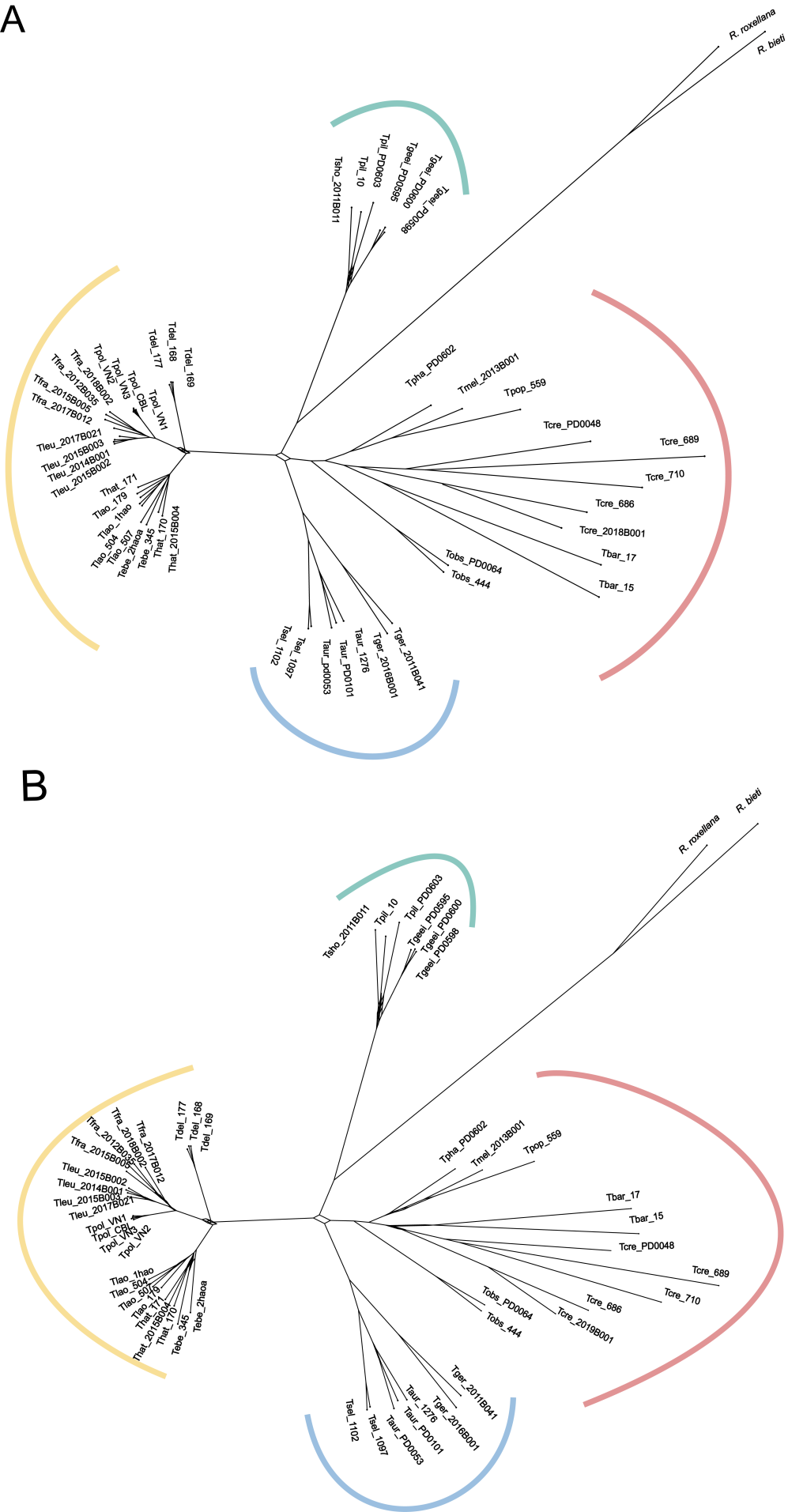

**Supplementary Figure 17 | SplitsTree consensus networks of the genus *Trachypithecus*.** Networks summarize the topological variations across 13,960 genomic window trees (200 kb each). The branch structures depict major splits and alternative topologies among taxa. **A**, Consensus network with edges displayed only if the split is present in at least 20% of the gene trees. **B**, Consensus network with a lower threshold of 10%, displaying deeper levels of incomplete lineage sorting and reticulated evolutionary events. Species groups are highlighted by colored arcs.

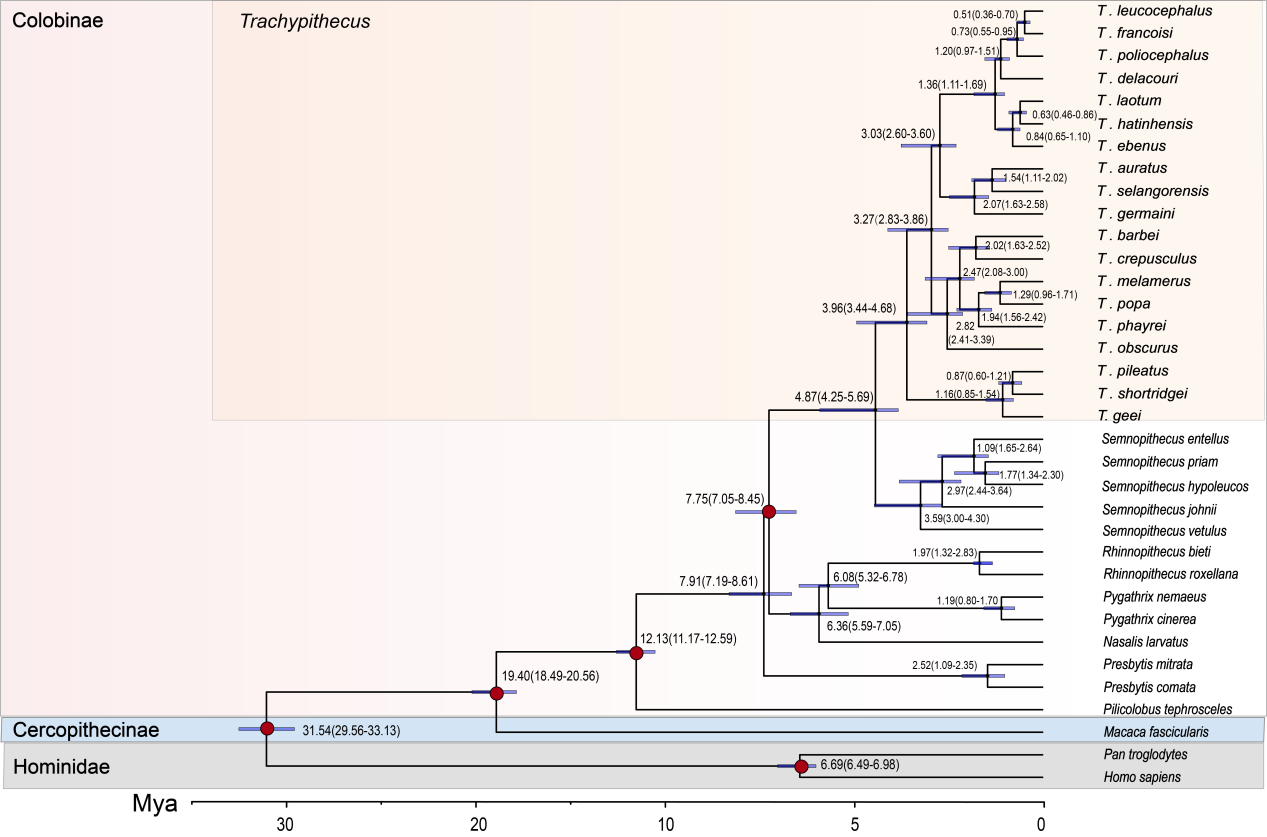

**Supplementary Figure 18 | Time-calibrated phylogenetic tree of the genus *Trachypithecus* and related primates.** The chronogram was estimated using MCMCtree based on genomic data and fossil calibrations. The numbers at each node indicate the estimated mean divergence times in millions of years ago (Mya). The blue horizontal bars represent the 95% highest posterior density (HPD) intervals for the node ages. Red circles mapped on specific nodes denote the fossil calibration points used in the analysis, which include the Hominoidea-Cercopithecoidea divergence (29.2–33.1 Mya), the Cercopithecinae-Colobinae split (18.0–20.6 Mya), the African-Asian colobines divergence (10.0–12.5 Mya), the odd-nosed monkeys-classical langurs divergence (6.5–8.5 Ma), the Hominini-*Pan* split (6.0‒7.0 Mya) Ma. The scale bar at the bottom indicates time in Mya. Background shading highlights the major taxonomic groups: Colobinae (with a specific focus on the *Trachypithecus* radiation), Cercopithecinae, and Hominidae.

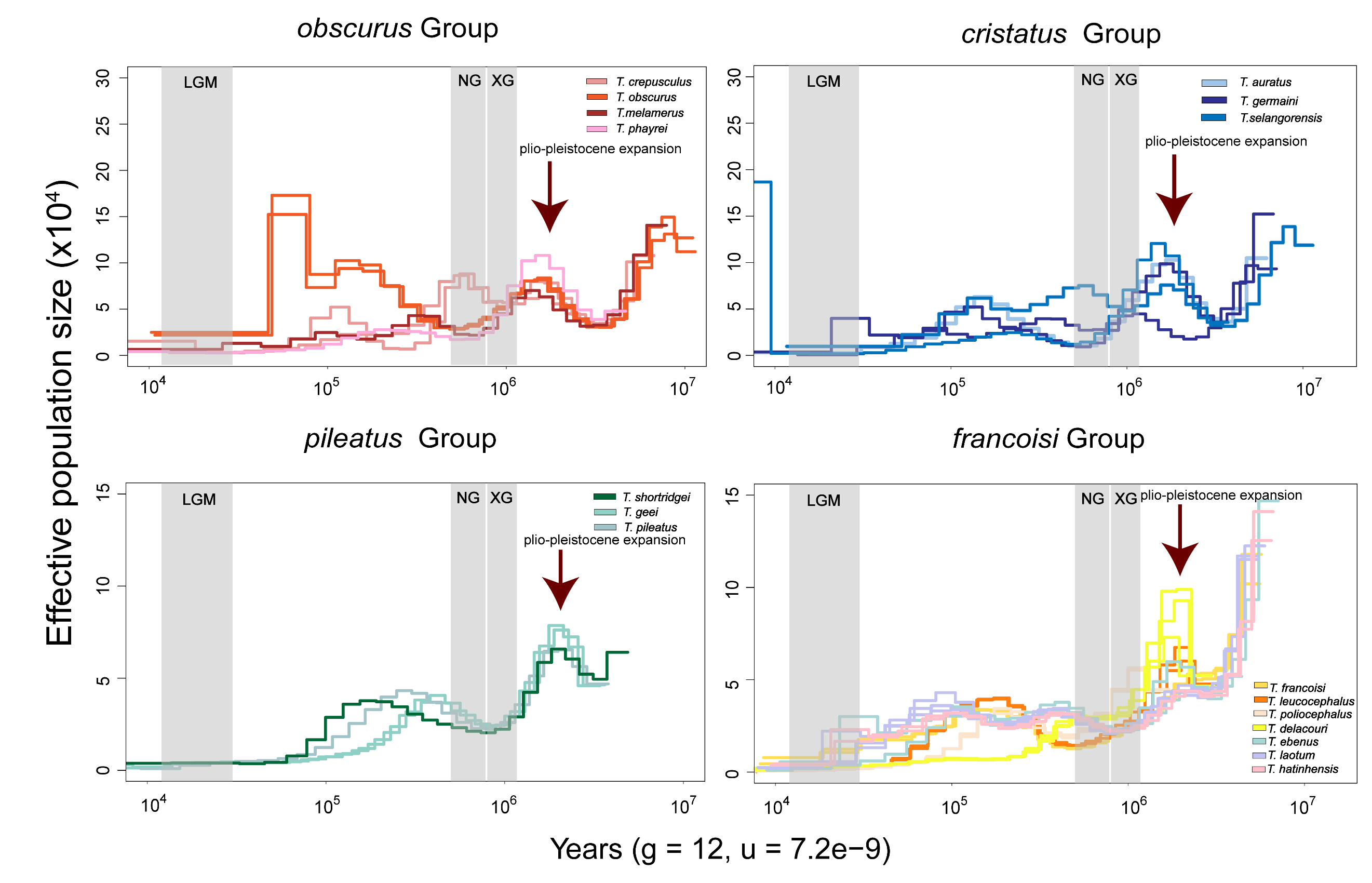

**Supplementary Figure 19 | Demographic history of the genus *Trachypithecus* inferred by PSMC. A–D** Pairwise sequentially Markovian coalescent (PSMC) models showing the fluctuations in effective population size (*Ne*) over time for the four species groups: *T. obscurus* group (**A**), *T. cristatus* group (**B**), *T. pileatus* group (**C**), and *T. francoisi* group (**D**). The x-axis represents time in years before present on a logarithmic scale, scaled with an assumed generation time (g) of 12 years and a mutation rate (μ) of 7.2e-9 per site per generation. The y-axis indicates the effective population size (x10^4^). Gray shaded vertical bars represent major Pleistocene glacial periods: LGM, Last Glacial Maximum (~15–30 thousand years ago, kya); PG, Penultimate Glaciation (~130–200 kya); NG, Naynayxungla Glaciation (~0.5–0.78 million years ago, Mya); and XG, Xixiabangma Glaciation (~0.9–1.1 Mya). Arrows indicate the *Trachypithecus* population expansion event that occurred approximately 2-3 Mya.

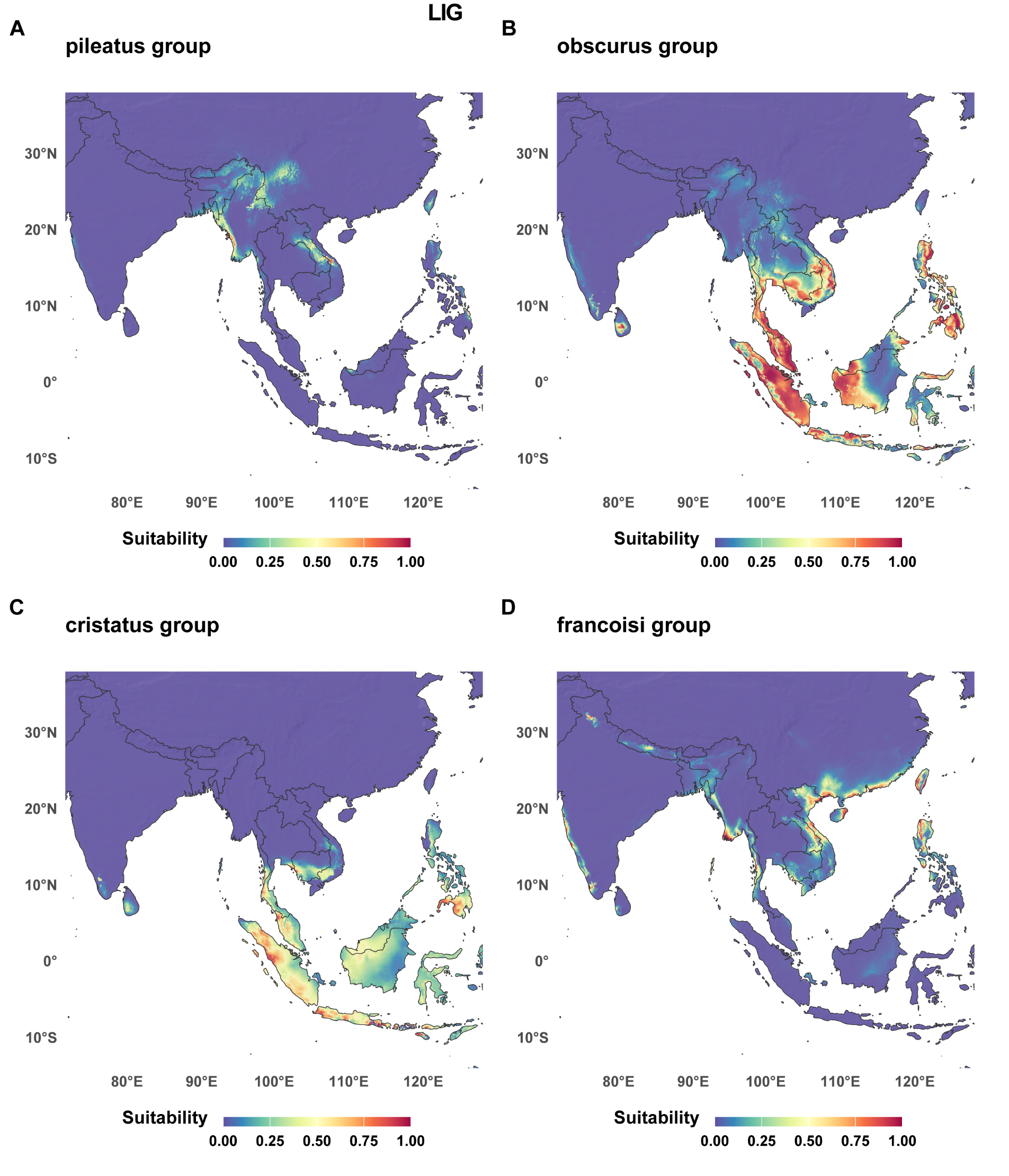

**Supplementary Figure 20 | Predicted potential habitat suitability for the four *Trachypithecus* species groups during the Last Interglacial (LIG).** Potential geographic distribution during the LIG period (~130,000 years ago) was modeled using MaxEnt based on filtered occurrence records and bioclimatic variables from WorldClim v1.4. The color gradient represents the probability of habitat suitability, ranging from red (high suitability/predicted presence) to blue (low suitability/unsuitable).

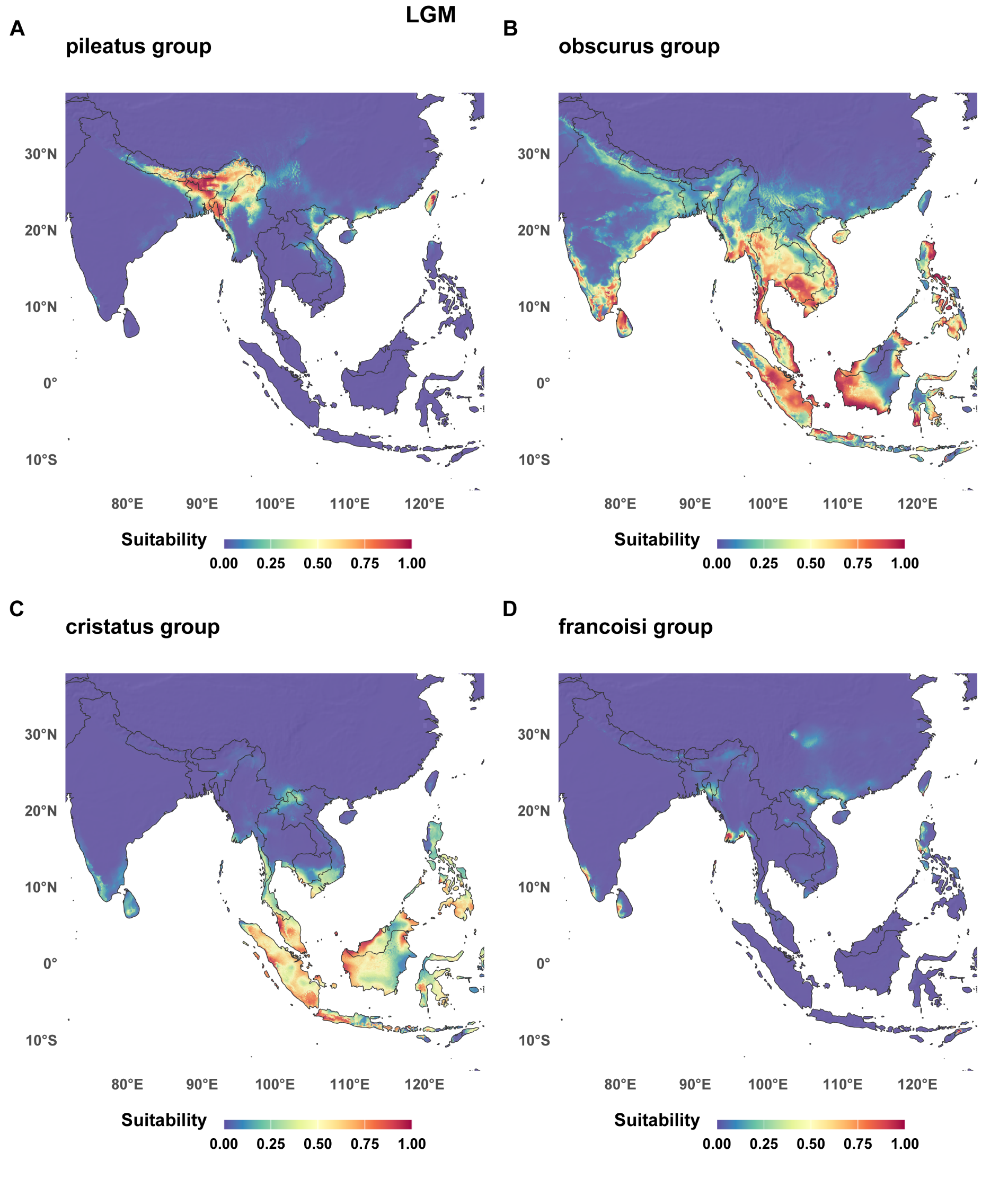

**Supplementary Figure 21 | Predicted potential habitat suitability for the four *Trachypithecus* species groups during the Last Glacial Maximum (LGM).** Potential geographic distribution during the LGM period (~22,000 years ago) was modeled using MaxEnt based on filtered occurrence records and bioclimatic variables from WorldClim v1.4. The color gradient represents the probability of habitat suitability, ranging from red (high suitability/predicted presence) to blue (low suitability/unsuitable).

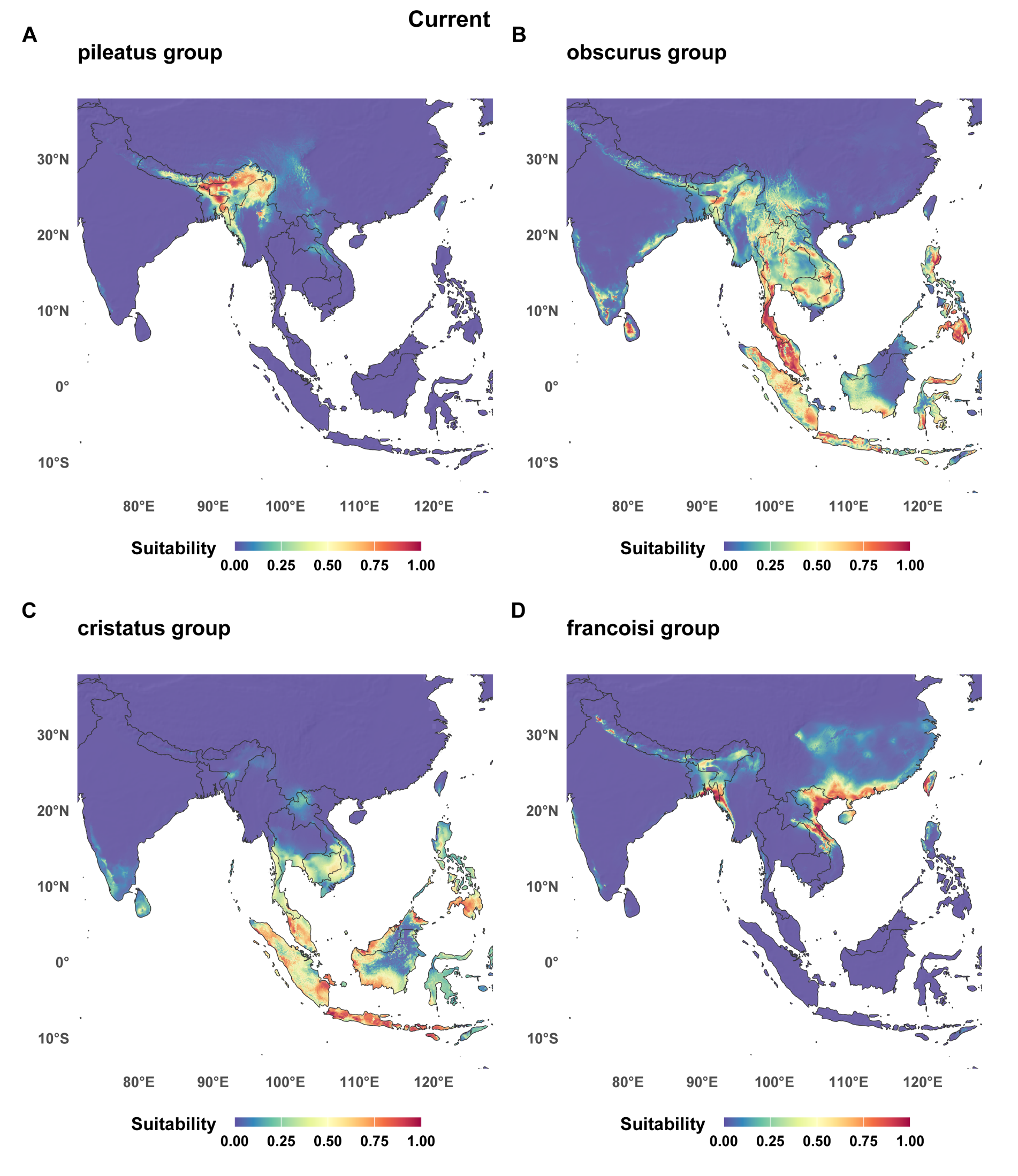

**Supplementary Figure 22 | Predicted potential habitat suitability for the four *Trachypithecus* species groups under current climatic conditions.** Potential geographic distribution under contemporary climate was modeled using MaxEnt based on filtered occurrence records and bioclimatic variables from WorldClim v1.4. The color gradient represents the probability of habitat suitability, ranging from red (high suitability/predicted presence) to blue (low suitability/unsuitable).

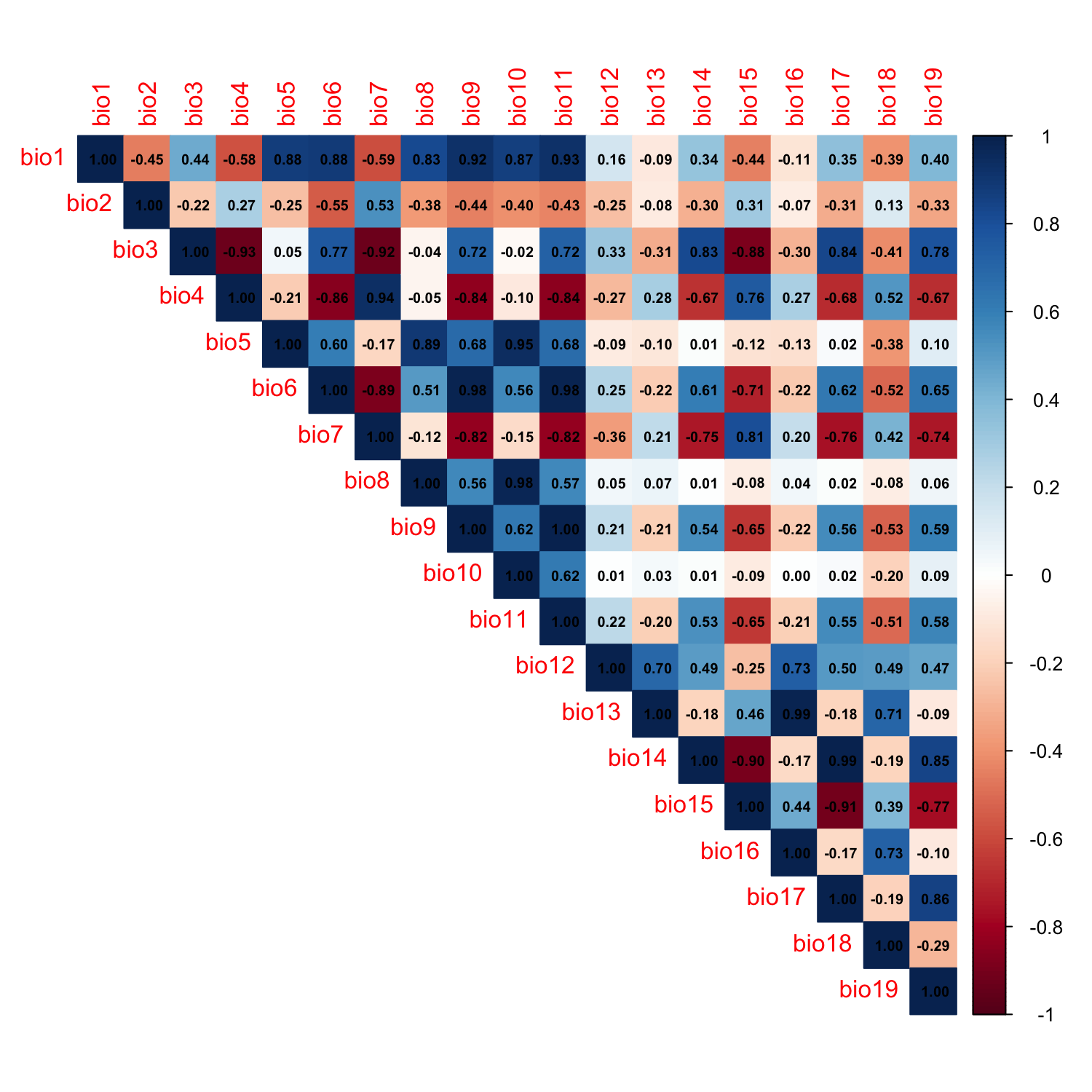

**Supplementary Figure 23 | Relative importance of bioclimatic variables in predicting habitat suitability for the four *Trachypithecus* species groups.** The bar chart illustrates the relative contribution of each retained environmental variable to the MaxEnt species distribution model. The x-axis represents the selected bioclimatic variables from WorldClim v1.4, and the y-axis indicates their percent contribution (or permutation importance) to the predictive model. Annual precipitation (Bio12) and Isothermality (Bio3) emerged as the primary environmental drivers shaping the geographic distribution of the lineages. This highlights the critical dependency and narrow physiological tolerance of the limestone-endemic primates to specific moisture regimes and temperature stability.

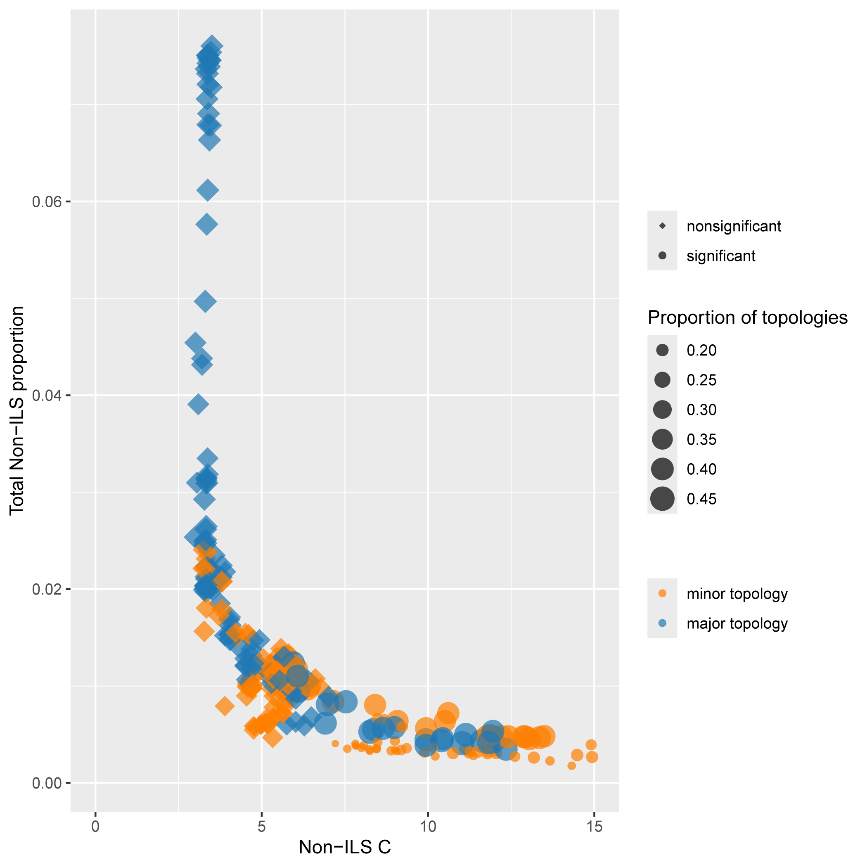

**Supplementary Figure 24 | QuIBL analysis of genome-wide introgression versus incomplete lineage sorting (ILS).** The scatter plot illustrates the relationship between internal branch lengths (Non-ILS C, in coalescent units) and the estimated fraction of introgression (Total Non-ILS proportion) across all tested triplets using QuIBL. Each point represents a triplet analysis, with shapes indicating statistical significance (circles: *P* < 0.05; diamonds: non-significant) and colors distinguishing between the major (blue) and minor (orange) alternative topologies. Point size corresponds to the relative proportion of the respective topology across the genome. The low Total Non-ILS proportions observed for the vast majority of triplets, particularly for the minor topologies, indicate that the observed gene tree heterogeneity in *Trachypithecus* is predominantly driven by pervasive ILS rather than extensive inter-species introgression.

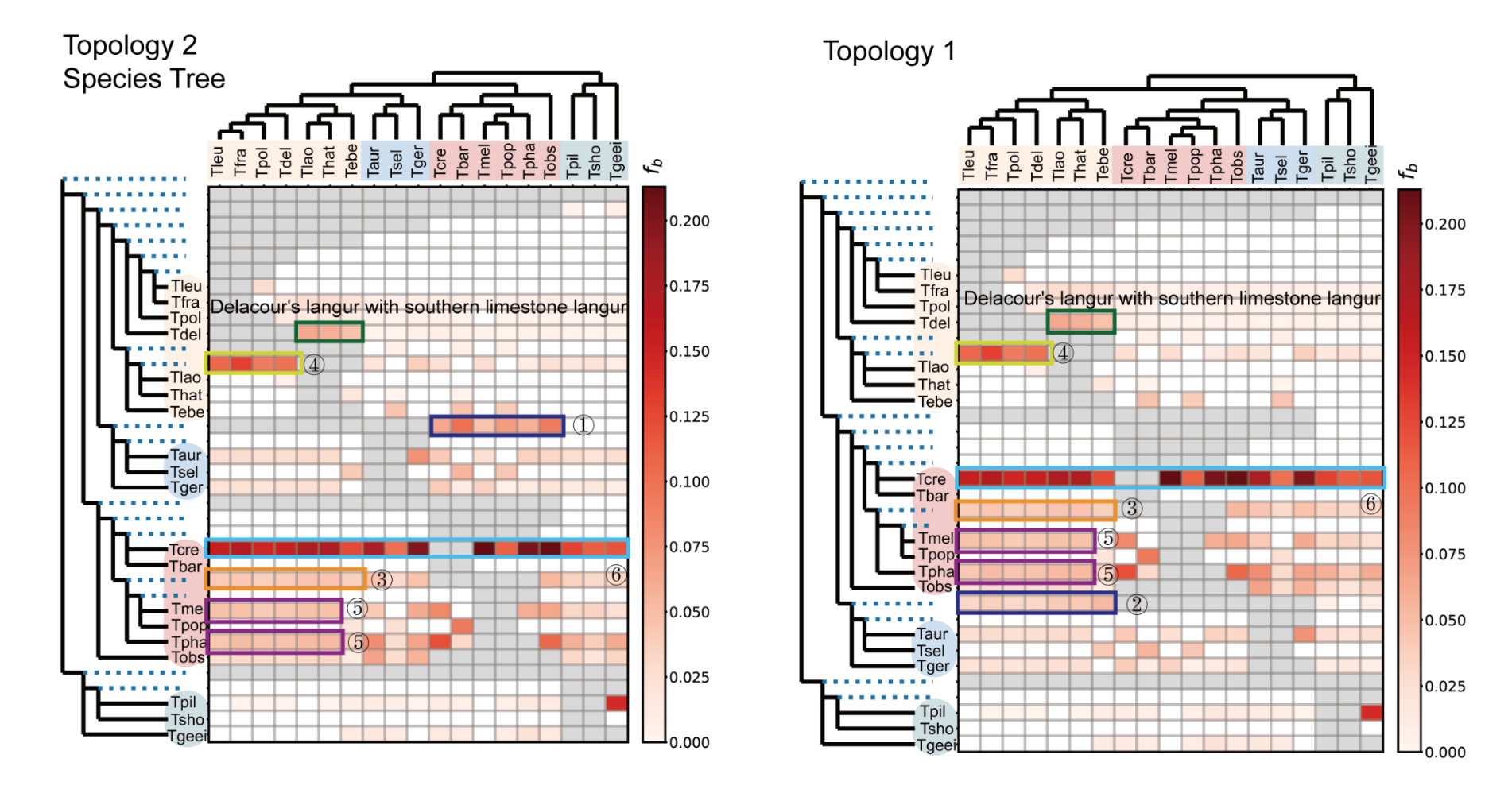

**Supplementary Figure 25 | Localization of introgression signals using *f*branch (*f*b) statistics.** Heatmaps display the *f*b values calculated within the D-suite framework to identify specific gene flow events across the *Trachypithecus* phylogeny. Results are shown for two reference topologies: the inferred species tree (Topology 2, left) and an alternative topology (Topology 1, right). The numbered symbols (1–5) indicate the five primary introgression events identified and discussed in detail in the main text. Green boxes highlight the genetic affinity between Delacour’s langur (*T. delacouri*) and the southern limestone langur group. Color intensity represents the strength of the *f*b signal, localizing signatures of excess allele sharing onto specific internal branches of the species tree.

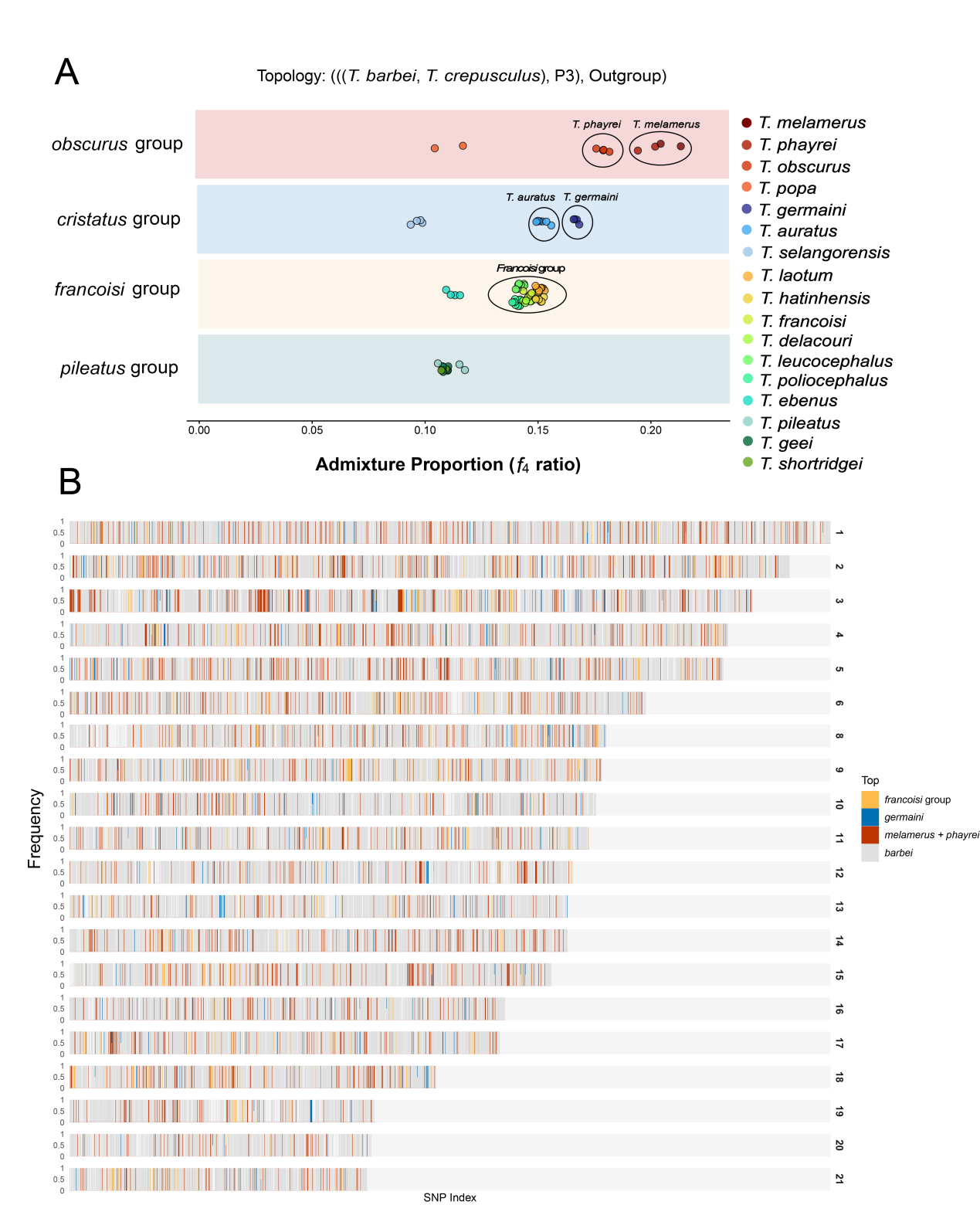

**Supplementary Figure 26 | Admixture proportions and genome-wide mosaicism of introgression in *Trachypithecus crepusculus*.** **A**, Estimation of admixture proportions using *f*4-ratio statistics. The analysis assumes the baseline phylogenetic framework (((*T. barbei*, *T. crepusculus*), P3), Outgroup), evaluating the extent of gene flow by systematically iterating P3 across four major species groups (*obscurus*, *cristatus*, *francoisi*, and *pileatus*). Colored dots represent the individual P3 taxa tested, corresponding to the right-hand legend. Black circles highlight specific lineages exhibiting prominent introgression signals, identifying them as key potential admixed species or major genetic donors in this reticulate evolutionary network. **B**, Genome-wide local ancestry and topology weighting across the autosomes. Horizontal tracks represent individual chromosomes (1–21, excluding chromosome 7, which represents the X chromosome) sorted by SNP index. Colors denote the relative frequency and genomic distribution of different phylogenetic affinities: the expected baseline sister-group relationship with *T. barbei* (grey), alongside three distinct introgressed ancestries originating from the *francoisi* group (orange), *T. germaini* (blue), and the *T. melamerus* + *T. phayrei* clade (red). The dense, alternating vertical bands visually capture the highly mosaic genomic architecture, illustrating the complex historical gene flow and differential introgression dynamics across the genome.

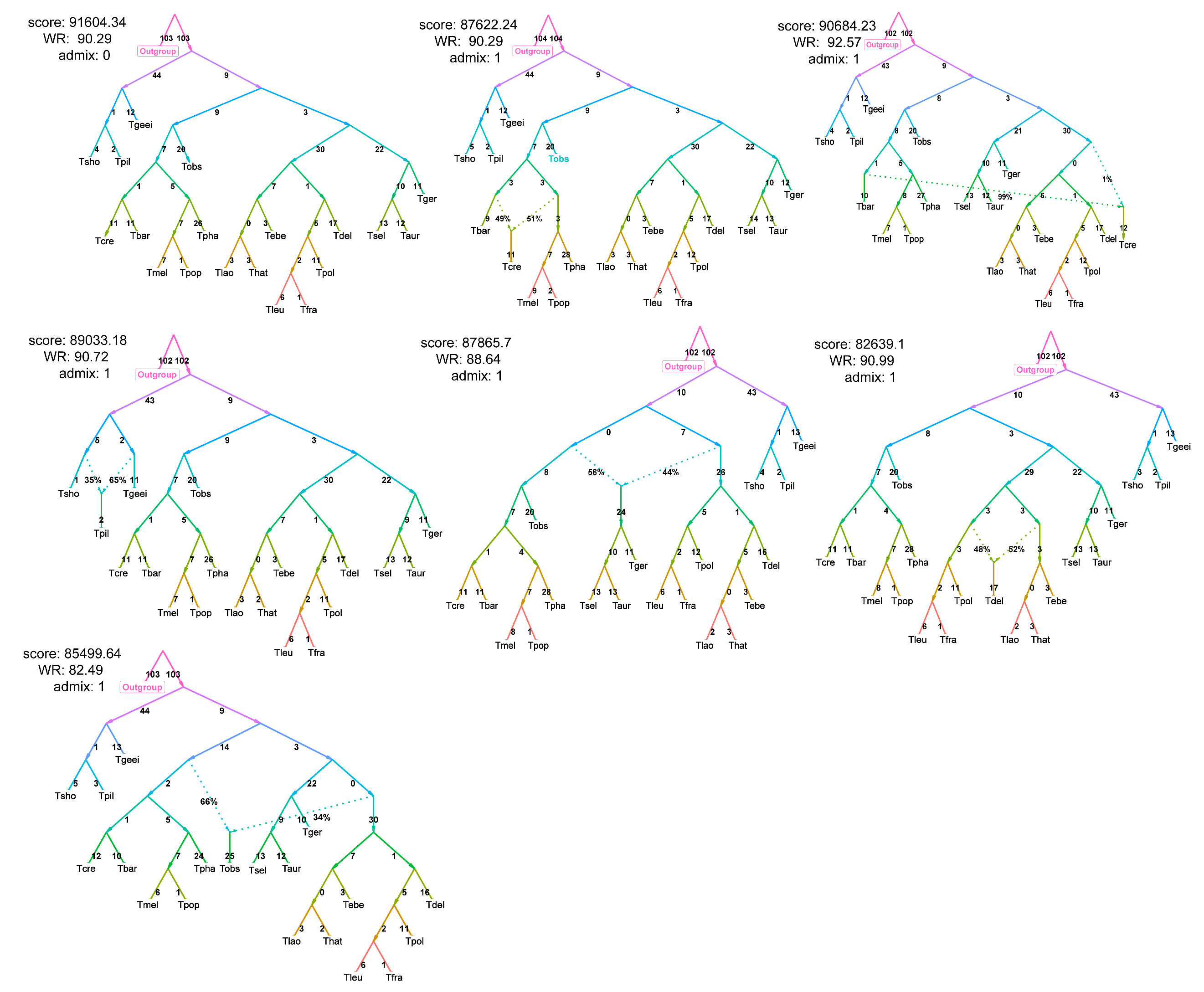

**Supplementary Figure 27 | Candidate admixture graph models (0–1 migration edges) for the genus *Trachypithecus*.** The panels display a series of candidate phylogenetic networks inferred using qpGraph based on the *f*2 matrix. The models represent increasing levels of evolutionary complexity, incorporating 0-1 migration edges (admixture events). For each graph, the log-likelihood score and the worst residual (WR) value are provided in the upper-left corner; a lower WR indicates a better fit between the modeled and observed genetic distances. Dashed lines represent divergent lineages, while numbers indicate inferred admixture events with their estimated ancestry proportions (%).

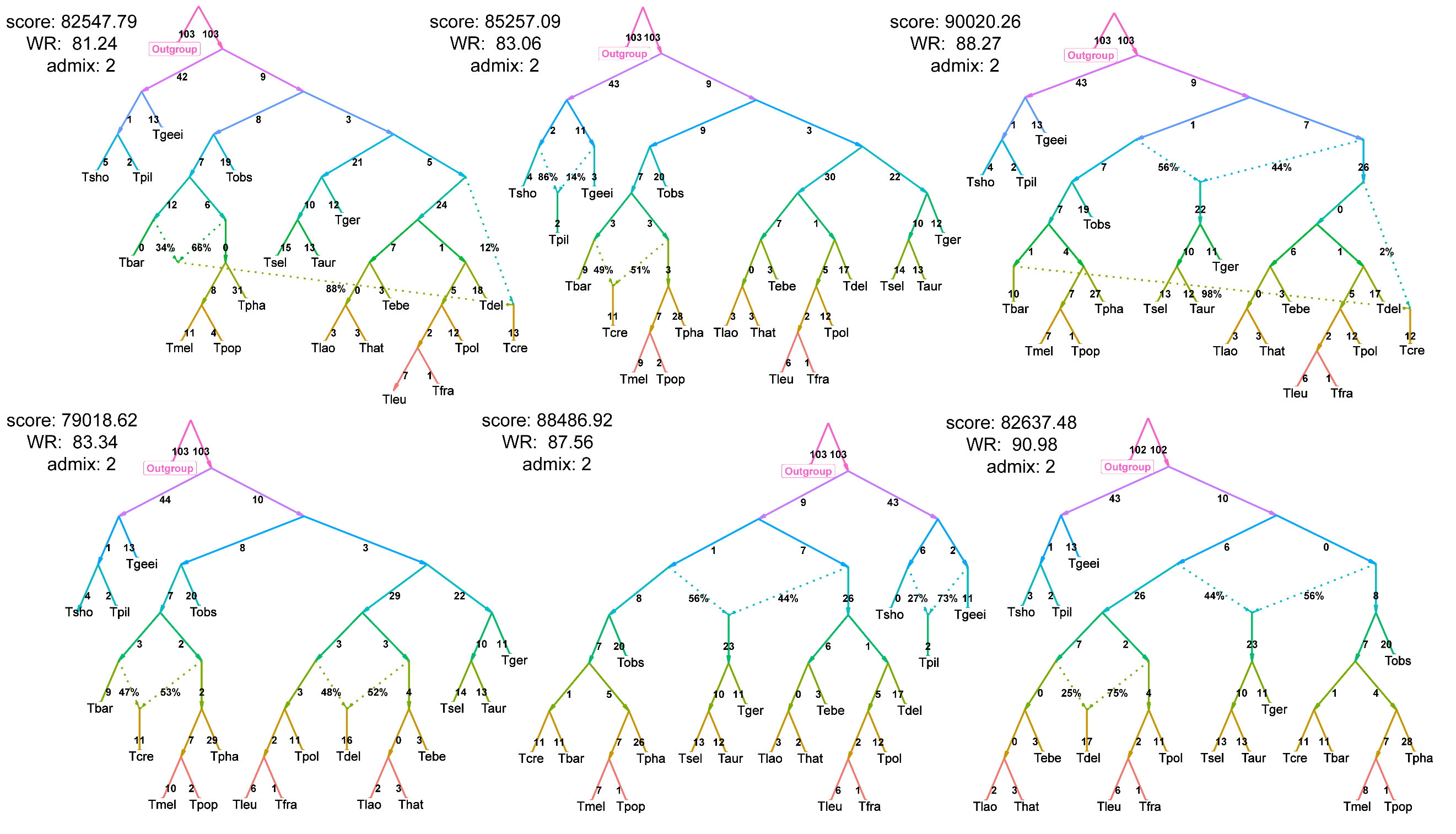

**Supplementary Figure 28 | Candidate admixture graph models (2 migration edges) and identifying the best-fitting phylogenetic network.** The panels display a series of candidate phylogenetic networks inferred using qpGraph based on the *f*2 matrix, representing high-complexity scenarios with 2 migration edges (admixture events). For each graph, the log-likelihood score and the worst residual (WR) value are provided in the upper-left corner; a lower WR indicates a better fit between the modeled and observed genetic distances. Dashed lines represent divergent lineages, while numbers indicate inferred admixture events with their estimated ancestry proportions (%).

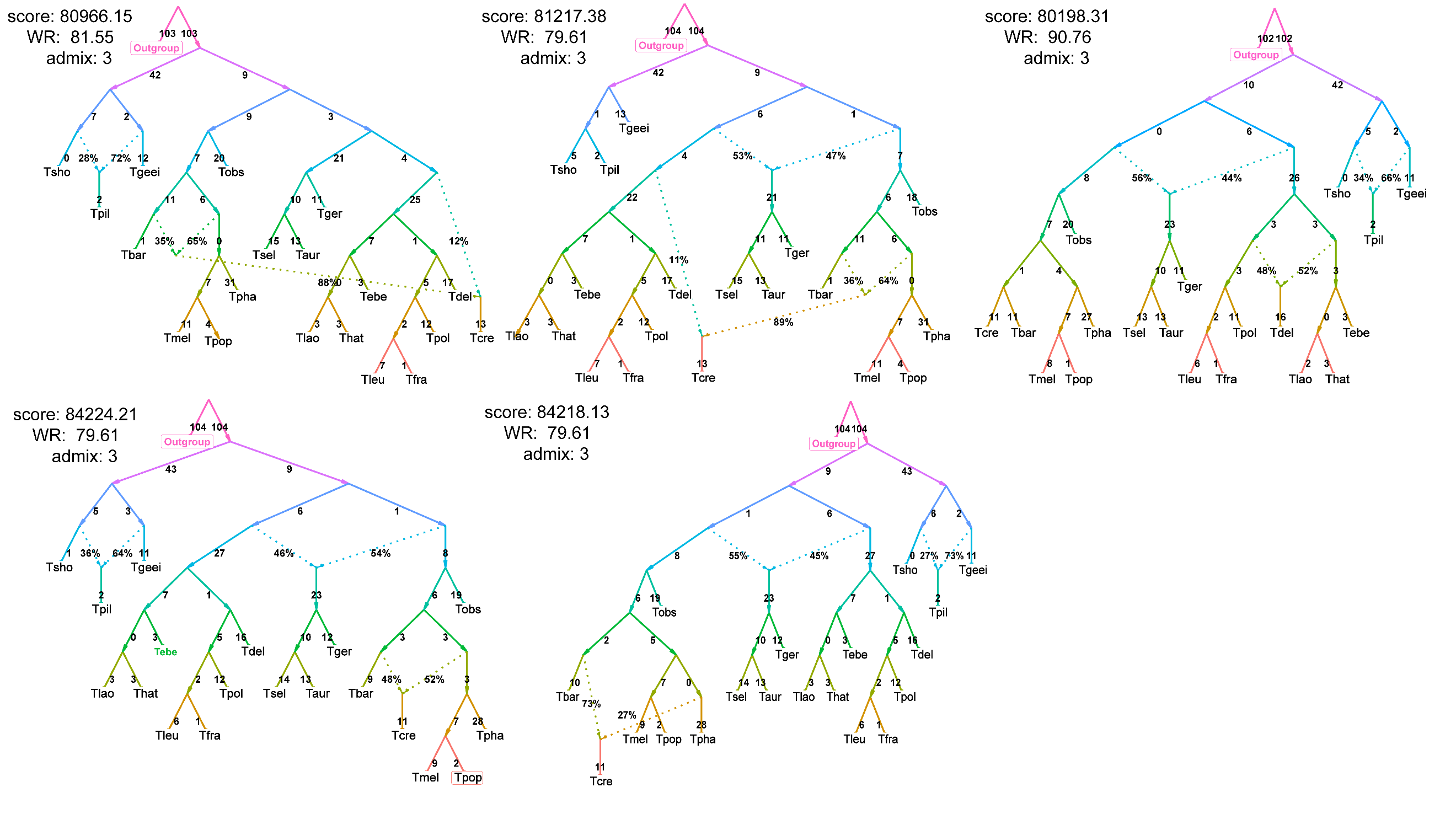

**Supplementary Figure 29 | Candidate admixture graph models (3 migration edges) and identifying the best-fitting phylogenetic network.** The panels display a series of candidate phylogenetic networks inferred using qpGraph based on the *f*2 matrix, representing high-complexity scenarios with 3 migration edges (admixture events). For each graph, the log-likelihood score and the worst residual (WR) value are provided in the upper-left corner; a lower WR indicates a better fit between the modeled and observed genetic distances. Dashed lines represent divergent lineages, while numbers indicate inferred admixture events with their estimated ancestry proportions (%).

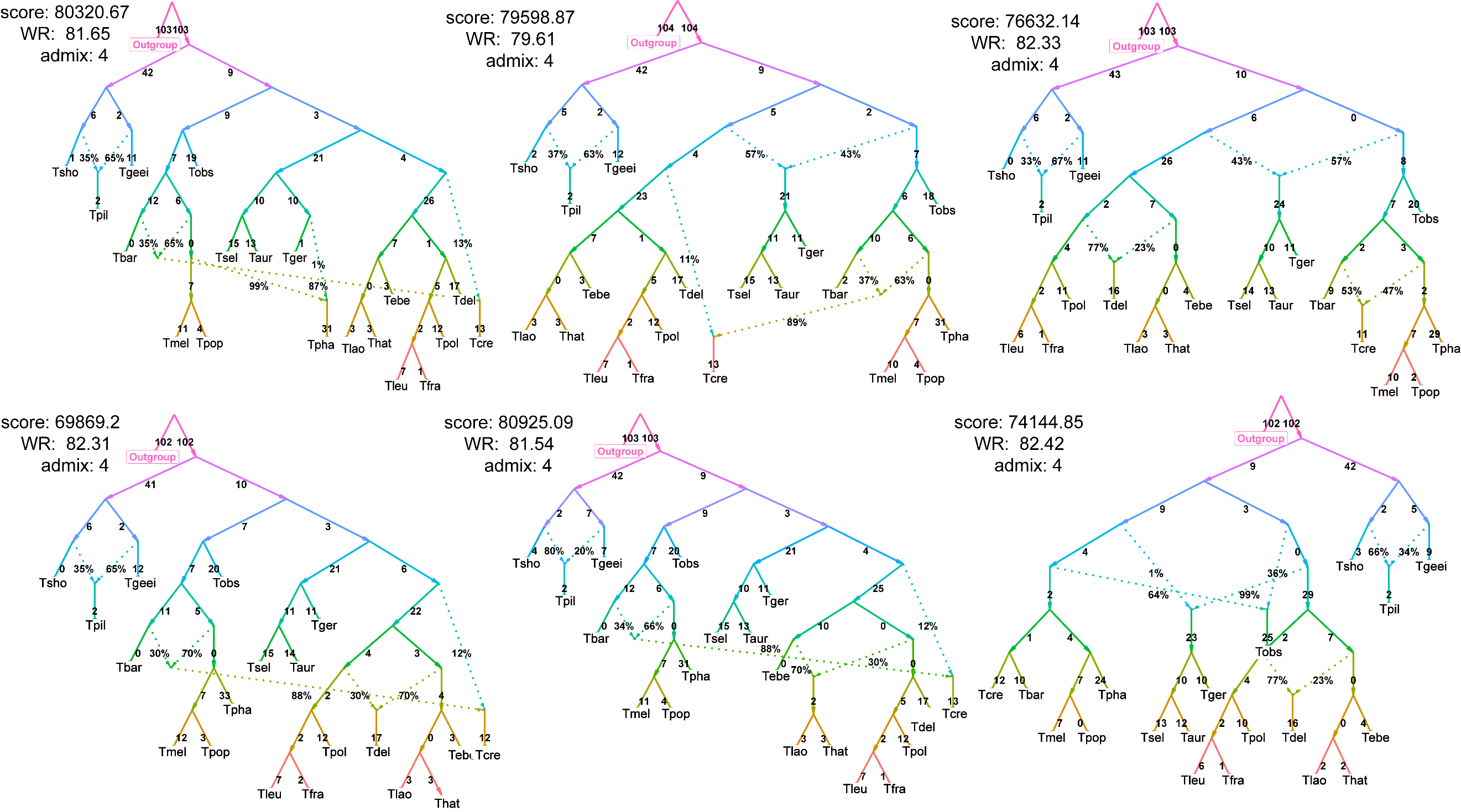

**Supplementary Figure 30 | Candidate admixture graph models (4 migration edges) and identifying the best-fitting phylogenetic network.** The panels display a series of candidate phylogenetic networks inferred using qpGraph based on the *f*2 matrix, representing high-complexity scenarios with 4 migration edges (admixture events). For each graph, the log-likelihood score and the worst residual (WR) value are provided in the upper-left corner; a lower WR indicates a better fit between the modeled and observed genetic distances. Dashed lines represent divergent lineages, while numbers indicate inferred admixture events with their estimated ancestry proportions (%).

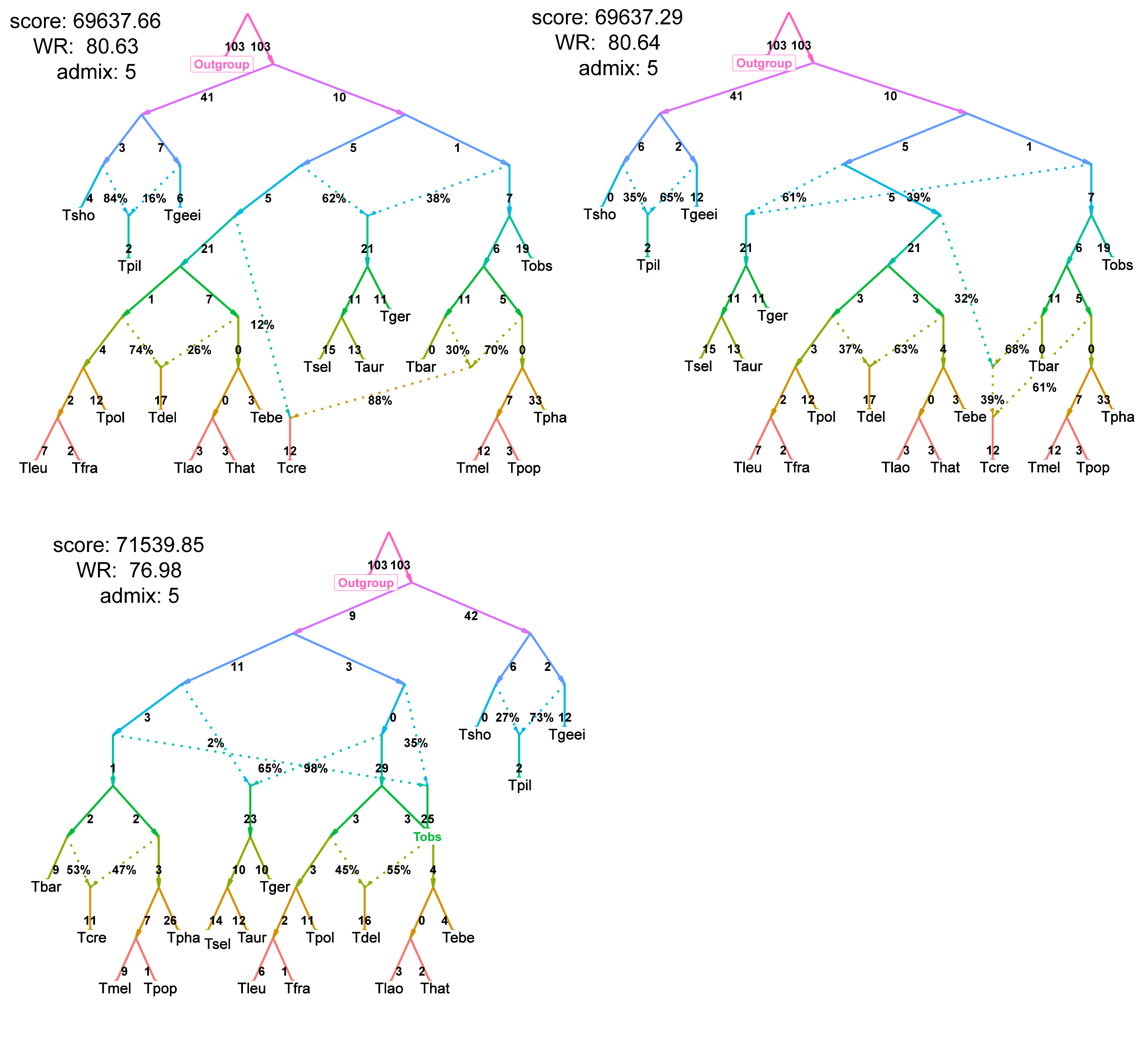

**Supplementary Figure 31 | Candidate admixture graph models (5 migration edges) and identifying the best-fitting phylogenetic network.** The panels display a series of candidate phylogenetic networks inferred using qpGraph based on the *f*2 matrix, representing high-complexity scenarios with 5 migration edges (admixture events). For each graph, the log-likelihood score and the worst residual (WR) value are provided in the upper-left corner; a lower WR indicates a better fit between the modeled and observed genetic distances. Dashed lines represent divergent lineages, while numbers indicate inferred admixture events with their estimated ancestry proportions (%).

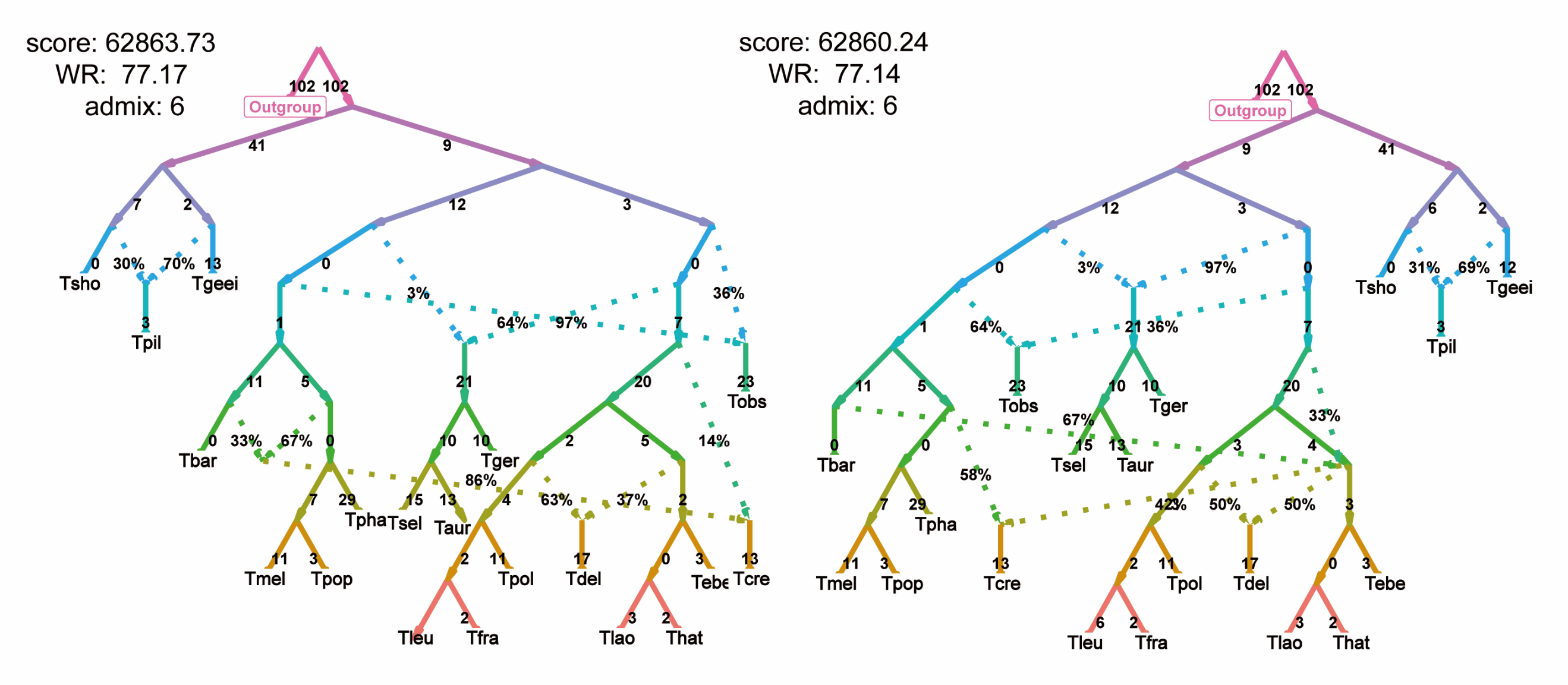

**Supplementary Figure 32 | Candidate admixture graph models (6 migration edges) and identifying the best-fitting phylogenetic network.** The panels display a series of candidate phylogenetic networks inferred using qpGraph based on the *f*2 matrix, representing high-complexity scenarios with 6 migration edges (admixture events). For each graph, the log-likelihood score and the worst residual (WR) value are provided in the upper-left corner; a lower WR indicates a better fit between the modeled and observed genetic distances. Dashed lines represent divergent lineages, while numbers indicate inferred admixture events with their estimated ancestry proportions (%).

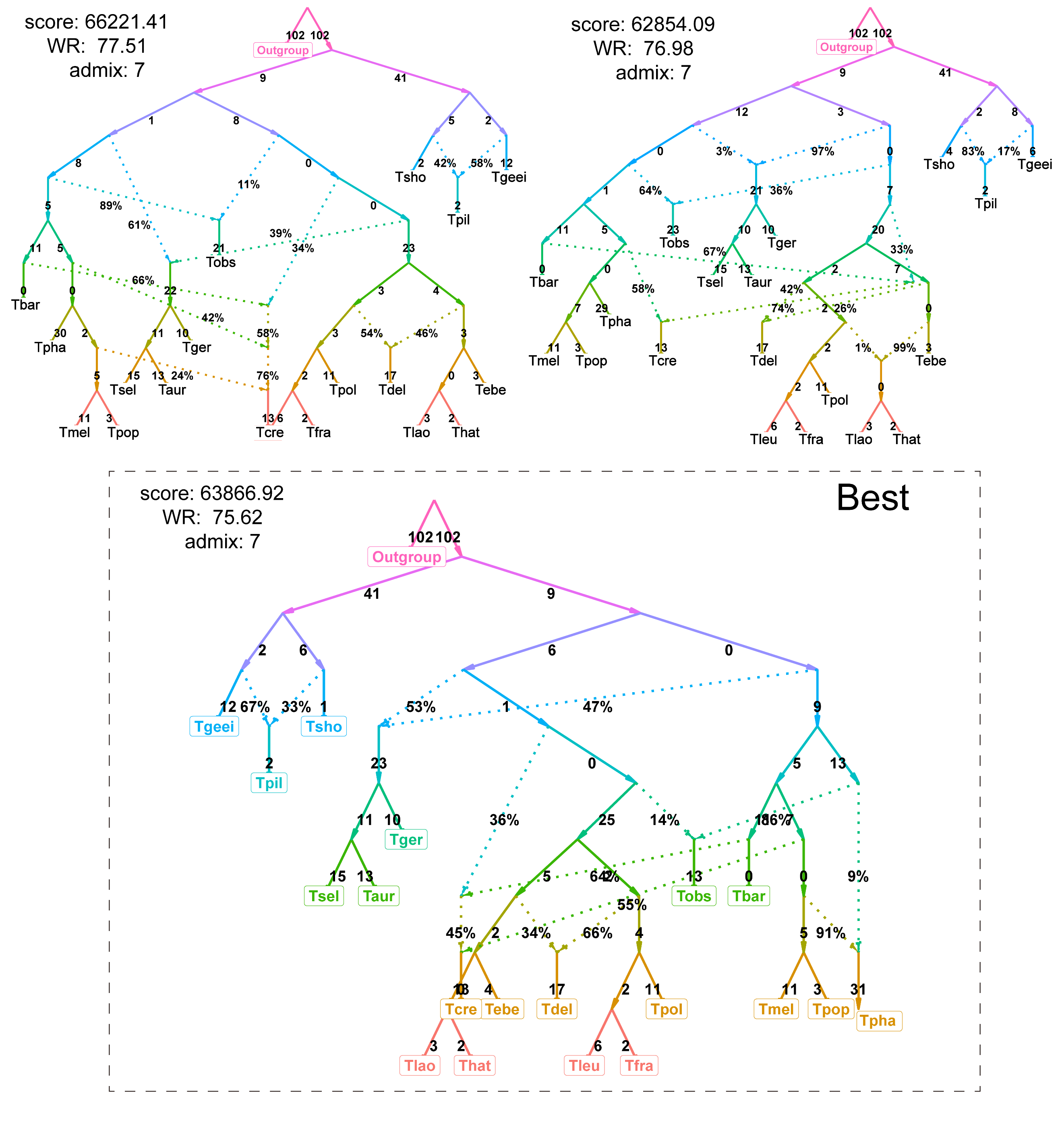

**Supplementary Figure 33 | Candidate admixture graph models (7 migration edges) and identifying the best-fitting phylogenetic network.** The panels display a series of candidate phylogenetic networks inferred using qpGraph based on the *f*2 matrix, representing high-complexity scenarios with 7 migration edges (admixture events). For each graph, the log-likelihood score and the worst residual (WR) value are provided in the upper-left corner; a lower WR indicates a better fit between the modeled and observed genetic distances. The model enclosed in the dashed box represents the best-fitting model across all tested scenarios, as indicated by the globally lowest WR value (WR = 75.62). Dashed lines represent divergent lineages, while numbers indicate inferred admixture events with their estimated ancestry proportions (%).

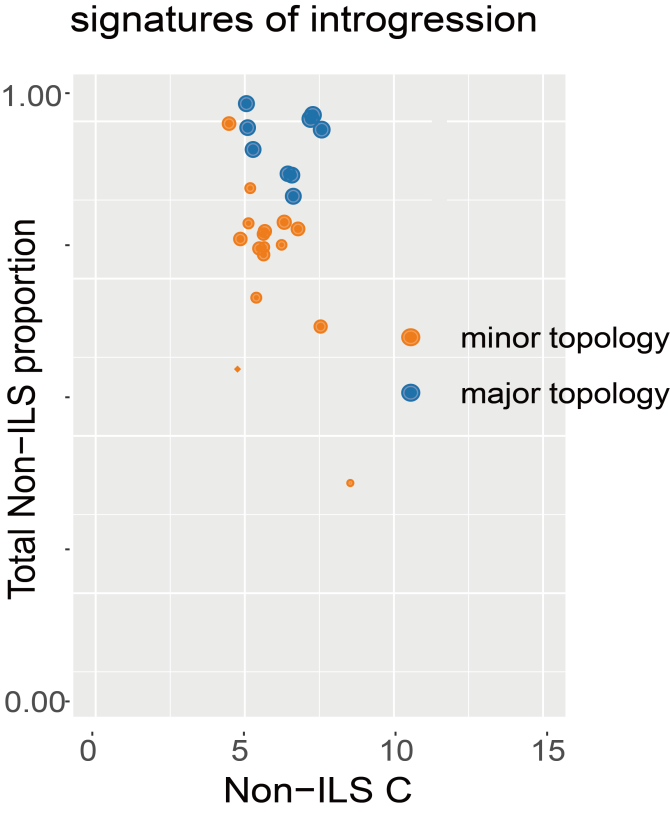

**Supplementary Figure 34 | QuIBL analysis of genome-wide introgression versus incomplete lineage sorting (ILS) within the *francoisi* species group.** The scatter plot illustrates the relationship between internal branch lengths (Non-ILS C, in coalescent units) and the estimated fraction of introgression (Total Non-ILS proportion) across all tested triplets using QuIBL. Each point represents a triplet analysis, with shapes indicating statistical significance (circles: *P* < 0.05; diamonds: non-significant) and colors distinguishing between the major (blue) and minor (orange) alternative topologies. Point size corresponds to the relative proportion of the respective topology across the genome. Notably, all analyzed triplets across the topologies consistently support a certain proportion of introgression, demonstrating that gene tree heterogeneity within the *francoisi* species group is driven by non-negligible gene flow alongside ILS.

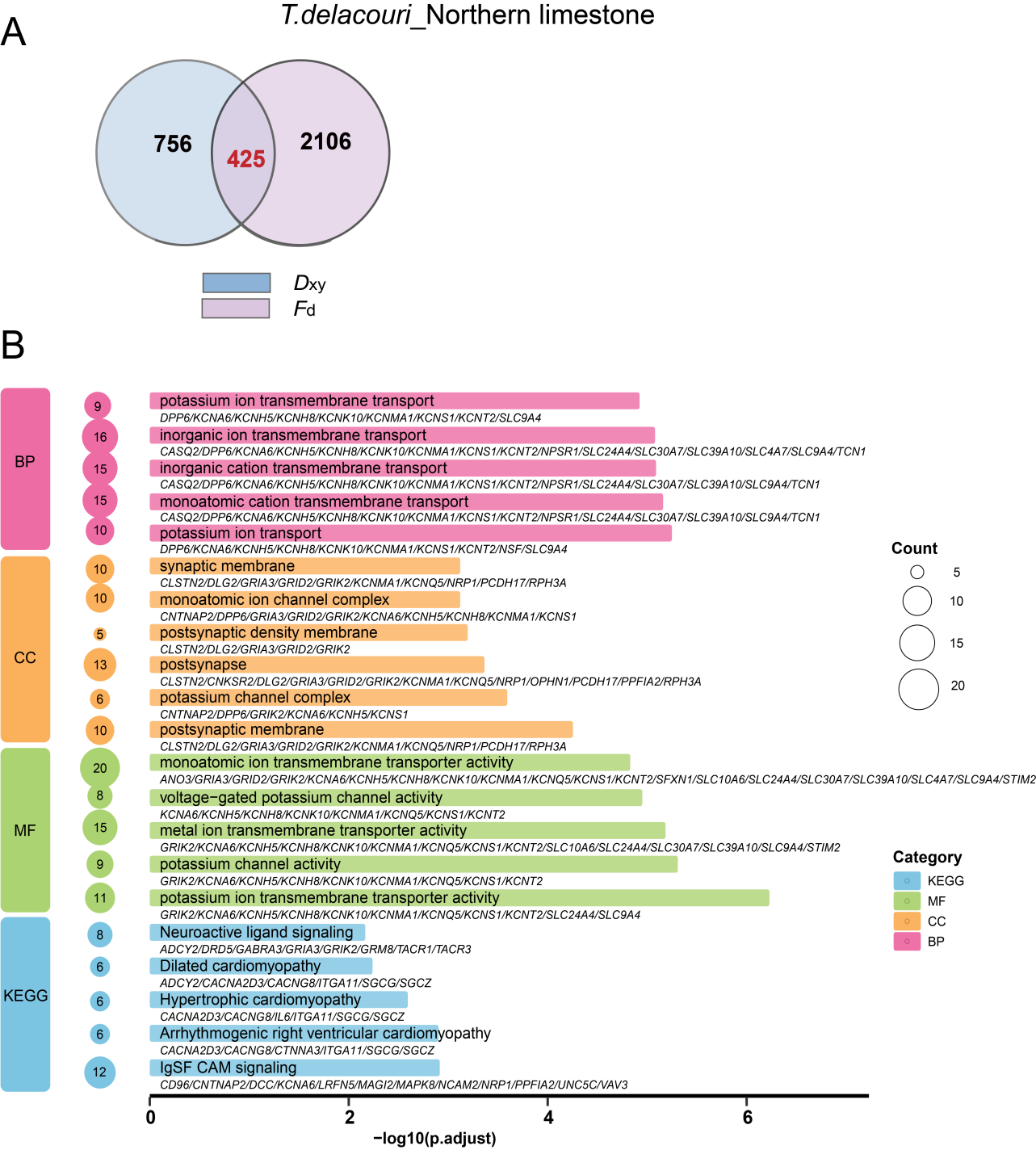

**Supplementary Figure 35 | Genomic signatures and functional enrichment of introgression from northern limestone langurs into *T. delacouri*.** **A**, Venn diagram illustrating the intersection of introgressed candidate genes identified by the top 3% outlier windows of the *F*_d_ (pink) and *D*_xy_ (blue) statistics. A total of 425 genes show consistent signals of introgression across both metrics. **B**, Gene Ontology (GO) and KEGG pathway enrichment analysis for the 425 overlapping genes. The bar plot displays the top five significantly enriched terms across Biological Process (BP, pink), Cellular Component (CC, orange), Molecular Function (MF, green), and KEGG pathways (blue). The x-axis indicates the statistical significance (-log10 *P*-value), while the circle size corresponds to the number of introgressed genes mapping to each specific term or pathway. The specific candidate genes associated with each enriched category are listed below to the corresponding bars.

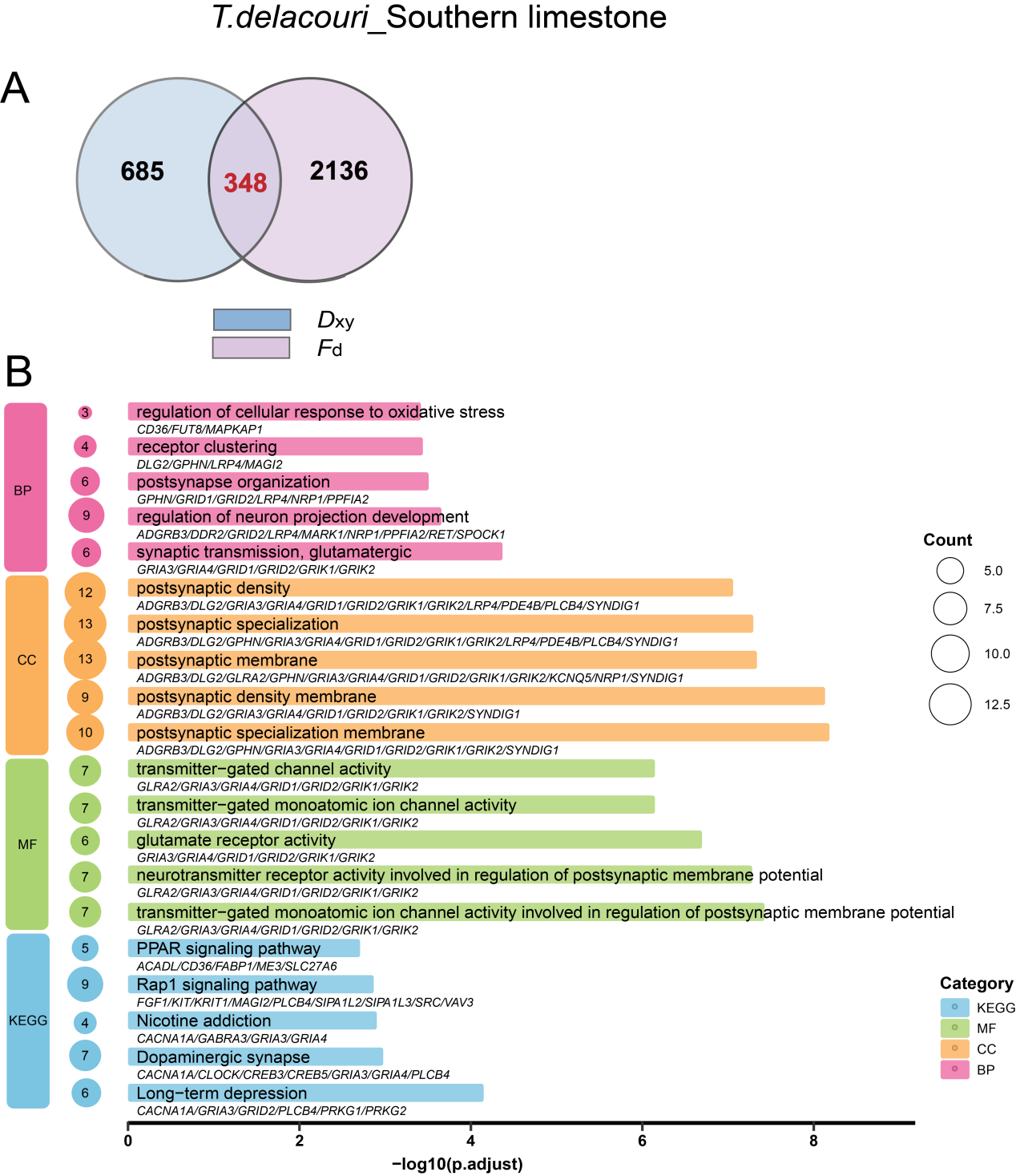

**Supplementary Figure 36 | Genomic signatures and functional enrichment of introgression from southern limestone langurs into *T. delacouri*.** **A**, Venn diagram illustrating the intersection of introgressed candidate genes identified by the top 3% outlier windows of the *F*_d_ (pink) and *D*_xy_ (blue) statistics. A total of 348 genes show consistent signals of introgression across both metrics. **B**, Gene Ontology (GO) and KEGG pathway enrichment analysis for the 348 overlapping genes. The bar plot displays the top five significantly enriched terms across Biological Process (BP, pink), Cellular Component (CC, orange), Molecular Function (MF, green), and KEGG pathways (blue). The x-axis indicates the statistical significance (-log10 *P*-value), while the circle size corresponds to the number of introgressed genes mapping to each specific term or pathway. The specific candidate genes associated with each enriched category are listed below to the corresponding bars.

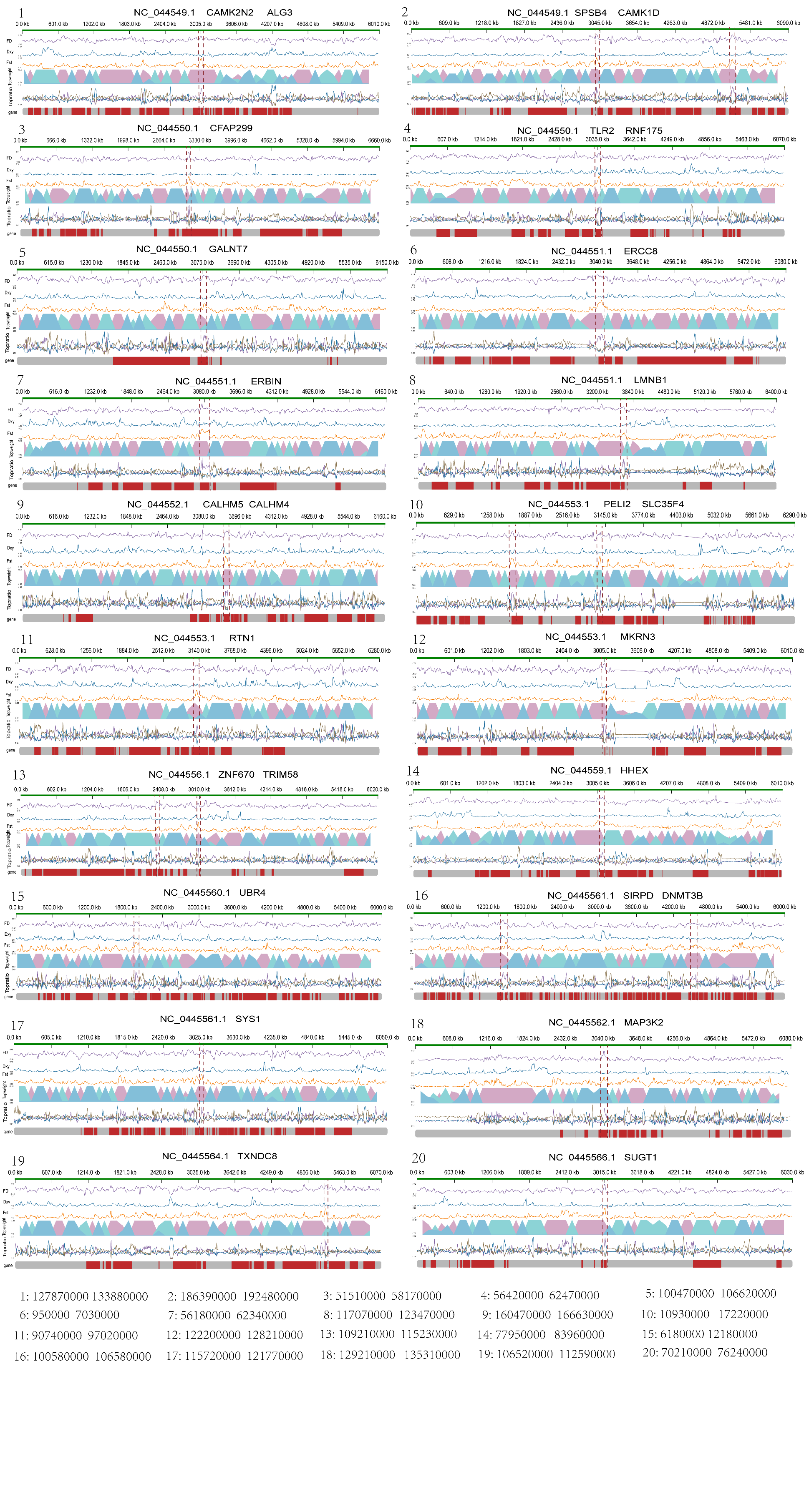

**Supplementary Figure 37 | Local genomic landscape of core introgression from northern limestone langurs into *T. delacouri*.** Each panel represents a specific genomic region identified as potentially inherited from northern limestone langurs. The scaffold ID and associated candidate genes are labeled at the top of each subplot. From top to bottom, the tracks represent: (1) *F*_d_ statistic, indicating the fraction of the genome shared via introgression; (2) *D*_xy_ statistic, representing absolute sequence divergence; (3) *Fst* statistic, showing population differentiation between *T. delacouri* and southern limestone langurs; (4) Twisst results, depicting local topology weighting across the genomic interval; (5) ERICA inference, showing local ancestry and origin classification based on a Convolutional Neural Network (CNN) framework; (6) Gene annotation, where red blocks indicate protein-coding regions. Dashed red vertical lines highlight the specific genomic segments with significant signals of introgression. Figures were generated using the Rectchr package.

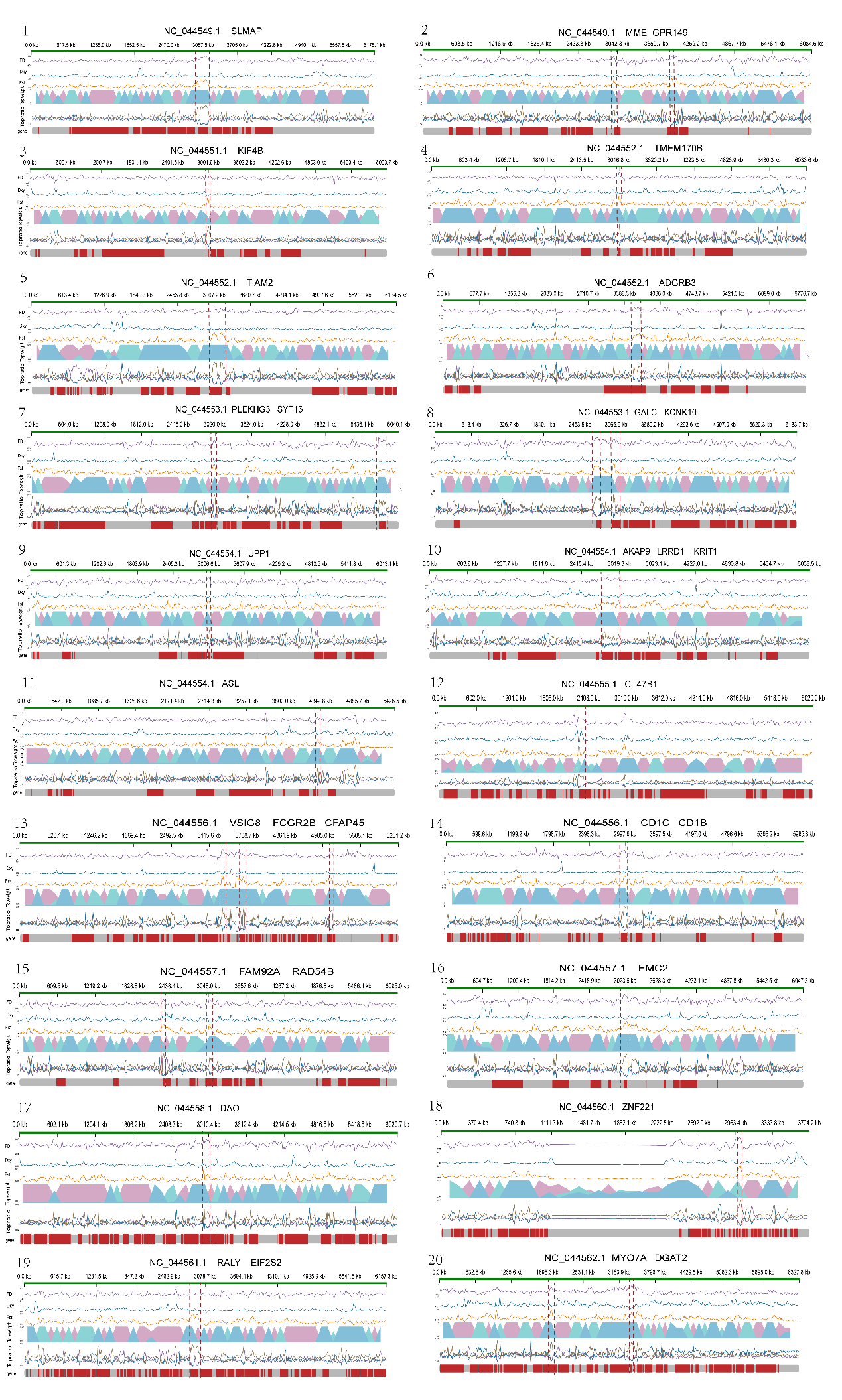

**Supplementary Figure 38 | Local genomic landscape of core introgression from southern limestone langurs into *T. delacouri*.** Each panel represents a specific genomic region identified as potentially inherited from southern limestone langurs. The scaffold ID and associated candidate genes are labeled at the top of each subplot. From top to bottom, the tracks represent: (1) *F*_d_ statistic, indicating the fraction of the genome shared via introgression; (2) *D*_xy_ statistic, representing absolute sequence divergence; (3) Fst statistic, showing population differentiation between *T. delacouri* and northern limestone langurs; (4) Twisst results, depicting local topology weighting across the genomic interval; (5) ERICA inference, showing local ancestry and origin classification based on a Convolutional Neural Network (CNN) framework; (6) Gene annotation, where red blocks indicate protein-coding regions. Dashed red vertical lines highlight the specific genomic segments with significant signals of introgression. Figures were generated using the Rectchr package.

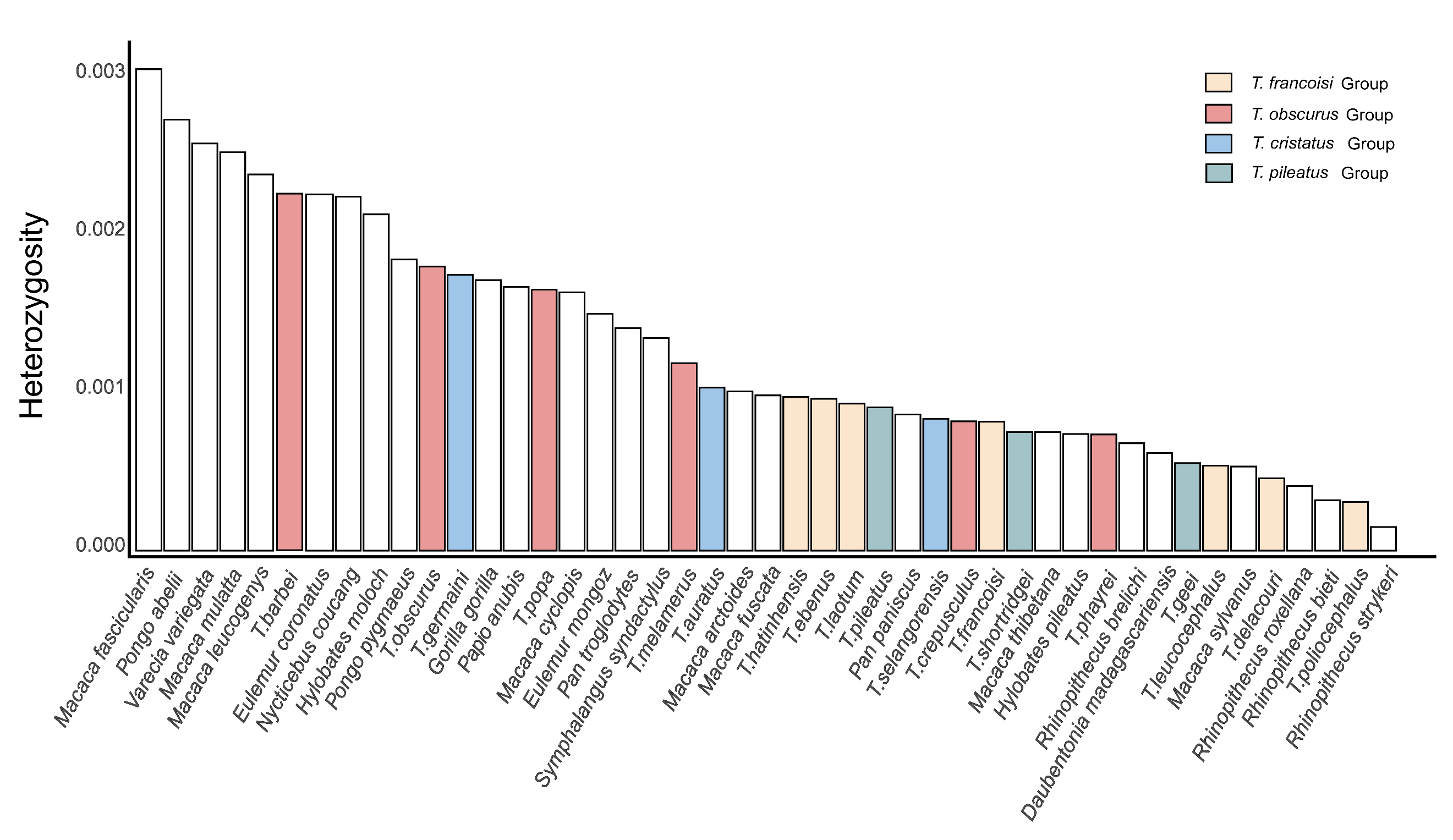

**Supplementary Figure 39 | Comparison of genome-wide heterozygosity between *Trachypithecus* and other primate species.** Bar plot displaying the estimated genome-wide heterozygosity (y-axis) across diverse primate taxa (x-axis). Species are ranked in descending order based on their heterozygosity values. The bars for *Trachypithecus* species are color-coded to denote their species groups (*francoisi*, *obscurus*, *cristatus*, and *pileatus* groups, as defined in the legend). The white bars represent other comparative non-*Trachypithecus* primate species. The plot illustrates that the genus *Trachypithecus* collectively exhibits notably low genetic diversity, approaching the lowest value currently recorded among non-human primates (*Rhinopithecus strykeri*).

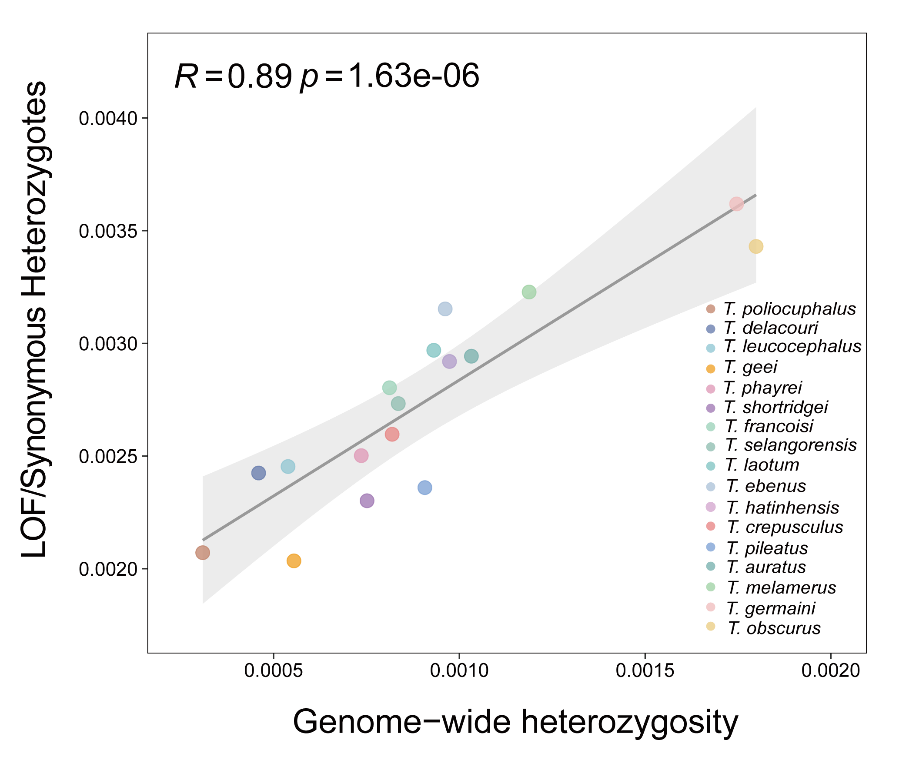

**Supplementary Figure 40 | Positive correlation between genome-wide heterozygosity and masked genetic load across *Trachypithecus* species.** Pearson correlation analysis between genome-wide heterozygosity (x-axis) and the ratio of heterozygous loss-of-function (LoF) to heterozygous synonymous variants (y-axis), which serves as a proxy for the masked genetic load. Each point represents an individual species, color-coded according to the legend. The solid grey line indicates the linear regression fit, and the shaded grey area represents the 95% confidence interval.
